# Disentangling heterogeneity of Malignant Pleural Mesothelioma through deep integrative omics analyses

**DOI:** 10.1101/2021.09.27.461908

**Authors:** Lise Mangiante, Nicolas Alcala, Alex Di Genova, Alexandra Sexton-Oates, Abel Gonzalez-Perez, Azhar Khandekar, Erik N. Bergstrom, Jaehee Kim, Colin Giacobi, Nolwenn Le Stang, Sandrine Boyault, Cyrille Cuenin, Severine Tabone-Eglinger, Francesca Damiola, Catherine Voegele, Maude Ardin, Marie-Cecile Michallet, Lorraine Soudade, Tiffany M. Delhomme, Arnaud Poret, Marie Brevet, Marie-Christine Copin, Sophie Giusiano-Courcambeck, Diane Damotte, Cecile Girard, Veronique Hofman, Paul Hofman, Jérôme Mouroux, Stephanie Lacomme, Julien Mazieres, Vincent Thomas de Montpreville, Corinne Perrin, Gaetane Planchard, Isabelle Rouquette, Christine Sagan, Arnaud Scherpereel, Francoise Thivolet, Jean-Michel Vignaud, Didier Jean, Anabelle Gilg Soit Ilg, Robert Olaso, Vincent Meyer, Anne Boland, Jean-Francois Deleuze, Janine Altmuller, Peter Nuernberg, Sylvie Lantuejoul, Akram Ghantous, Charles Maussion, Pierre Courtiol, Hector Hernandez-Vargas, Christophe Caux, Nicolas Girard, Nuria Lopez-Bigas, Ludmil B. Alexandrov, Françoise Galateau Salle, Matthieu Foll, Lynnette Fernandez-Cuesta

**Affiliations:** Rare Cancers Genomics Team (RCG), Genomic Epidemiology Branch (GEM), International Agency for Research on Cancer/World Health Organisation (IARC/WHO), Lyon, 69008, France; Institute for Research in Biomedicine (IRB Barcelona), The Barcelona Institute of Science and Technology, Baldiri Reixac, 10, 08028 Barcelona, Spain; Department of Cellular and Molecular Medicine, Department of Bioengineering and Moores Cancer Center, UC San Diego, La Jolla, CA, 92093, USA; Department of Biology, Stanford University, Stanford, USA; Department of Computational Biology, Cornell University, Ithaca, New York, USA; UMR INSERM 1052, CNRS 5286, Cancer Research Center of Lyon, MESOPATH-MESOBANK, Department of Biopathology, Cancer Centre Léon Bérard, 69008 Lyon, France; Cancer Genomic Platform, Translational Research and Innovation department, Centre Léon Bérard; Lyon, 69008; France; EpiGenomics and Mechanisms Branch (EGM); International Agency for Research on Cancer/World Health Organisation (IARC/WHO), Lyon, 69008, France; TERI (Tumor Escape, Resistance and Immunity) Department, Centre de Recherche en Cancérologie de Lyon (CRCL), Centre Léon Bérard (CLB), Université de Lyon, Université Claude Bernard Lyon 1, INSERM 1052, CNRS 5286, 69008 Lyon, France; Cypath & Cypath-rb; Villeurbanne, 69100, France; University of Lille, CHU Lille, Institut de Pathologie, Tumorothèque du Centre Régional de Référence en Cancérologie, Avenue Oscar Lambret, F-59000 Lille, France; Department of Pathology, CHU NORD, Marseille, France; Centre de Recherche des Cordeliers, Sorbonne Université, INSERM, Université de Paris, Team Inflammation, complement and cancer, F-75006, Paris, France and Department of Pathology, Hôpitaux Universitaire Paris Centre, Tumorothèque /CRB Cancer, Cochin Hospital, Assistance Publique - Hôpitaux de Paris, Paris, France; CHU of Nantes, Tumorothèque, Nantes, F44093, France; Université Côte d’Azur, Laboratory of Clinical and Experimental Pathology, Nice Center Hospital, FHU OncoAge, Biobank BB-0033-00025, and IRCAN Inserm U1081/CNRS 7284, 06002 Nice, France; Université Côte d’Azur, Department of Thoracic Surgery, Nice Center Hospital, FHU OncoAge, and IRCAN Inserm U1081/CNRS 7284, 06002 Nice, France; Nancy Regional University Hospital, CHRU, CRB BB-0033-00035, INSERM U1256, Nancy, France; Toulouse University Hospital, Université Paul Sabatier, Toulouse, France; Department of Pathology; Marie Lannelongue Hospital; Le Plessis Robinson; 92350 France; Hospices Civils de Lyon, Institut de Pathologie, Centre de Ressources Biologiques des HCL, Tissu-Tumorothèque Est, Lyon, France; CHU of Caen, 14 033 Caen; MESOPATH Regional Center; Centre de Pathologie des Côteaux, Centre de Ressources Biologiques (CRB Cancer), IUCT Oncopole, 31500, Toulouse; University of Lille, CHU Lille, INSERM, OncoThAI, NETMESO network, Lille, France; Department of Biopathology, CHRU Nancy, Vandoeuvre-les-Nancy, France and BRC, BB-0033-00035, CHRU Nancy, Vandoeuvre-les-Nancy, France; Centre de Recherche des Cordeliers; Inserm; Sorbonne Université; Université de Paris; Functional Genomics of Solid Tumors; Paris, 75006; France; Direction Santé Environnement Travail, Santé Publique France; Université Paris-Saclay, CEA, Centre National de Recherche en Génomique Humaine (CNRGH), 91057, Evry, France; Cologne Centre for Genomics (CCG), Cologne, Germany; Grenoble Alpes University; Owkin, Inc., New York, NY, USA; UMR INSERM 1052, CNRS 5286, UCBL1, Cancer Research Center of Lyon, Centre Léon Bérard, 69008 Lyon, France, Lyon Cancer Research Center (CRCL); Institut Curie, Institut du thorax Curie Montsouris, Paris, France, and UVSQ, Paris Saclay, Versailles, France; Institució Catalana de Recerca i Estudis Avançats (ICREA), Barcelona, Spain

**Keywords:** Malignant Pleural Mesothelioma, genome, transcriptome, epigenome, heterogeneity, classification, tumor evolution, cancer task

## Abstract

Malignant Pleural Mesothelioma (MPM) is an aggressive cancer with rising incidence and challenging clinical management. Using the largest series of whole-genome sequencing data integrated with transcriptomic and epigenomic data using multi-omic factor analysis, we demonstrate that MPM heterogeneity arises from four sources of variation: tumor cell morphology, ploidy, adaptive immune response, and CpG island methylator phenotype. Previous genomic studies focused on describing only the tumor cell morphology factor, although we robustly find the three other sources in all publicly available cohorts. We prove how these sources of variation explain the biological functions performed by the cancer cells, and how genomic events shape MPM molecular profiles. We show how these new sources of variation help understand the heterogeneity of the clinical behavior of MPM and drug responses measured in cell lines. These findings unearth the interplay between MPM functional biology and its genomic history, and ultimately, inform classification, prognostication and treatment.

**Graphical abstract:** 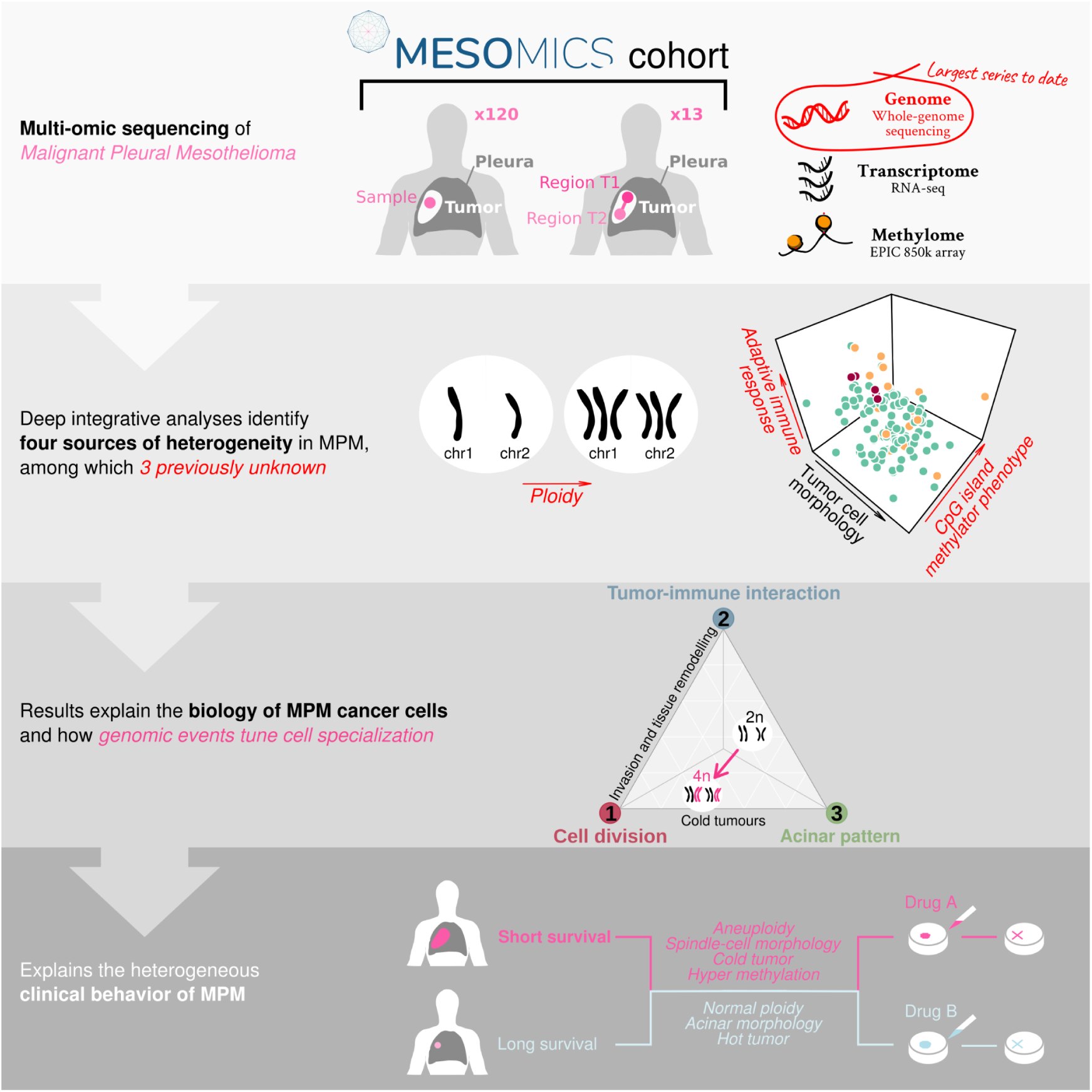

## Introduction

Malignant Pleural Mesothelioma (MPM) is a poorly-understood, rare, and aggressive disease associated with asbestos exposure (Carbone et al., 2019). The current WHO classification distinguishes three major histological types: epithelioid (MME), biphasic (MMB), and sarcomatoid (MMS) (IARC/WHO, 2015). In the past decade, knowledge on the molecular profile of MPM has rapidly expanded, owing to cohorts combining whole-exome sequencing, transcriptomic, and epigenomic data (Bueno et al., 2016; Hmeljak et al., 2018; de Reyniès et al., 2014). These genomic studies uncovered molecular profiles (clusters) related to MPM’s histopathological classification. Additional studies revealed a molecular continuum of types that explained the prognosis of the disease more accurately than discrete clusters (Alcala et al., 2019a; Blum et al., 2019). The clinical impact of these important findings has been limited by the vast morphological (Nicholson et al., 2020) and molecular heterogeneity of MPM (Fernandez-Cuesta et al., 2021), which remains largely unexplained. Several additional histopathological and molecular features have been described, such as variations between epithelioid histological subtypes (Nicholson et al., 2020), variable immune infiltration (Alcala et al., 2019a), and large-scale genomic aberrations such as aneuploidy (Hmeljak et al., 2018), and structural rearrangements (Mansfield et al., 2019). As new treatment opportunities are being made available, such as antiangiogenic agents and immunotherapies, with unpredictable benefits at the individual patient level, a better understanding of these aspects is mandatory.

Malignant transformation and cancer development depend on genomic aberrations that can result in a wide range of molecular profiles, and provide actionable treatment targets (Cortés-Ciriano et al., 2020; PCAWG Consortium, 2020; Quinton et al., 2021). Genomic events have not been fully described in MPM as previous efforts have been restricted to profiling only exomes or a reduced representation of genomes (Bueno et al., 2016; Hmeljak et al., 2018; de Reyniès et al., 2014). There is also a lack of comprehensive integrative analyses examining how molecular features affecting multiple omic layers, in particular genomic aberrations, interact to generate the observed heterogeneous tumor phenotypes. Biological functions performed by tumor cells, and the role of genomic events in shaping these functions remain largely unknown, hindering any meaningful progress in the diagnosis, classification, and treatment of the disease.

We have designed the MESOMICS study (http://rarecancersgenomics.com/mesomics/) to dissect MPM tumor heterogeneity, uncover its main sources of molecular variation, and identify its underlying biological functions. We characterize the impact of genomic aberrations on these biological functions, and use them to identify potential therapeutic opportunities. We performed multi-omics analyses combining genomic, transcriptomic, and epigenomic data, with detailed clinical and histopathological annotations, providing the most complete profile of MPM to date, avoiding blindspots in sources of variation. Taking advantage of the first cohort-level whole-genome sequencing data of 115 tumors, in addition to 109 transcriptomes, 119 epigenomes, and 13 multi-region samples, we mapped the genomic landscape of 120 mesotheliomas and characterized its implications for the molecular profiles. Replicating our findings in MPM cell lines previously tested for multiple drugs allowed us to identify therapeutic vulnerabilities across the spectrum of MPM heterogeneity.

### Integrative multi-omics analyses uncover three novel axes of molecular variation

To find the major independent molecular profiles underlying MPM heterogeneity, and infer associations between omics layers, we performed a Multi-Omics Factor Analysis (MOFA) (Argelaguet et al., 2020) including genomic, transcriptomic, and epigenomic data. MOFA identified four latent factors (LFs) explaining individually more than 10% of molecular variation (Figures 1A and **S1A-G; Table S2**). We found features from all omics layers associated with each LF, and only one, LF2, associated with the current WHO histopathological classification, the recent artificial intelligence score based on digital pathology (Courtiol et al., 2019), and the previously proposed molecular classifications (maximum *q*-value = 2.57×10^-10^; Figure 1B; Alcala et al., 2019; Blum et al., 2019; Bueno et al., 2016; Hmeljak et al., 2018; de Reyniès et al., 2014). This data suggests that these LFs inform interactions between omics layers, and capture novel molecular profiles.

**Figure 1.**
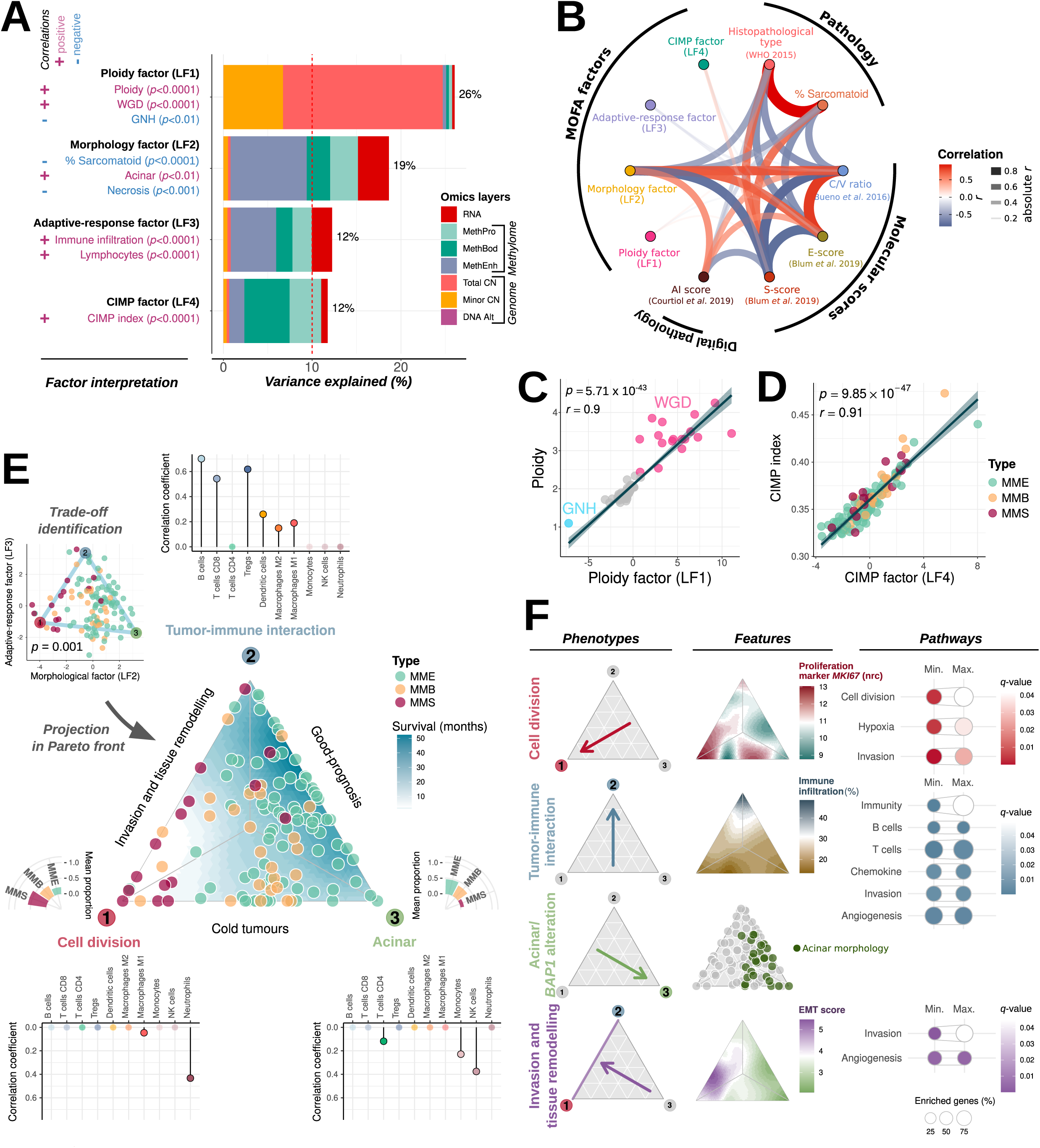
Multi-omics Factor Analysis (MOFA) of whole-genomes, transcriptomes, and methylomes of the MESOMICS cohort (n=120). (A) Interpretation of the four main latent factors (LFs) identified by MOFA, and the proportion of variance in each ’omic layer’ they explain. (B) Network of the correlations between LFs, tumor histopathology, and previously published molecular scores. Arc colors, width and transparency correspond to Pearson correlation coefficients. (C) Correlation between Ploidy factor (LF1) and ploidy. WGD: whole-genome doubling, GNH: genomic near-haploidization. (D) Correlation between GIMP factor (LF4) and GIMP index. (E) Trade-offs between cancer tasks identified from the Morphology (LF2) and Adaptive-response factors (LF3). Upper left: sample positions along LF2 and LF3 are contained within a triangle formed by three phenotypic archetypes (colored vertices). Middle: Ternary plot representing the sample’s distance from the three archetypes. Lollipop plots present the correlation between RNA-seq-estimated immune cell infiltration and proportion of archetypes. Bar plots represent the association between archetypes and histopathological types. (F) Summary table of the main phenotypes, features, and overexpressed pathways (columns) identified in each profile (rows). Left: arrows indicate the focal profile of each row. Middle: ternary plots with colour-filled background representing key features for each profile. nrc: normalized read count. Right: Cancer tasks inferred from each phenotype. Point colours correspond to the range (Min. - Max.) of the enrichment q-values from the corresponding pathways and point size to the proportions of enriched genes in each pathway. In (A), (B), (C), (D), p-value and coefficients r correspond to Pearson correlation tests. See also **Figure S1** and **Tables S2** and **S3.**

LF1, largely explained by copy number variants (CNVs), ranged from a genomic near-haploidization (GNH) sample to whole-genome doubled (WGD) samples (*q*-value = 3.3×10^-35^; Figure 1C). Aneuploidy was previously reported in the TCGA’s MPM cohort (Hmeljak et al., 2018), now captured by this axis. We found LF1 strongly correlated with ploidy (*r* = 0.90) and named it the Ploidy factor. As described below, LF2 summarizes the current knowledge on the molecular profiles of histological types (Figure 1A-B), and we therefore named LF2 the Morphology factor. Also described below, LF3 summarizes immune infiltration with adaptive response effectors (lymphocytes), separating “hot” (high infiltration, in particular of effector cells) from “cold” phenotypes (low immune infiltration), and was named the Adaptive-response factor (Figure 1A). For these two factors, enhancer methylation was the major omic contributor (explaining about half of the variance of LF2 and LF3; Figure 1A). This is partly explained by enhancer methylation implication in the Invasion-and-tissue-remodeling phenotype (see below), and its variability in MPM likely driven by cell-type heterogeneity (tumour cell type and immune cell type mixtures; **Figure S1H-J; Table S2**). The major contribution to LF4 came from methylation at gene body and promoter regions, and most of its molecular variation was strongly associated with the CpG island methylator phenotype (CIMP) index (*q*-value = 3.2×10^-35^; Figure 1D), thus we named it the CIMP factor.

Gene expression variation is mainly captured by the interdependent Morphology and Adaptive-response factors. Looking at these factors through the lens of multi-task theory (Hausser and Alon, 2020) provided a map of the biological functions performed by MPM cells, and the degree to which tumours specialize in each function. Interdependence between molecular axes occurs when tumor cells implement trade-offs between different tasks (Hatzikirou et al., 2012). Such trade-offs are expected to leave a specific footprint in omics data, where values in one axis are constrained by values in a second axis, forming specific geometric shapes named the “Pareto front” (such as triangles, or tetrahedra; Hausser et al., 2019). In the case of MPM, samples formed a robust triangle within the LF2 and LF3 space, demonstrating a significant trade-off between three tasks corresponding to the extreme profiles captured by LF2 and LF3 (Pareto fit model *p*-value = 0.001; upper left panel, Figure 1E). The projection of samples within this triangular Pareto front provides a map of MPM task specialization, where vertices, known as phenotypic archetypes, correspond to task specialists, and the center to multi-tasks generalists (central panel, Figure 1E).

Archetype 1 (Arc-1) corresponds to the Cell division phenotype, with tumors closest to Arc-1 displaying upregulation of pathways within the universal “cell division” task (Hatzikirou et al., 2012) identified through Integrative Gene-Set Enrichment Analysis (IGSEA; maximum *q*-value = 3.8×10^-2^; Figure 1F first row**; Table S3**). This archetype was enriched for sarcomatoid tumors and biphasics with a large sarcomatoid component, with samples presenting higher levels of necrosis, higher grade, and greater percentage of infiltrating neutrophils (maximum *q*-value = 2.29×10^-2^). Arc-1 was associated with high expression levels of the proliferation marker *MKI67*, and increased genomic instability (estimated from genomic, transcriptomic and epigenomic data; maximum *q*-value = 0.001). Arc-2 is the Tumor-immune-interaction phenotype as supported by upregulated immune-related pathways identified with IGSEA (maximum *q*-value = 3.1×10^-2^; Figure 1F second row; **Table S3**), and high immune infiltration with an enrichment for adaptive-response cells: lymphocytes B, T-CD8+, and T-reg (maximum *q*-value = 3.11×10^-9^). Arc-3 was named the Acinar phenotype upon its enrichment in samples of this epithelioid subtype, and the few upregulated pathways based on IGSEA (Acinar association *q*-value = 0.009; Figure 1F third row; **Table S3**). In line with the better prognosis reported for this subtype (Nicholson et al., 2020), the Acinar phenotype is characterized by the highest levels of global methylation (*q*-value = 3×10^-10^); global hypomethylation is a characteristic of many cancers and is known to occur during rapid cell division and growth (Shipony et al., 2014). The Cell-division and Tumor-immune-interaction phenotypes showed common enrichment for pathways in the Invasion-and-tissue-remodeling universal cancer task (maximum *q*-value = 4.09×10^-2^; Figure 1F fourth row**; Table S3**). We also observed a higher Epithelial-to-Mesenchymal Transition (EMT) score amongst tumors closest to these phenotypes, driven by upregulation of mesenchymal genes and hypomethylation of their associated enhancers (maximum *q*-value = 6.22×10^-5^). In MPM, *in vitro* studies have shown that asbestos may induce EMT (Turini et al., 2019) and in line with this, we found a positive correlation between mesenchymal genes expression and asbestos exposure score, and a negative correlation between mesenchymal gene enhancer methylation and asbestos exposure score (Pearson’s correlation coefficient *r* = 0.44, *q*-value = 0.01, and *r* = -0.33, *q*-value = 0.02, respectively). The Cell-division and the Acinar phenotypes were both correlated with the presence of innate immune response cells, but of different individual types: neutrophils for the Cell-division phenotype, and monocytes and NK cells for the Acinar phenotype.

### Multi-regional sequencing reveals variable intra-tumoral heterogeneity in the Morphology, Adaptive-response, and CIMP factors

Using a multi-regional sub-cohort from 13 patients, we inferred intra-tumoral heterogeneity (ITH) in MPM (**Table S4**) and observed that ITH can be greater than inter-tumoral heterogeneity in all molecular axes except the Ploidy factor (Figures 2A **and S2A**), and affected most omic layers except the genome (**Figure S2B-C**). This heterogeneity matched pathological annotations and impacted tumor specialization, that is, movement within the Pareto front (Figure 2B, arrow length corresponding to task specialization ITH matches arrow width, corresponding to histopathological ITH). Interestingly, ITH detected along the Morphology factor is driven by two alternative sources: the tumor cell morphology, as observed in between-region variation in EMT score (sample MESO_002, Figure 2C**, left**) and the innate immune response, as observed in between-region variation in the difference between neutrophil and monocytes and NK cells (sample MESO_052, Figure 2C**, right**). In general, the strongest ITH was due to different immune infiltration profiles, including the small changes in innate response cell compositions (7%) shown in Figure 2C and stronger changes in adaptive immune response cell proportions (in particular macrophages M1; Figure 2D). Substantial ITH in the CIMP index was also found in three out of the 13 patients (Figure 2E), either in conjunction with regional differences in histopathological type, immune infiltration, or no detectable difference.

**Figure 2.**
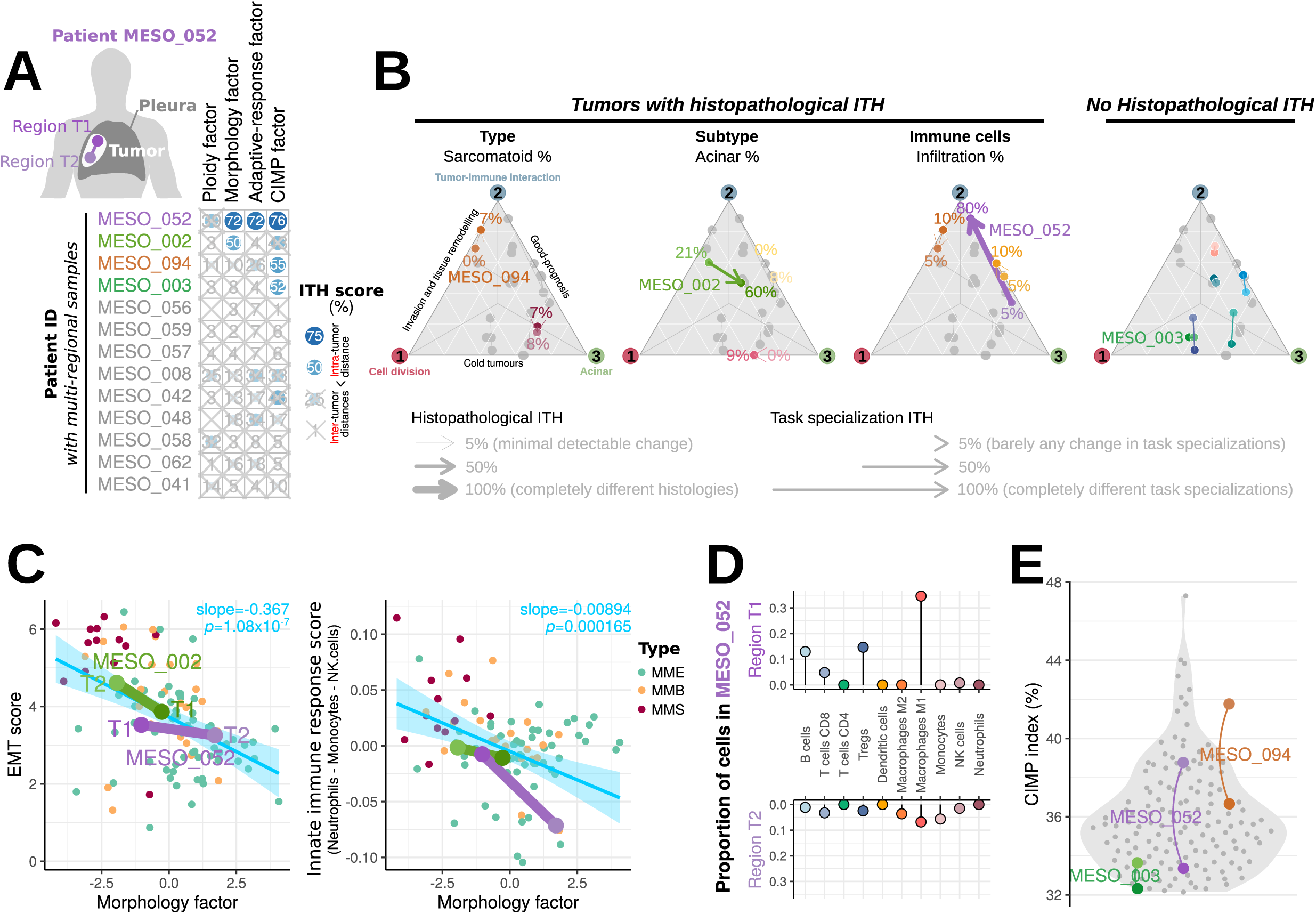
Multi-omic intra-tumor heterogeneity (1TH) of 13 multi-region samples. (A) 1TH score, ranging from 0% (no 1TH) to 100% (1TH greater than the maximum observed inter-tumor heterogeneity in the cohort), for each sample (row) and each MOFA latent factor (column). The score is computed as the percentage of inter-tumor distances in a MOFA factor that are lower than the observed intra-tumor distance between regions. The four samples with 1TH score greater than 50% are highlighted in color. (B) Relationship between histopathological heterogeneity and cancer task specialization. Ternary plots depicting task specialization in three cancer tasks (see Figure 1E). For each histopathological feature, a colored arrow connects regions from tumors with differences in this feature. Numbers correspond to the percentage of this feature in the tumor as estimated by our pathologist. The right ternary plot represents all samples with no histopathological 1TH. (C) EMT score and innate immune composition score as a function of MOFA’s Morphology factor . Small points correspond to all samples from the MESOMIGS cohort, and large points connected by segments to regions from the 3 patients with GIMP factor 1TH highlighted in (A). (D) Lollipop plot of the estimated proportion of immune cells in two regions of a sample with 1TH in the adaptive-response factor highlighted in (A). (E) GIMP index in regions of two tumors with substantial 1TH in the GIMP factor highlighted in (A) (colored points connected by an arc), compared to that of the rest of the cohort (grey points). See also Figure 52 and **Table 54.**

### WGS uncovers a heterogeneous genomic landscape and characteristic MPM drivers

In our MESOMICS series, which is the largest WGS cohort of MPM to date, we identified a wide range of large-scale genomic events for 111 out of 115 samples with available data (97%) (Figure 3A). As captured by the Ploidy factor, MPM samples have various ploidies, ranging from haploid to tetraploid (Figure 3B). The average CNV profile is highly consistent between cohorts (**Figure S3A**), with several recurrent chromosome arm-level CNVs, as well as focal alterations (deletions, del; amplifications, amp) encompassing known cancer genes: *BAP1* del (chr *3p21.1*), *TERT* amp (chr *5p*), *EZH2* del (chr *7q36.1*), *CDKN2A*/*B* and *MTAP* del (chr *9p21.1*), *RBFOX1* del (chr *16p13.3*), and *NF2* del (chr *22q*) (Figure 3C**; Table S8**). While *CDKN2A*/*B* and *MTAP* presented mostly homozygous deletions, *NF2* and *BAP1* were more often affected by heterozygous deletions (Figure 3A, left panel; **Table S7**); both events impacted gene expression levels (Figure 3A, middle-right panel). As previously reported (Chapel et al., 2020), most of the *MTAP* alterations co-occurred with *CDKN2A*/*B* deletions with only five and six samples, respectively, presenting alterations in *MTAP* or *CDKN2A/B* exclusively (Figure 3A; **Table S7**). In addition, we found recurrent deletions of a prominent immune recognition gene, *B2M* (chr *15q14*; Figure 3C).

**Figure 3.**
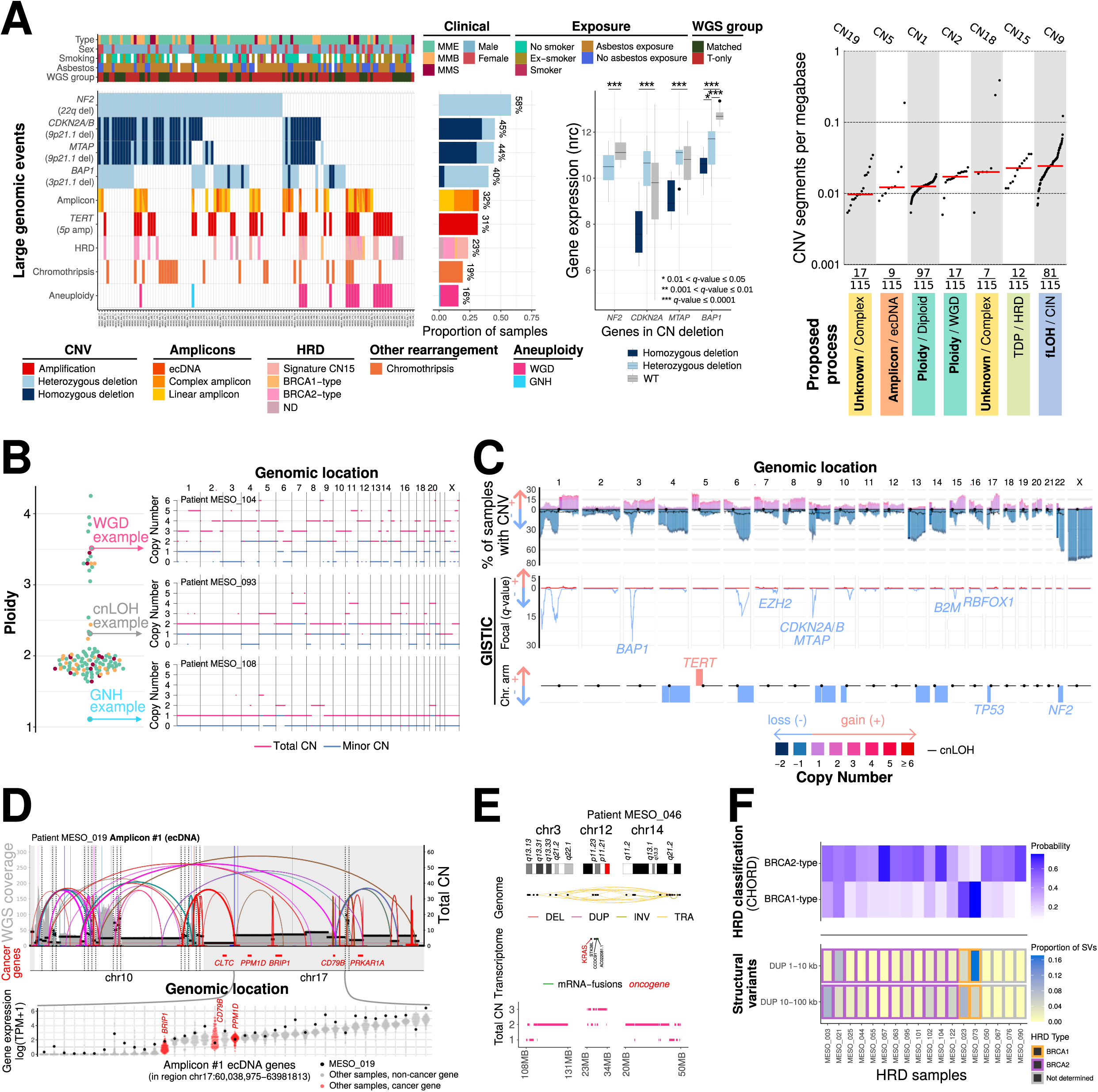
Genomic characterization of MPM from the ME50MIC5 cohort. (A) Recurrent large genomic events. Upper panel: clinical, epidemiological, morphological, and technical features per sample. Far left: oncoplot describing genomic events per sample. Middle-left panel: barplot of the frequency of each event within the cohort. Middle-right panel: comparison of gene expression (in normalized read count, nrc) of cancer-relevant genes belonging to frequent deletions detected by GISTIC, with regards to their CN status. Far right panel: CNV signatures identified in the cohort, from Steele et al. (2021). TOP: tandem duplicator phenotype; fLOH: focal LOH. (B) Left: ploidy spectrum of the cohort. Right: three extreme ploidy cases, a genomic near haploidization sample (GNH, bottom), a sample with high levels of copy neutral LOH (cnLOH, middle), and a sample with WGD (top). (C) Cohort level copy number profile (top-track), with significantly targeted regions identified by GISTIC, in focal peaks (middle track) and at chromosome arm level (bottom track). (D) Reconstructed ecDNA (amplicon) structure (top) and its impact on gene expression (bottom). Black horizontal bars represent the CN profile and colored arcs represent structural variants (SVs). Expression of genes located within the ecDNA region are ordered by median TPM. The red dots display the expression of oncogenes found within the ecDNA region. (E) Example of a region displaying a chromothripsis pattern (cluster of SVs, a CN profile that oscillates, and interspersed LOH) and generating mRNA fusions. (F) Overview of HRD samples predicted by CHORD. The upper panel displays the probability for BRCA1 and BRCA2 HRD types. The lower panel indicates the proportions of duplications (DUP), binned by length (1-10kb and 10-100kb) out of all SVs, observed in each positive HRD sample and the predicted HRD type classes. See also Figure 53 and **Tables 55, 56, 57, 58,** and **59.**

CNV signatures (Steele et al., 2021) illustrated the processes leading to the deletions and amplifications, but also the heterogeneity of chromosomal rearrangements affecting MPM, such as those resulting in extrachromosomal DNA (ecDNA) and chromothripsis (Figure 3A, right panel; **Table S9**). We found CN9 to be positively correlated with focal *CDKN2A*/*B* and *BAP1* deletions (Pearson’s correlation coefficient *r* = 0.23, *q*-value = 0.039 and *r* = 0.27, *q*-value = 0.02 respectively), in line with recent data linking this signature with *CDKN2A*/*B* deletions in breast cancer (Steele et al., 2021). CN5 was associated with ecDNA in our cohort (Figure 3A, right panel). Oncogenes encoded on ecDNA are among the most highly expressed genes in tumor transcriptomes (Wu et al., 2019). Consistently with this, and despite the general pattern of alterations compatible with a disease driven by the inactivation of TSGs (Figure 3C), the one ecDNA sample with available transcriptomic data (out of the six MPM with ecDNA, **Figure S3B**) showed an increased expression of the genes predicted to be part of the ecDNA sequence, including the known oncogene *BRIP1* (Figure 3D). ecDNA can be driven by kataegis (Bergstrom et al., 2021). In line with this, we observed that the aforementioned ecDNA sample co-occurred with and might be fuelled by kataegis (**Figure S3C**), despite the rarity of kataegis in our cohort, contributing to only 2% of the MPM clustered mutations (**Table S9**).

CN18 and CN19 are associated with complex CNV patterns, such as chromothripsis (Steele et al., 2021). In our cohort, a pattern compatible with chromothripsis was observed in 19% of the samples (Figure 3A**, Figure S3D; Table S9**), and this pattern was also observed at the transcriptomic level as fusion transcripts, in half of the positive samples (see example in Figure 3E**; Figure S3E; Table S6**). A signature of clustered structural variants (SVs) was detected and was significantly associated with a high SV load and chromothripsis (**Figure S3F; Table S6**). CN15 corresponded to the tandem-duplicator phenotype and homologous recombination-deficiency (TDP/HRD) signature, which was not associated with *BAP1* status in our series. Overall, 23% of the samples showed an HRD phenotype, identified either by CN analyses or other previously validated methods (Ladan et al., 2021) (Figure 3F; **Table S9**).

Mutational signature analysis of single base substitutions (SBS) identified 10 previously reported COSMIC signatures (**Figure S3G**), and we found none of them significantly associated with asbestos exposure, as previously reported (Bueno et al., 2016; Hmeljak et al., 2018). Although APOBEC signature activity was low in our cohort, we identified six samples with APOBEC-related signatures (SBS2/SBS13) that might be sensitive to epigenetic drugs (Levatic et al., 2021).

Despite the low mutational rate (0.98 non-synonymous single-nucleotide variants, SNVs, per megabase, **Figure S4A; Table S5**), MPM tumors carry a particularly high number of SVs relative to tumors with similarly low mutational burden (Figure 4, top panel, **Figure S4B**). The top genes altered by SVs (≥ 5%) were *RBFOX1, NF2, BAP1, MTAP*, and *PCDH15* (**Figure S4C;** Figure 4). Closer examination of the *RBFOX1* rearrangements reveals that 14 out of 52 samples have two separate events with most of them deleting CDS number 6 (**Figure S4D**), which encodes part of the RNA binding protein domain. Many of these genomic rearrangements resulted in fusion transcripts detected at the transcriptomic level; tumor suppressor genes (*MTAP, BAP1,* and *NF2)* were the most frequently affected by fusion transcripts (**Figure S4E**).

**Figure 4.**
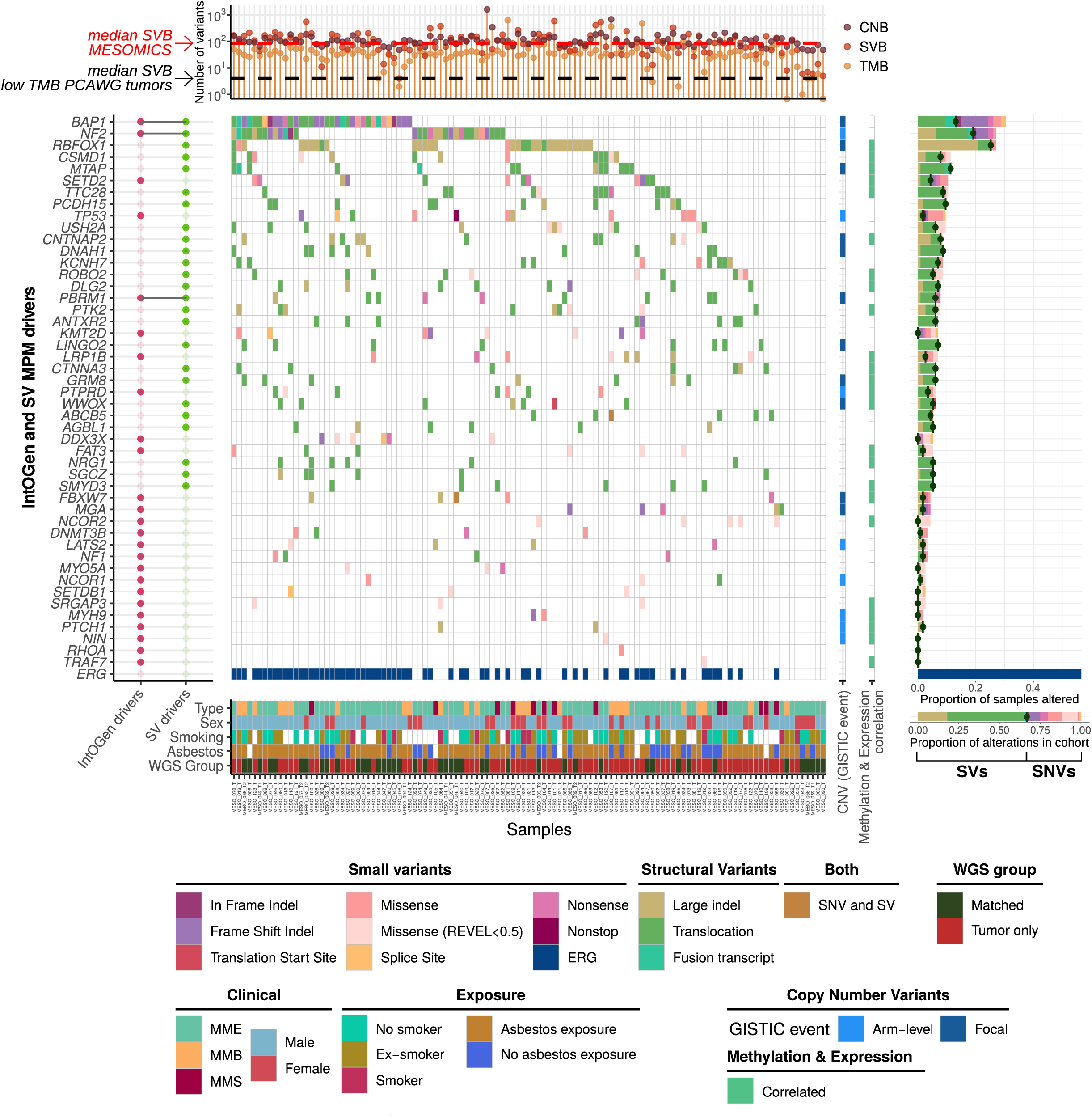
MPM driver genes in the MESOMICS cohort. Top: tumor mutational burden (TMB), CN burden (CNB), and SV burden (SVB) of each sample. Main: oncoplot describing genomic alterations in lntOGen and SV MPM driver genes per sample. Bottom row (ERG) indicates if the sample has one or more alterations in an epigenetic regulatory gene. Each gene is also annotated for belonging to one focal or arm-level GISTIC event, as well as for being regulated by DNA methylation (right bars). Right: frequency of alterations within the cohort. Bottom: key clinical, epidemiological, morphological, and technical features of each sample. See also **Figure S4** and **Tables S5, S6, S7, SB,** and **S9.**

Combining the MESOMICS dataset with the additional two other large datasets from Bueno et al. (2016) and the TCGA (Hmeljak et al., 2018), we reached the sample size (*n*≈300) needed to detect low frequency (1%) MPM driver genes in such low-mutated tumors. We used the well-established Integrative OncoGenomics (IntOGen) pipeline on point mutations and indels (Martínez-Jiménez et al., 2020). Thirty genes were identified as putative driver genes (**Figure S4F**). Five genes - *BAP1, NF2, SETD2, TP53*, and *LAST2* - were called in the three series individually and in the combined analysis, and are all known MPM altered genes. Among the other 25 genes, some had been previously reported as recurrently mutated in MPM (*PBRM1, KMT2D, DDX3X, PIK3CA, FBXW7, MGA, NF1, SETDB1, MYH9, PTCH1, RHOA,* and *TRAF7*; De Rienzo et al., 2016; Kato et al., 2016; Shukuya et al., 2014), or altered by SVs (*PTPRD* and *LRP1B;* Mansfield et al., 2019), two were previously found overexpressed in cell lines but not mutated (*DNMT3B* and *EZH2*; McLoughlin et al., 2017), and for some, germline mutations have been discovered, suggesting they may be genetic susceptibility genes (*NCOR1*; Pastorino et al., 2018; *MYO5A*; Hylebos et al., 2018). The remaining eight driver genes have, to our knowledge, not been previously reported in MPM, but are all known cancer genes as reported in COSMIC: *FAT3, NIN, ARHGAP5, HLA-A, NCOR2, SRGAP3,* and *WNK2*. Beyond extending the list of putative MPM drivers, combining point mutations with SVs allowed for the refinement of the frequency of altered key MPM genes (Figure 4, shade of green in right panel; **Tables S5-6**).

### Genomic alterations tune the molecular profiles of MPM

Most of the key identified genomic alterations were associated with the MOFA LFs (Figure 5A; **Table S11**). In addition to ploidy, *NCOR2* alterations and *TERT* amp were associated with the Ploidy factor (maximum *q*-value = 7.5×10^-4^; Figure 5A, left panel). While no association was previously detected between *TERT* promoter mutations and WGD (Bielski et al., 2018), here we found that both *TERT* amp and expression were associated with WGD events (*p*-value = 1.6×10^-10^, Fisher’s exact test, *p*-value = 0.009, linear regression, respectively; **Figure S5A**). Differential gene expression analyses showed that the most up-regulated enriched pathways in WGD+ *vs* WGD- MPM tumors were E2F targets, G2M checkpoints, myogenesis, downregulation of KRAS signaling, and glycolysis (maximum *q*-value = 0.04; Figure 5B**; Table S10**), revealing a specific profile with tumor vulnerabilities (Quinton et al., 2021).

**Figure 5.**
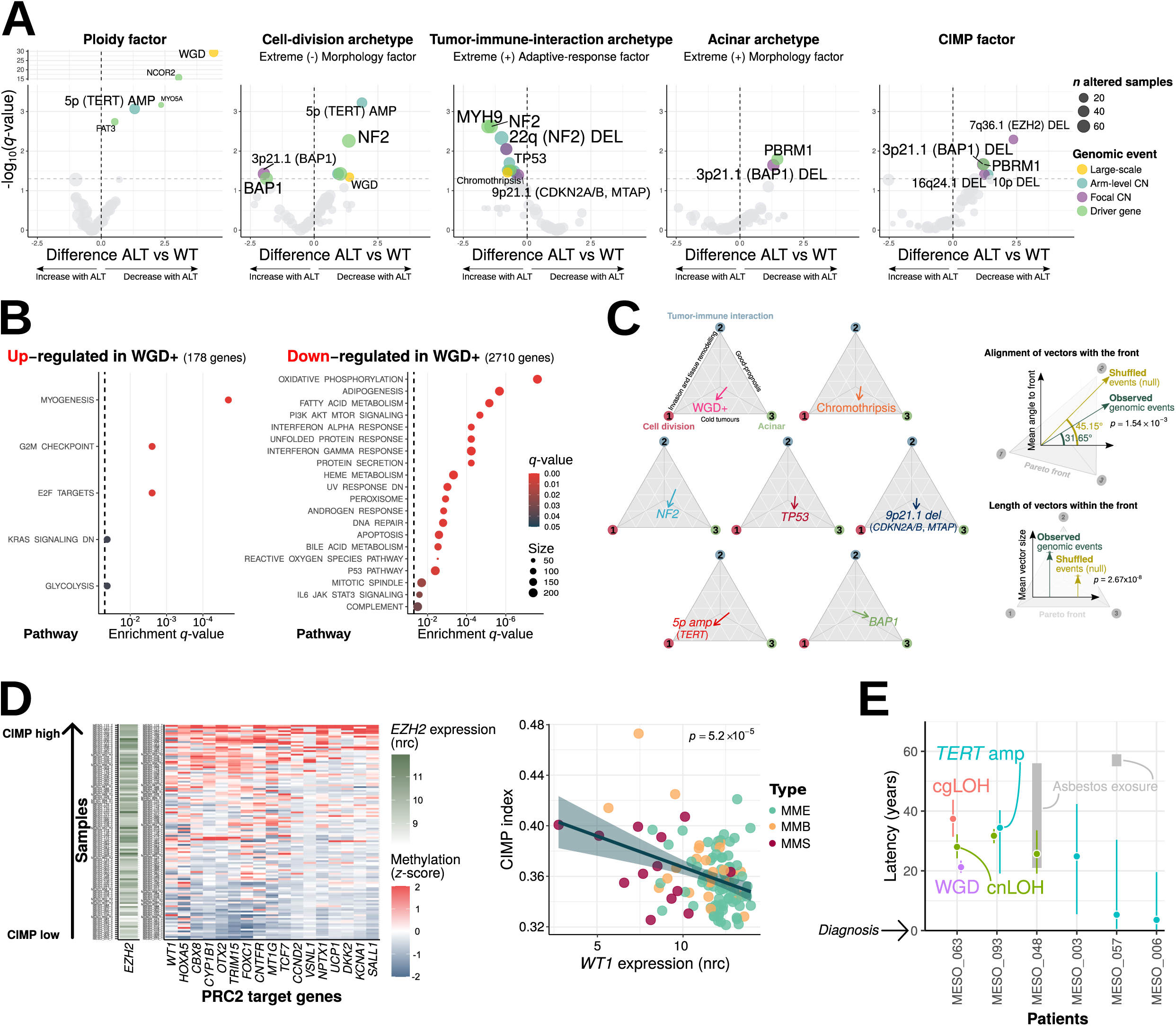
Impact of genomic events on MPM molecular profiles. (A) Associations between genomic events and MOFA factors. For each event, the ALT vs WT difference corresponds to the difference between the factor value of wild-type samples and that of altered samples, and q-values correspond to an adjusted ANOVA p-value. (B) Pathway enrichment analysis (hypergeometric test) of differentially-expressed genes in WGD compared to non-WGD tumors. (C) Left: effect vector of key alterations affecting specialization in tumor tasks from Figure 1E. Right: alignment of vectors with the Pareto front in degree (0°: perfectly aligned, 90°: completely orthogonal) and length of the vector. p -values correspond to Wilcoxon tests between observed and shuffled vector distributions. (D) Association between GIMP index, EZH2 expression (n=109 samples), and PRG2 target gene methylation (n=119 samples). Left: heatmaps of *EZH2* gene expression (nrc) and GpG island methylation (z-score) of PRG2 target genes whose methylation level was significantly positively correlated with GIMP index (q S 0.05), for samples ordered by GIMP index. Right: correlation between *WT1* expression and GIMP index with p-value for Pearson correlation test. (E) Latency between large-scale amplifications and age at diagnosis. cnLOH: copy neutral loss of heterozygosity; cgLOH: copy gain LOH; WGD: whole-genome doubling. Points represent estimates of the timing of genomic events, and bars, 95% credibility intervals obtained through Bayesian inference. Grey rectangles indicate known exposure times. See also **Figure S5** and **Tables S10, S11,** and **S12.**

All types of genomic events were associated with the position of the samples in the Pareto front (Figure 5A, three middle panels). Multi-task evolution theory further allows to quantify how alterations tune tumor specialization by computing effect vectors –difference in position between the position of altered and wild-type samples– for each alteration within MOFA space. Alignment of vectors with the Pareto front indicates that the alterations tune specialization, and their lengths indicate the strength of the specialization (Hausser et al., 2019). We found that genomic events highlighted in Figure 5A were significantly aligned with the front and had significantly large sizes (permutation test vs shuffled vectors, *p*-values of 1.54×10^-3^ and 2.67×10^-8^, respectively; Figure 5C). WGD and chromothripsis tended to specialize tumors towards the Cell-division and Acinar phenotypes, both characterized by a “cold” phenotype (low immune infiltration) (maximum *q*-value = 0.05; Figure 5*C*). WGD+ MPM tumors had downregulation of the interferon response pathway (*q*-value = 5.8×10^-5^; Figure 5*B*), which might explain this “cold” phenotype (Quinton et al., 2021). We identified *B2M* as the second most downregulated gene in WGD+ tumors (**Figure S5*C***). B2M is part of the MHC-I being involved in the presentation of peptide antigens to the immune system (Sreejit et al., 2014); the downregulation of *B2M* and interferon genes might be important mechanisms for WGD+ tumors to avoid the immune response. Chromothripsis has also been associated with low immune infiltration as part of the chromosomal chaos that silences immune surveillance (Zanetti, 2017). *NF2* alterations, chr *9p21.1* del (*CDKN2A*/*B* and *MTAP*) and *TP53* alterations also converged upon “cold” tumours (maximum *q*-value = 0.04). *TERT* amp moved tumors towards the Cell-division phenotype (*q*-value = 6.6×10^-4^; Figure 5*C*). Mutations in the *TERT* promoter have been previously described in MPM and were associated with the non-epithelioid types and shorter survival (Pirker et al., 2020; Quetel et al., 2020), and *TERT* overexpression has been shown to promote EMT (Liu et al., 2013), angiogenesis (Zhou et al., 2009), and cancer cell proliferation (Choi et al., 2008) in other cancer types. Here, we found 29 samples (25%) with *TERT* amp as part of a chr *5p* amp event, which led to increased expression of the gene (*p*-value = 1.8×10^-5^; **Figure S5A**). In total, 36 MPMs exhibited an increased expression of *TERT* (31%), with chr *5p* amp as the main underlying mechanism (80%). Finally, chr *3p21.1* del encompassing MPM drivers *BAP1*, *DNAH1*, and *PBRM1*, as well as *BAP1* mutations moved tumors towards the Acinar phenotype (maximum *q*-value = 0.02; Figure 5C).

We found enrichment for epigenetic regulator genes (ERGs), including *NCOR2* and *EZH2*, among the genes whose expression was significantly positively correlated with the CIMP index (*p*-value = 0.003). Moreover, chr *7q36.1* del, encompassing *EZH2,* further tuned the position of the samples along the CIMP factor (*q*-value = 5.6×10^-3^; Figure 5A). EZH2 is a histone methyltransferase and MPM driver that functions as part of the PRC2 complex to promote gene silencing of specific targets (Margueron and Reinberg, 2011). Indeed, genes whose CpG island methylation level was highest in CIMP-high tumors were enriched for PRC2 target genes (*p*-value = 0.01; Figure 5D). Nine out of the 30 driver genes were also ERGs, and GSEA confirmed a significant enrichment for ERGs in the IntOGen drivers lists (Fisher’s exact test *q*-value = 2.3×10^-8^), further supporting the role of this category in MPM beyond the CIMP index (**Figure S5B**). A recent pan-cancer and multi-omics study has shown that ERGs, when disrupted through genetic or non-genetic mechanisms, may act as drivers (“epidrivers”) in cancer development (Halaburkova et al., 2020).

A well-known effect of the CIMP-high phenotype is epigenetic silencing of TSGs (Baylin and Jones, 2016). We identified five COSMIC TSGs (Sondka et al., 2018), whose expression was both negatively correlated with the CIMP index and the methylation level of their CpG island(s): *CBFA2T3*, *FBLN2*, *PRF1, SLC34A2,* and *WT1* (Pearson’s correlation maximum *q*-value = 0.028, **Table S12**). Particularly interesting is *WT1*, a PRC2 target for which a vaccine against is currently being assessed in clinical trials for mesothelioma (Zauderer et al., 2017).

The specialization of tumors was influenced by early genomic events. Using the proportion of small variants present in one or multiple copies allowed us to infer the relative timing of CN gains (**Figure S5D-F**), and following Zhang et al. (2021b) who focused on similarly low tumor mutational burden (TMB) tumors, and given that the small variants accurately followed a simple linear model of accumulation as a function of patient age (**Figure S5G**), we used an estimated acceleration rate of 1x to time the events in an individual’s lifetime. WGD and chr *5p* amp (containing *TERT*) that influenced the Ploidy, Morphology, and Adaptive-response factors, can occur up to 20-30 years before diagnosis (Figures 5E). Similarly, *BAP1, NF2*, *CDKN2A*/*B* and *MTAP* CN losses that impact the tumor specialization map (Figure 5C), were preferentially clonal and thus likely to have happened early in tumor development (**Figure S5H**). This suggests that molecular profiles are constrained early during carcinogenesis by genomic events and therefore plasticity might be limited. Neutral tumor evolution was significantly associated with closeness to the Acinar phenotype (ANOVA *p*-value = 0.005; **Figure S5I**), with all neutrally evolving samples showing lack of BAP1 protein expression (**Table S1**) and two out of three harboring *BAP1* genomic alterations. *BAP1* small variants were all highly clonal (**Figure S5J**); nonetheless, as noted in previous studies (Quetel et al., 2020; Zhang et al., 2021a), we detected a minority of subclonal CNVs affecting the *22q* region encompassing *NF2*, suggesting that recent genomic events can in some cases further influence specialization.

### Axes of molecular variation explain the clinical heterogeneity of MPM

The four axes of molecular variation predicted the observed inter-patient heterogeneity in overall survival (OS) and efficacy of drugs. All the identified factors were orthogonal and associated with OS (Figures 6A and **S6A-D**; **Table S13**). The Ploidy factor associated WGD+ tumors with poor OS; the Morphology factor separated good and poor OS samples along the epithelioid-sarcomatoid continuum; the Adaptive-response factor linked “hot” tumors with better OS and also separated “cold” from “hot” sarcomatoids; and the CIMP factor associated CIMP-low tumors with a better OS (Figure 6A). We trained LF-based survival models and tested their performance over previously proposed prognostic factors in both the MESOMICS cohort (using 4-fold cross-validation to avoid overfitting) and the TCGA series (fitting models on the MESOMICS cohort and using TCGA as a purely external test set). Each factor individually provided a prediction value similar to that of the histopathological types, as well as that of other morphological and molecular, discrete and continuous, prognostic factors previously described (Figures 6B and **S6E-J**; **Table S13**). When combining the four factors there was an increase in their AUC value, suggesting that they capture molecular characteristics with independent prognostic value. This is supported by the performance of models using *(i)* the Morphology and CIMP factors and *(ii)* the Ploidy and adaptive-response factors, in predicting short- and long-term survivors, respectively (Figures 6C and **Figure S6E**, **H**). Interestingly, we found that *MKI67* gene expression, associated with the Morphology factor, allowed for clear separation between epithelioid with better and worse OS (left panel, Figure 6D, **Table S2**) while the four-factor model distinguished bad from good OS samples in the tested series (right panel, Figure 6D).

**Figure 6.**
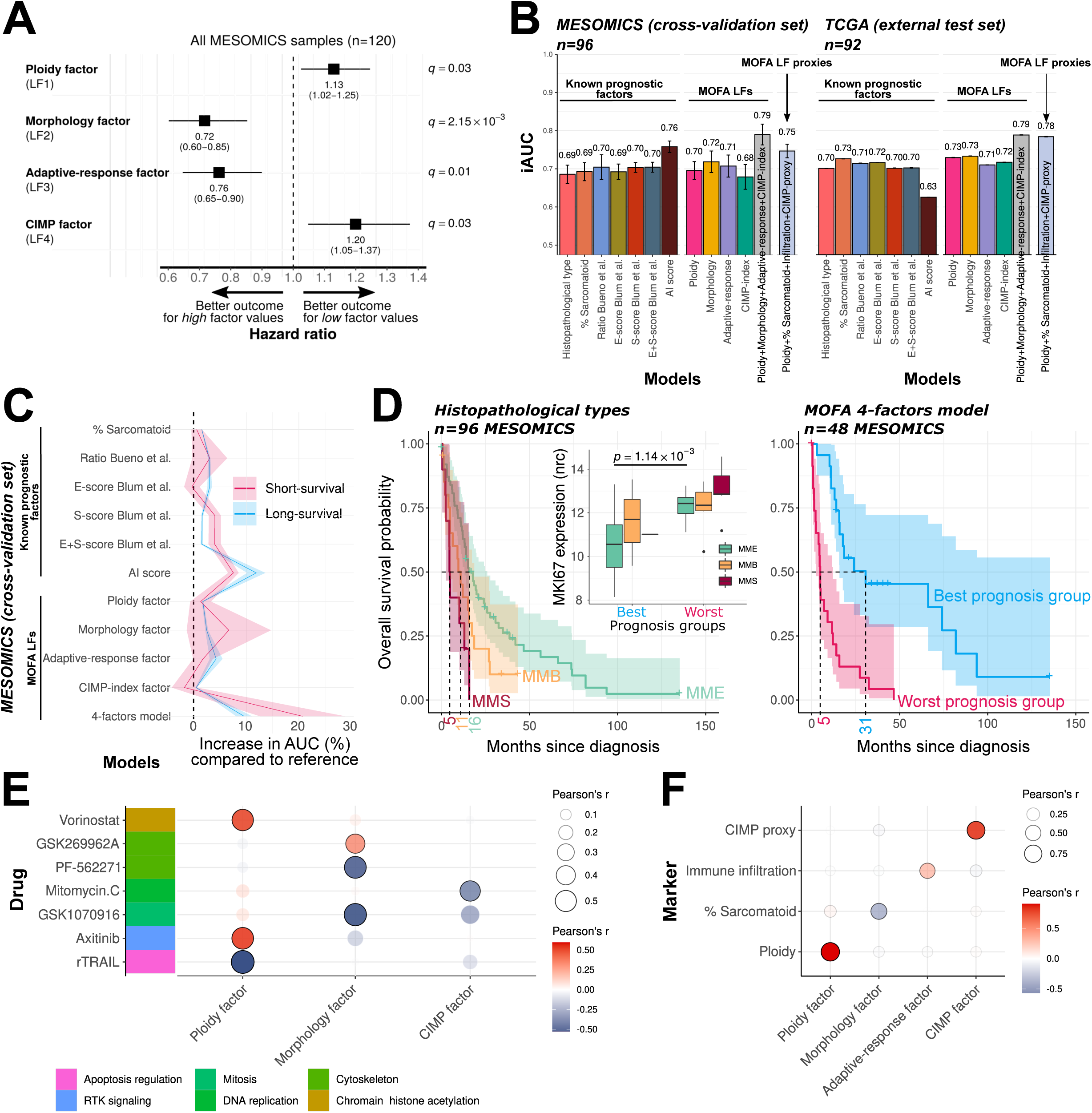
Clinical relevance of the four identified factors. (A) Forest plot of hazard ratios of MOFA latent factors for overall survival (OS). (B) Integral AUC (iAUC) of 11 Cox proportional hazards survival models: (i) a model based on the three histopathological types; (ii) a model based on the proportion of sarcomatoid content; (iii) a model based on the log2 of the *CLDN15*/*VIM* (C/V) expression ratio from Bueno et al. (2016); (iv-vi) models based on the E and S-score from Blum et al. (2019), individually and in combination; (vii) a model based on the AI prognostic score from Courtiol et al. (2019); (vii-xi) models based on each MOFA LF individually; (xii), a model based on all four MOFA LFs; (xiii) a model based on simpler proxies of the four MOFA LFs. Left: iAUC in the MESOMICS cohort, estimated using cross-validation. Right: iAUC estimated in the TCGA cohort. Bars correspond to standard errors of estimations (based on cross-validation sets for MESOMICS and bootstrapping for TCGA). (C) Increase in AUC in models (ii)-(xi) compared to model (i) as a function of OS, in two sample groups: long- (OS > 30 months) and short-survival (OS < 10 months), in the MESOMICS cohort. Increases in AUC are quantified as a percentage of change compared to the AUC of model (i) (D) Kaplan–Meier curves of OS for the three histopathological types (left) and prognostic groups from Cox model (xi) predicted using cross-validation (right), in the MESOMICS cohort. Dashed lines represent the median OS of each group. Prognostic groups were defined as the 25% of samples with the most extreme good and bad prognostic based on the cross-validated predictions of the Cox model (xi). Up: *MKI67* expression level in nrc in the good and bad-survival prognostic groups defined by Cox model (xi). (E) Correlations between drug sensitivity (IC_50_) of MPM cell lines and MOFA factors. Drugs have been selected from **Figure S7 H-I**, as being significantly correlated with at least one of the four factors and displaying an extreme factor weight within the empirical feature weight distribution (within 2 standard deviations of the mean). Of note, only drugs for which at least 10 cell-lines have been tested have been considered. (F) Correlation between four clinically accessible features and MOFA factors. See also Figures S6 and S7 and **Tables S13** and **S14**.

We used data from the Iorio et al. (2016), de Reyniès et al. (2014), and Blum et al. (2019) MPM cell lines to find candidate drugs for each molecular profile, combining molecular data and response to drugs available (265 for Iorio and three for de Reyniès and Blum; Figures 6E and **S7**; **Table S14**). The Ploidy, Morphology, and CIMP factors were accurately reproduced in the Iorio et al. cell lines, which had all ‘omic layers available, and the Morphology factor was also accurately reproduced in the de Reyniès et al. cell lines (**Figure S7A-G**). We found that drug responses associated with the different factors were entirely orthogonal: all 27 drugs with significant associations between IC_50_ and MOFA factors (before multiple-testing correction) were associated with a single factor (**Figure S7H**); this highlights that MOFA factors capture independent sources of heterogeneity in drug response. We found that the Ploidy factor presented the largest number of significant associations with drugs (19 out of 27; Figures 6E and **S7**), which further supports the importance of large-scale genomic variation to understand clinical behavior of the disease. Among these drugs, we found one receptor tyrosine kinase signaling compound axitinib, VEGFRi and the HDAC inhibitor Vorinostat, to which low-ploidy samples might be specifically sensitive. On the other hand, high-ploidy samples might be sensitive to the apoptosis regulator r-TRAIL. In the case of the Morphology factor, MMS-like samples seem to be sensitive to GSK269962A (in line with the results from Blum et al., 2019) targeting the cytoskeleton through ROCK1/2 inhibition. MME-like samples may be more sensitive to PF-562271, a drug targeting the cytoskeleton and GSK1070916 regulating mitosis. CIMP-high samples may be specifically sensitive to mitomycin C, a drug targeting uncontrolled tumor DNA replication. While the aforementioned agents may not be in the clinic so far for MPM, these data suggest that stratification of patients may be relevant for clinical trials assessing new drugs, and support the clinical value of the biological functions captured by these factors.

Although MOFA factors were computed from thousands of molecular features, they represent phenotypes that can be captured by much simpler features (Figure 6F). Any robust estimate of the ploidy could be a proxy for the Ploidy factor. The Morphology factor is correlated with the pathologist-estimated percentage of sarcomatoid cells in the tumor (*q*-value = 6.98×10^-11^), and the Acinar phenotype is correlated with BAP1 expression measured by immunohistochemistry (IHC) (*r* = -0.38, *q*-value = 5.02×10^-5^) (**Figure S7K**). The Tumor-immune-interaction phenotype is significantly correlated with the pathologist-estimated immune content (*r* = 0.50, *q*-value = 6.08×10^-8^). The CIMP factor can be approximated by a small panel of genes such as the five genes proposed by Weisenberger et al. (2006), found to be significantly correlated with the CIMP factor (*r* = 0.91, *q*-value = 4.69×10^-29^). Importantly, these four simple features allow predicting survival almost as well as the actual four LFs (Figure 6B, light blue bars).

## Discussion

MPM is a recalcitrant cancer with an expected OS of less than two years following diagnosis. This extremely poor prognosis is largely explained by the little progress made in its clinical management over the past decades. The EURACAN/IASLC effort to provide a more multidisciplinary classification of MPM highlighted the current limitations in diagnosis, prognostication and classification (Nicholson et al., 2020), highlighting the impact of heterogeneity.

Regarding classification, characterization and refinement of the histopathological type dimension has been the major focus of the previously published studies, and all molecular groups described in the two major genomic cohorts of MPM generated to date (Bueno et al., 2016; Hmeljak et al., 2018) are solely correlated with our Morphology factor. The recently developed whole-image based AI prognostic score, which represents state-of-the-art survival predictions (Courtiol et al., 2019) was only correlated with the Morphology factor as well, further confirming that this factor mainly captures the molecular variation related to the morphological and not the molecular features, contrary to what has been shown for other cancer types (Fu et al., 2020). Here, we have uncovered three additional independent sources of major molecular variation, all with prognostic value, within MPM samples from the French MESOBANK. These findings were replicated using the the TCGA and Bueno datasets comprising mostly American patients, as well as in available data for MPM cell lines (Blum et al., 2019; Iorio et al., 2016; de Reyniès et al., 2014), showing that the factors represent overlooked but robust processes present in all omic cohorts to date.

Regarding prognostication, we found that the four factors provided complementary information about survival, and taking them all into account led to the most accurate prediction of prognosis. WGD, tumor infiltration, and CIMP status influence survival in other cancers and are being considered for their classification (Zhang et al., 2021c), but not for MPM. Our results, while further supporting the value of a refinement of the current histopathological classification using molecular data, suggest that ploidy, immune infiltration, and CIMP status are promising features to consider for prognostication. They allow us to distinguish “cold” from “hot” sarcomatoid tumors with different prognoses, which may also influence treatment decision-making. They even outperform the recently developed AI score (Courtiol et al., 2019), specifically designed to predict survival from a large series of pathology slides using deep learning. Our data also suggest that some simple markers could be used to separate samples: in line with published studies (Ghanim et al., 2015), we confirmed that gene expression of the routinely assessed KI-67 protein could help stratify epithelioid tumors into survival groups.

Regarding treatment, chemotherapy remains the standard first-line therapy for most MPM patients and may also be used in the end-stage setting (Baas et al., 2015). Several novel therapies have been or are currently being tested in clinical trials (Dulloo et al., 2021; Gray, 2021), most with limited success despite initially promising preclinical data. The benefit of targeted therapies only exists if they are applied to the right population. This type of personalized medicine is particularly challenging to implement in rare aggressive cancers due to limited molecular studies, and the pressure clinicians feel to offer “something” to patients with these otherwise deadly tumors. Our study has provided some interesting hints for new therapeutic opportunities and also valuable information on how to define target populations that will benefit the most from specific therapies, whether in clinical trials or already approved for MPM. Several examples are given below.

Tumors with *BAP1* alterations are expected to carry high levels of *EZH2* (LaFave et al., 2015), suggesting that EZH2i could be an effective therapy for patients carrying *BAP1*-mutated cancers. Based on this, EZH2i were tested in *BAP1* inactivated MPMs and a clinical trial is currently ongoing (NCT02860286). The molecular criteria for inclusion is lack of nuclear BAP1 staining by IHC, or evidence of loss of function by gene sequencing. However, we did not see a negative correlation between the expression of *BAP1* and *EZH2*. Moreover, samples in Arc-3 enriched for *BAP1* alterations, did not show a particular profile of high *EZH2* gene expression level. Considering the above-mentioned link between *EZH2* expression and CIMP index, patients with tumors showing high CIMP index may benefit the most from EZH2i.

Nuclear BAP1 regulates homologous recombination (HR) and cells harboring HR deficiency switch to base excision repair, which is assisted by PARP to repair DNA single-strand breaks (Helleday, 2011). The observed synthetic lethality led to approval of PARPi in BRCA1/2 cancers (Ledermann et al., 2014). Despite the lack of evidence to support *BAP1* status as a *bona fide* predictor of sensitivity to PARPi, several clinical trials have tested PARPi in BAP1-deficient samples (Dulloo et al., 2021). Our data does not support the use of PARPi in these tumors. Indeed, there was no co-occurrence of HRD phenotype and *BAP1* alterations in our series.

Major progress has been seen in the use of immunotherapy, with recent results from the CheckMate 743 trial supporting its use as a first-line treatment option for all patients with MPM (Baas et al., 2021a). Follow-up analyses highlighted the heterogeneity of treatment efficacy among histological types, likely due to the very different tumor microenvironment of epithelioid and non-epithelioid MPM (Di Maio and Tagliamento, 2021; Baas et al., 2021b). Despite the statistically significant improvement in OS, less than a quarter of patients were alive three years post diagnosis (Baas et al., 2021a). The failure to identify appropriate populations may be masking the beneficial effect of immunotherapy in selected patients. Our data suggest that tumors in Arc-2 may respond best to immunotherapy as supported by their high infiltration, high tumor inflammatory score (TIS) (Damotte et al., 2019), and high levels of *CTLA4* and *PD-1*. The samples presenting the highest levels of *PD-L1*, a traditional immunotherapy target, were located around Arc-1, an archetype with almost no expression of *CTL4A* and *PD-1*, and with very little infiltration (“cold” tumors), and therefore very unlikely to respond to immunotherapies.

All sources of variation captured by molecular factors, that may be used to define targeted therapy populations, could be easily detected by molecular markers already available in the clinic. As a proxy for the Ploidy factor, ploidy may be assessed by FISH (Wuilleme et al., 2005). Pathologist-estimated percentages of sarcomatoid cells in the tumor could be used as a proxy for the Morphology factor, complemented by the already routine use of IHC to detect BAP1 expression. The Tumor-immune-interaction phenotype can be captured by the pathologist-estimated immune content, and the interdependence between phenotypes enables us to distinguish tumors with the Cell-division phenotype from the others by crossing the immune content with the sarcomatoid component for MMS or MMB samples, or with BAP1 IHC for MME samples. The CIMP factor may be easily detected using a CIMP panel assay such as the MethyLight detection panel of five genes proposed by Weisenberger et al. (2006).

A comprehensive molecular understanding of a disease, coupled with detailed clinical information to make sense of the observed molecular variation and heterogeneity, are key to the success of clinical management of any cancer type. We report an experimental and analytical design to quantify and interpret the impact of genomic events on the heterogeneity of tumor phenotypes and translate the results into the clinic. This approach expands upon our previous efforts using unsupervised dimensionality reduction to uncover continuous sources of variation blindly to the WHO classification (Alcala et al., 2019a) in two complementary aspects. Firstly, experimental design, with the inclusion of whole-genome sequencing data on top of expression and methylation data; and secondly, statistical design, with the use of multi-omics dimensionality reduction (following Argelaguet et al., 2018, 2020). This approach allowed us to detect features such as aberrant gene expression shaped by large scale genomic events (WGD, GNH), as well as driver alterations, that were previously excluded from histological and molecular classifications (clusters from Bueno et al., 2016 and Hmeljak et al., 2018). We also propose a downstream framework to identify cancer tasks from multi-omics profiles, expanding the application of multi-task evolutionary theory (Hausser and Alon, 2020) to multi-omics tumor latent factors, rather than expression latent factors. We expect this approach to be particularly beneficial compared to traditional approaches based on clustering and differential analysis between clusters, e.g., using consensus clustering as in Bueno et al. (2016), or iCluster+ as in Hmeljak et al. (2016) or other TCGA studies (TCGA Network, 2012, 2013, 2014), for cancer types including mixtures of different histopathologies, and in cases where current histopathological and molecular classifications are poorly correlated.

We encourage further research pursuing the biological understanding of MPM and its impact on clinical management. Bulk sequencing data did not allow us to completely disentangle the relative importance of tumor and microenvironment expression along the Morphology and Adaptive-response factors. In particular, to what extent the characteristic continuous molecular profiles we identified in MPM result from a heterogeneous mixture of specialist cells or a homogeneous population of generalist cells. Finally, further studies are needed to develop biomarkers for prognosis and treatment stratification from the candidate features we highlight, before any recommendations for clinical practice can be made.

## Acknowledgements

This study is part of the MESOMICS project within the Rare Cancers Genomics initiative (www.rarecancersgenomics.com). We thank the patients for donating their tumour specimens. The human biological samples and associated data were obtained from the French MESOBANK. We also thank Ricard Argelaguet for his advice in using MOFA; Hugues Begueret, Nathalie Rousseau, Didier Bozonnet, Eric Wasielewski, Gilles Clapisson, Christelle Bonnetaud, Kevin Washetine, Audrey Lupo Mansuet, Cyrille Cuenin, and Estelle Clermont for their contribution to the biorepository. The authors would like to acknowledge the American Association for Cancer Research and its financial and material support in the development of the AACR Project GENIE registry, as well as members of the consortium for their commitment to data sharing. Interpretations are the responsibility of study authors. The results published here are part based upon data generated by the TCGA Research Network: https://www.cancer.gov/tcga. We also thank the National Program for pleural Mesothelioma Surveillance (PNSM) and Santé Publique France. This work has been funded by the French National Cancer Institute (INCa, PRT-K 2016-039 to L.F.C. and M.F.), the Ligue Nationale contre le Cancer (LNCC 2017 and 2020 to L.F.C. and M.F.). L.M. has a fellowship from the LNCC. This work also benefited from support of the France Génomique National infrastructure, funded as part of the “Investissements d’Avenir” program managed by the Agence Nationale pour la Recherche (contract ANR-10-INBS-09).

## Author Contributions (CRediT – Contributor Roles Taxonomy)

Conceptualization, L.F.-C., M.F.; Methodology, L.F.-C., M.F., L.M., N.A., A.DG., A.S.-O.; Software, L.M., N.A., A.DG., A.S.-O, A.G.-P., A.K., E.N.B., C.V.; Validation, L.M., N.A., A.DG., A.S.-O.; Formal Analyses, L.M., N.A., A.DG., A.S.-O., A.G.-P., A.K., E.N.B., C.V., M.A., C.M., P.C. A.G.-P., F.G.-S.; Investigation, L.M., N.A., A.DG., A.S-O., A.G.-P., A.K., E.N.B., J.K., C.G., M.A., L.S., T.M.D., A.P., C.M., P.C.; Resources, N.L.S., S.B., S.T.E., F.D., M.B., M.-C.C., S.G.-C., D.D., C.G., V.H., P.H., J.Mo., St.L., J.Ma., V.T.deM., C.P., G.P., I.R., C.S., A.S., F.T., J.-M.V., A.G.S.I., Sy.L., F.G.-S.; Data Curation, L.M., N.A., A.DG., A.S.-O., C.V.; Writing – Original Draft, L.F-C., M.F., L.M., N.A., A.DG., A.S.-O.; Writing – Review & Editing, L.F.-C., M.F., L.M., N.A., A.DG., A.S.-O., A.G.-P., A.K., H.H.-V., C.Ca., N.G., N.L.-B., L.B.A., F.G.-S.; Visualization, L.M., N.A., A.DG, A.S.-O.; Supervision, L.F.-C., M.F., N.A., H.H.-V., C.Ca., N.L.-B., L.B.A.; Project Administration, L.F.-C., M.F., L.M., N.A., M.-C. M., A.B., J.-F.D., J.A., P.N., A.G.; Funding Acquisition, L.F.-C., M.F., N.A.

## KEY RESOURCES TABLE

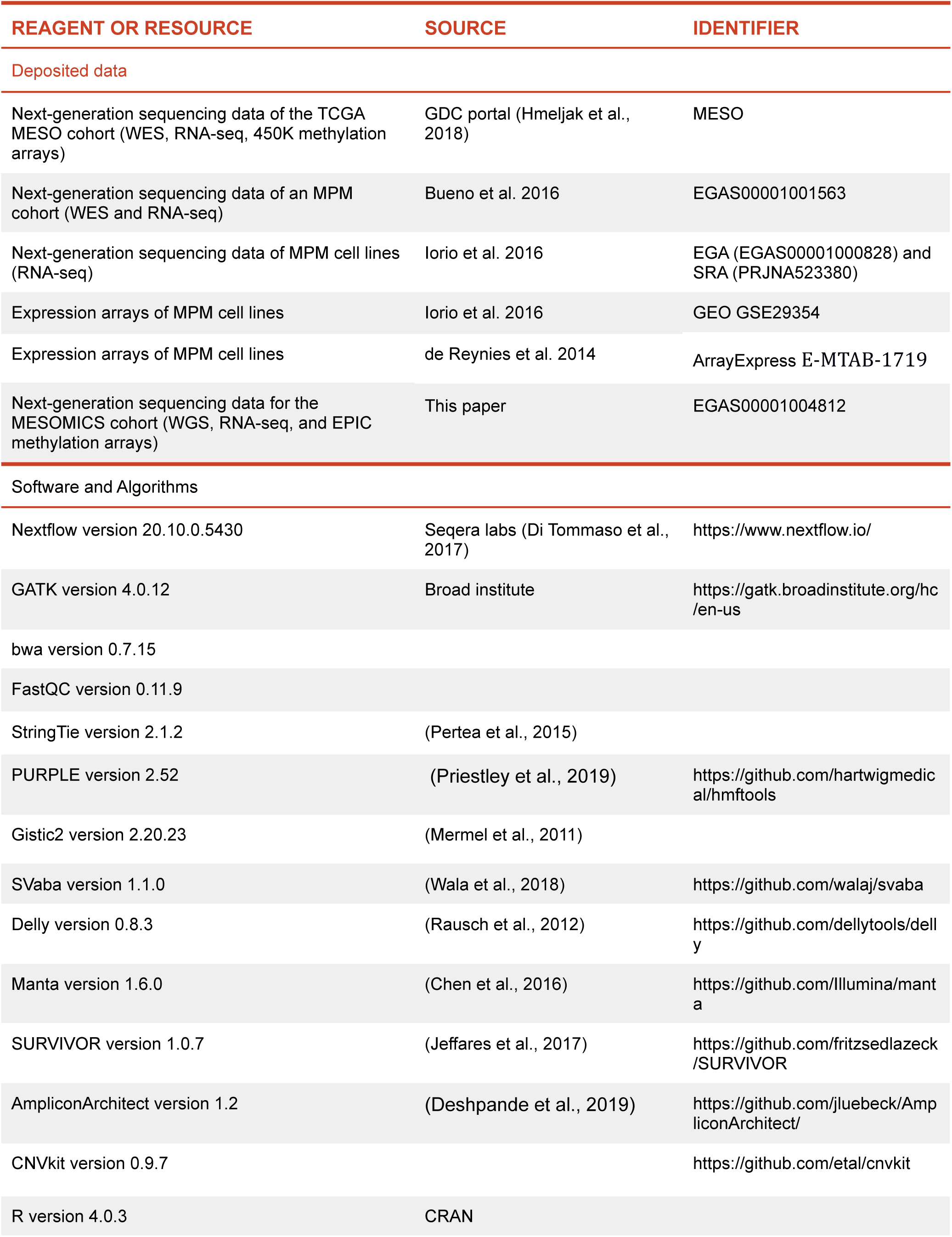
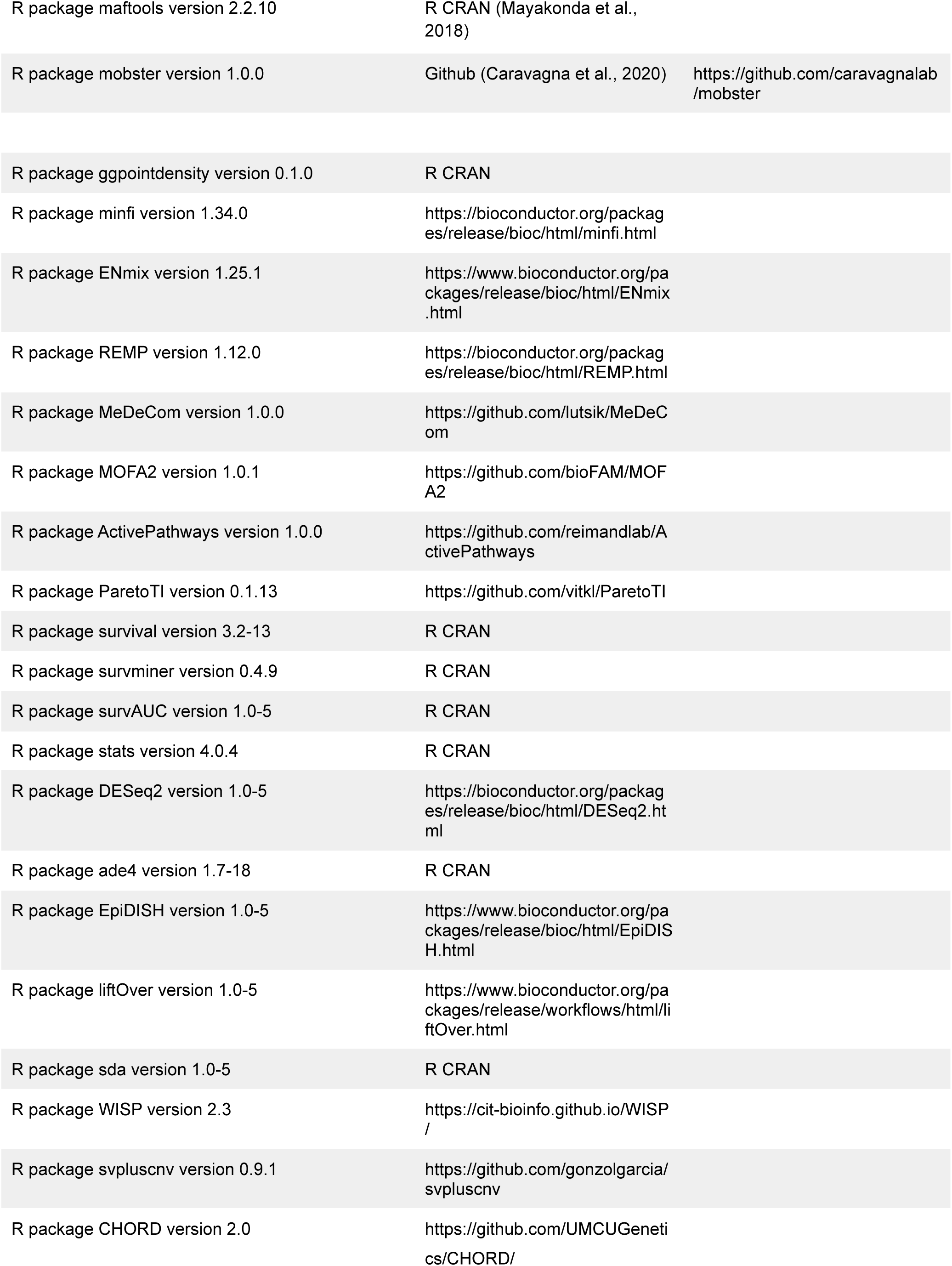

## RESOURCE AVAILABILITY

Further information and requests for resources should be directed to and will be fulfilled by Matthieu Foll.

### Data and code availability

The genome sequencing data, RNA-seq data, and methylation data have been deposited in the European Genome-phenome Archive (EGA) database, which is hosted at the EBI and the CRG, under accession number EGAS00001004812. TCGA whole-exome sequencing, RNA-seq, and methylation array data are available from the GDC portal (TCGA-MESO cohort), the whole-exome sequencing and RNA-seq data from the Bueno and colleagues cohort are available from the European Genome-phenome Archive, EGA:EGAS00001001563. Small variants lists, RNA-seq, expression array, and methylation data for the Iorio and colleagues cohort (Iorio et al., 2016) are available from the GEO (GSE29354), EGA (EGAS00001000828), and SRA (PRJNA523380) websites, and corresponding drug responses are available from the cancerrxgene.org website (accessed July 2021). Expression array data for the de Reyniès and colleagues cohort (de Reyniès et al., 2014) are available from the ArrayExpress platform (E-MTAB-1719), and corresponding drug response data from the supplementary material of Blum et al. (2019). All the other data supporting the findings of this study are available within the article and its supplementary information files.

## METHODS

### Ethics

All methods were carried out in accordance with relevant guidelines and regulations. This study is part of a larger study -the MESOMICS project-aiming at the comprehensive molecular characterization of malignant pleural mesothelioma, approved by the IARC Ethical Committee (Project No. 15-17). The samples used in this study belong to the virtual biorepository French MESOBANK, which guidelines include obtaining the informed consent from all subjects.

### Clinical data

Age at diagnosis (in years), sex (male or female), smoking status (no-smoker, ex-smoker, and smoker), asbestos exposure (exposed or non-exposed), previous treatment with chemotherapy drugs (yes or no), treatment information (surgery, chemotherapy, radiotherapy, immunotherapy, and cancer history), and survival (calculated in months from surgery to last day of follow up or death) data were collected for all the 123 patients. Median age at diagnosis was 67.5 years and 73.3% of patients were male. Detailed information from Santé Publique France (SPF), the French National Public Health Agency, about probability of exposure (no evidence found-0, possible-1/3, likely-2/3, and very likely-1), frequency (sporadic-0.25, intermittent-0.5, frequent-0.75, and permed-1), intensity (low-1, intermediate-2, high-3, and very high-4), and duration of the asbestos exposure (in years) was available for 47 patients as the result of a supervised survey, the National Program for pleural Mesothelioma Surveillance (PNSM) (soit Ilg et al., 2020). In order to compare exposure levels between patients and to reduce the number of variables, we computed a lifetime exposure score in units of years of permed low-intensity asbestos exposure, by multiplying the probability, frequency, intensity, and duration of the exposure. This score is analogous to the pack-years concept used for tobacco smoking that also balances intense, short-durations with weaker, long-duration exposures (Schaeffner et al., 2001). Indeed, 10 years of very likely, sporadic, very high intensity asbestos exposure leads to the same score (10×1×0.25×4=10) as 10 years of very likely, permed, low-intensity exposure (10×1×1×1=10) (**Table S1**). In order to improve the power of some of the statistical analyses, we regrouped some levels of the age variable which was discretized into 3 classes ((29, 63], (63, 71], and (71, 90] years).

We tested the associations between clinical variables; in particular, between a batch variable (sample provider) and the main variable of interest (histopathological type or major epithelioid subtype) or important biological covariables such as sex, age, smoking status, and asbestos exposure, using Fisher’s exact test. We found that the sample provider was not significantly associated with the clinical variables (from **Table S1**), while sex was significantly associated with smoking status and asbestos exposure (Fisher’s exact test *q*-value = 0.0002 and *q*-value = 0.03, respectively).

### The MESOMICS cohort

Samples were collected from surgically-resected tumours, applying local regulations and rules at the collecting site, and including patient consent for molecular analyses as well as collection of de-identified data. Samples underwent an independent pathological review. The MESOMICS cohort includes biological material from 120 MPM patients kindly provided by the French MESOBANK, annotated with detailed clinical, epidemiological, and morphological data. Based on the French MESOPATH reference panel, out of the 120 MPM tumor samples, 79 belong to the epithelioid type (MME), 26 are biphasics (MMB) and 15 are sarcomatoids (MMS). Out of the 105 samples with an epithelioid component (79 MME and 26 MMB), solid, acinar, trabecular, and tubulopapillary architectural patterns were the most frequent in the series (*n* = 36, 31, 17, and 14, respectively). Note that this distribution of MPM types and epithelioid subtypes does not represent the true clinical real distribution because of the bias we have introduced by including only samples with sufficient tumor content and good DNA and RNA quality. The tumor content estimated by our pathologist (FGS) in the series ranged from 10 to 100%. Similarly, the presence of infiltration was also evaluated in the H&E slides, and it ranged from 0% to 45% (**Table S1**). In addition, using the H&E slides, whole-image artificial intelligence analyses were undertaken to identify the most clinically relevant morphological features, and a score was calculated using a previously published validated algorithm (Courtiol et al., 2019).

As expected, the median overall survival (OS) for the whole series was 14 months (IQR 12-17.1), with epithelioid (MME) showing the longest OS (15.8 months, IQR 13.9-24.5) followed by biphasic (MMB; 10.8 months, IQR 6.3-17.2) and sarcomatoid (MMS; 4.5 months, IQR 2.2-NA). The tubulopapillary subtype showed the best OS (ranging from 20.9 to 41.9 months), followed by trabecular (15-18.4), acinar (13.1-15.9) and solid (11.9-14.4), which showed the worst OS. In addition, the proportion of solid subtype was negatively associated with survival (*q*-values < 0.01) (**Table S13**). The ratio of men to women is 2.76, with no statistical association between sex and histological type or subtype.

### Discovery and ITH cohorts

Among the 123 MPM patients, 13 have two tumour specimens collected to study intratumoral heterogeneity (ITH). The one with the highest tumor content, estimated by pathological review, has been selected for this descriptive study and is reported in **Table S1**, and the other region is described in **Table S4**. Additionally, three patients have been reported as non-chemonaive and they were excluded from the analyses except if explicitly mentioned otherwise in the Methods.

### Pathological review

For all 136 samples (123 tumors +13 additional regions) an H&E (hematoxylin and eosin) stain from a representative FFPE block was collected for pathological review. Our pathologist (FGS) performed a pathological review and classified all tumours according to the 2015 WHO classification (Galateau-Salle et al., 2016; 2015). The H&E stain was also used to assess the quality of the frozen material selected for molecular analyses and to confirm that all frozen samples contained at least 70% of tumour cells.

Tumour grade, immune infiltration, presence of necrosis and vessels were assessed for all 136 samples. In addition of histopathological types, we also assessed the epithelioid histopathological characteristics (architectural subtypes, cytological variants and stromal characteristics), which we subdivided into three subtypes, based on the recent IASCL-EURACAN interdisciplinary meeting recommendations (Nicholson et al., 2020): favorable prognosis (regrouping the acinar and papillary subtypes, and samples with abundant myxoid stroma), intermediate-prognosis (trabecular subtype), and unfavorable prognosis (solid subtype). Finally, we also assessed the sarcomatoid histopathological characteristics (simple, desmoplastic, low and high grade fusocellular, and with pleomorphic, heterologous, or transitional component) and both epithelioid and sarcomatoid histopathological characteristics in case of biphasic samples.

### Artificial Intelligence analysis

Whole-slide image based Artificial Intelligence (AI) prognostic score was computed using the AI MesoNet model, based on morphological features, developed by OWKIN, AI for Medical Research company (Courtiol et al., 2019). The model has been trained on a randomly selected training dataset of 2903 slides from the MESOPATH-NETMESO INCa network/MESOBANK (excluding samples from our MESOMICS cohort) and applied to the slides of our MESOMICS cohort. This model is trained to predict overall survival using only one H&E stained whole slide image per patient as input. It is therefore completely agnostic of any genomic information.

### Immunohistochemistry

Formalin fixed paraffin embedded (FFPE) tissue sections (3µm thick) from 136 MPM samples were deparaffinized and stained with the Santa Cruz BAP1 (cloneC-4) (dilution one to 50). Nuclear staining was considered positive (when nuclear expression was retained) or negative (complete loss of staining of all tumor cells with a positive internal control on the slides: fibroblast, lymphocytes and other non-tumor cells). Consequently, the positivity of BAP1 was reported as a score ranging from 0 complete loss of nuclear staining and 1 nuclear staining retained in 100% nuclei. Results are presented in **Table S1**.

### Statistical analyses

All tests involving multiple comparisons were adjusted using the Benjamini-Hochberg procedure controlling the false discovery rate using the p.adjust R function (stats package version 3.4.4).

### Survival analysis

Survival analysis has been performed using Cox’s proportional hazard model from which the significance of the hazard ratio between the reference and the other levels has been evaluated using Wald tests. We assessed the global significance of the model using the logrank test statistic (R package survival v. 2.41-3) and drew Kaplan–Meier and forest plots using R package survminer (v. 0.4.2). The proportional hazards hypothesis was checked using the Schoenfeld residuals (zph function). Univariate Cox analyses were performed for each important biological data such as sex, age, smoking status, and asbestos exposure as explanatory variable to evaluate their individual association with survival. In order to respect the minimal proportion of samples per group at 10%, we gathered current and former smoker groups together. Among the clinical data tested, only age and sex were both significantly associated with survival (Cox model *p*-value = 0.00021 and *p*-value = 0.045 respectively) (**Table S13**). As a results of univariate analyses results, in order to assess survival associations with continuous molecular variables, we fited Cox’s models by including sex and used the attained age scale, which provides a control for age effects without needing to fit an additional age parameter compatible with the proportional hazards assumption (Griffin et al., 2012) (**Table S13**).

### DNA extraction

Samples included were extracted using the Gentra Puregene tissue kit 4g (Qiagen, Hilden, Germany), following the manufacturer’s instructions. All DNA samples were quantified by the fluorometric method (Quant-iT PicoGreen dsDNA Assay, Life Technologies, CA, USA), and assessed for purity by NanoDrop (Thermo Scientific, MA, USA) 260/280 and 260/230 ratio measurements. DNA integrity of Fresh Frozen samples was checked with Tapesation system (Agilent Biotechnologies, Santa Clara, CA95051, United States) using Genomic DNA ScreenTape (Agilent Biotechnologies).

### RNA extraction

Samples included were extracted using the Allprep DNA/RNA extraction kit (Qiagen, Hilden, Germany), following manufacturer’s instructions. All RNA samples were treated with DNAse I for 15 min at 30 °C. RNA integrity of frozen samples was checked with Tapesation system (Agilent Biotechnologies, Santa Clara, CA95051, United States) using RNA ScreenTape (Agilent Biotechnologies).

Because of unsuccessful extraction (impacting either the quality or the quantity), we obtained different numbers of MPM samples for which Whole-Genome sequencing, DNA methylation, or RNA-sequencing data is available (**Table S1**).

### DNA Sequencing

#### Whole-Genome DNA Sequencing (WGS)

Whole-genome sequencing was performed by the Centre National de Recherche en Génomique Humaine (CNRGH, Institut de Biologie François Jacob, CEA, Evry, France) on 130 fresh-frozen MPMs, 54 of which with matched-normal tissue or blood samples. After a complete quality control, genomic DNA (1µg) has been used to prepare a library for whole genome sequencing, using the Illumina TruSeq DNA PCR-Free Library Preparation Kit (Illumina Inc., CA, USA), according to the manufacturer’s instructions. After quality control and normalization, qualified libraries have been sequenced on a HiSeqX5 platform from Illumina (Illumina Inc., CA, USA), as paired-end 150 bp reads. Two lanes of HiSeqX5 flow cells have been produced for each sample paired with matched-normal tissue or blood, in order to reach an average sequencing depth of 60x and one lane for the others in order to reach an average sequencing depth of 30x for the others. Sequence quality parameters have been assessed throughout the sequencing run and standard bioinformatics analysis of sequencing data was based on the Illumina pipeline to generate FASTQ files for each sample.

#### Data processing

WGS reads were mapped to the reference genome GRCh38 (with ALT and decoy contigs) using our in-house workflow (https://github.com/IARCbioinfo/alignment-nf, release v1.0), as described in (Alcala et al., 2019b). In summary, this workflow relies on the Nextflow domain-specific language (Di Tommaso et al., 2017) and consists in 4 steps: reads mapping (software bwa version 0.7.15; Li and Durbin, 2009), duplicate marking (software samblaster, version 0.1.24), reads sorting (software sambamba, version 0.6.6; Tarasov et al., 2015), and base quality score recalibration using GATK v4.0.12.

#### Variant calling and filtering on DNA

We performed somatic variant calling using software MuTect2 from GATK v4.1.5.0 (Benjamin et al.; Van der Auwera and O’Connor, 2020) as implemented in our Nextflow workflow based on the GATK best practices (https://github.com/IARCbioinfo/mutect-nf release v2.2b), using a set of 79 blood samples (16 from the MESOMICS cohort and 73 from Gabriel et al., 2021) coming from the same sequencing machine from CNRGH in Paris as panel of normal and using GATK4’s filtering module (FilterMutectCalls) with the recommended known variants VCF (gnomAD variants from GATK’s Mutect2 bundle) and per sample estimates of the contamination rate obtained with GATK4’s CalculateContamination (using the small panel of EXAC SNPs from GATK4’s Mutect2 bundle). Multi-region samples were processed jointly using the multi-sample calling mode of Mutect2. We called germline variants using Strelka2 v2.9.10-0 (Kim et al., 2018) using our Nextflow workflow (https://github.com/IARCbioinfo/mutect2-nf release v1.2a). Normalization of resulting variant calling format (VCF) files was performed with BCFtools v1.10-2 as implemented in our workflow (https://github.com/IARCbioinfo/vcf_normalization-nf v1.1), and annotation was then performed with ANNOVAR (2018Aprl16) (Wang et al., 2010) using the GENCODE v33 annotation, COSMIC v90, REVEL databases.

To call somatic variants on tumor-only samples (72/115) a similar procedure was performed (Mutect2 tumor-only mode) but including further germline filtering steps using a random forest (RF) classifier. A total of 20 features (gnomad, cosmic, genomic location/impact, and features obtained directly from Mutect2) were selected to build a RF model to classify single nucleotide variants and small Indels into somatic or germline. For training the RF model a total of 46 tumors with matched normal mesothelioma whole-genome sequences were used. Variants on this subset were called using both the tumor-only and matched modes of Mutect2. The matched somatic calls (ground-truth) were used to split the variants of the tumor-only calls into germline and somatic classes and subsampled to mitigate bias arising from class imbalance during training (1:1 somatic:germline ratio, *n*=407,984). The dataset was divided into 75% for training (*n*=305,988) and 25% for testing (*n*=101,996), and the trained model reached an accuracy of 0.93 in the test set. A random forest model for SNVs (rfvs01) was trained using a total of 326,388 (80%) variants (1:1 ratio). For indels, a random forest model (rfvi01) was built using a total of 337,442 variants (1:1 ratio, including 305,988 SVNs and 31,454 indels). To control the false positives (RF Model FDR=6.4%), given the highest expected proportion of germline variants in the prediction set, we set a cutoff (RF votes) of 0.5 and 0.75 for coding and non-coding variants, respectively. Finally, the RF models (rfvs01 and rfvi01) were used to classify a total of 1,454,942 variants (SNVs=1,317,200 and indels=137,742) of which 217,436 variants (including SNVs and indels) were classified as somatic. The point mutation calls for the MESOMIC cohort (*n*=448,434) include the matched calls (43 WGS) and the filtered tumor-only calls (72 WGS) (**Table S5**). The source code and the random forest models are available in the Github repository at https://github.com/IARCbioinfo/RF-mut-nf.

#### Copy number variant calling

Somatic Copy Number Variants (CNVs) were called using the PURPLE software (Priestley et al., 2019) , as implemented in our Nextflow workflow (https://github.com/IARCbioinfo/purple-nf, version 1.0). We used a total of 57 (including multi-region samples) matched whole-genome sequences (WGS) of MPM for benchmarking the tumor-only mode of PURPLE. We ran PURPLE twice for each matched sample: first using as input the matched WGS Normal/Tumor pair, and second, using as input only the tumor WGS. We benchmarked the PURPLE tumor-only mode by comparing the estimation of tumor purity, tumor ploidy, number of segments, percentage of genome changed (amplified, deleted), percentage of genome in neutral LOH (Loss Of Heterozygosity), and major/minor copy number alleles at gene level for matched and tumor-only PURPLE calls. CNV calls are reported in **Table S7** and presented in Figure 3B-C, and compared to that from the Bueno and TCGA cohorts in **Figure S3A**. TCGA copy number data has been downloaded from the TCGA portal (TCGA-MESO, https://portal.gdc.cancer.gov/, March 2021) corresponding to the allele-specific copy number segment data from genotyping arrays.

We observed a high concordance (pearson correlation) between tumor-only and matched PURPLE calls for tumor purity (r > 0.98), tumor ploidy (r=1), number of cnv segments per tumor (r > 0.98), percentage of genome changed (amplified, deleted, r > 0.99), and percentage of genome in neutral LOH (r > 0.99). Moreover, the concordance between tumor-only and matched PURPLE calls was also high at gene level with major and minor copy number alleles reaching R > 0.96. However, we observed artifactual focal amplifications and deletions near telomeric and centromeric regions that were not called when using the matched data. These regions were identified and the segments overlapping these regions were removed from the tumor-only calls. The copy number calls for the MESOMIC cohort (115 WGS) include the matched PURPLE calls (43 WGS) and the filtered tumor-only PURPLE calls (72 WGS). Whole genome doubling samples were called in genomes with more than 10 autosomes with major allele copy number > 1.5. Near haploid samples were identified as those with LOH genome percentage larger than 80%. Finally, recurrent genomic regions of DNA copy-number alterations in the 115 WGS were identified with GISTIC2.0 (Mermel et al., 2011, version 2.20.23, -conf 99%) using as input the PURPLE CNV calls (log2(totalcopynumber)-1) (**Table S8**).

For replication of the analyses using the whole-exome sequencing data from the TCGA and Bueno cohorts (**Figure S3A**), because PURPLE is only suited for WGS data, we used software Facets (Shen and Seshan 2016) instead, as implemented in our pipeline (https://github.com/IARCbioinfo/facets-nf v. 2.0).

#### Structural variant calling

To identify somatic structural variants (SVs), including insertions, deletions, duplications, inversions, and translocations, we built a consensus SVs call set by integrating SvABA (v1.1.0, Wala et al., 2018), Manta (v1.6.0, Chen et al., 2016), and Delly (v0.8.3, Rausch et al., 2012) calls with SURVIVOR (v1.0.7, Jeffares et al., 2017). For matched WGS, Delly was run in somatic SV discovery mode using the hg38 blacklisted Delly regions (“-x human.hg38.excl.tsv”; the list excludes centromere, telomere, and heterochromatin regions, as well as alt, decoy, and unknown contigs of hg38) and the tumor/normal WGS pairs. A list of somatic SVs passing all filters was generated using the Delly somatic filter, which considers somatic SVs as those with at least 10 fold coverage in the tumor sample and without evidence of normal read support for the alternative allele (ALT support in normal equal 0). Manta was run in somatic SV discovery mode using the tumor/normal WGS (--normalBAM and –tumorBAM options) and excluding the non-chromosome contig sequences (alt and decoy) of hg38 (--callRegions option). The somatic SVs passing all Manta filters (minPassSomaticScore >=30) were considered for the consensus step. SvABA was run using our in-house Nextflow workflow (https://github.com/IARCbioinfo/svaba-nf, revision number 1.0) to identify somatic and germline SVs using the tumor/normal WGS. The somatic SVs passing all SvABA filters were considered for the consensus step. The overlap of filtered somatic SV calls was performed using SURVIVOR (merge subcommand) considering as matching SV breakpoints those at a maximum distance of 1kb (ignoring SV type and SV strand). Somatic SVs (minimum SV size 50bp) identified by at least two callers and single caller predictions with a minimum read support of 15 pairs (including pair-end and split-read evidence) were included in the consensus set of each matched sample.

To filter germline SVs in tumor-only samples we trained a random forest model for each SV caller. The SV random forest model includes a total of 19 features, which are associated with external SV databases, custom panel-of-normal SVs, genomic regions, and SV features derived from SV callers. The training of the random forest model for each SV caller was performed using the matched WGS (57 including multi-region samples). First, the somatic calls from the matched WGS were used as the ground-truth during training and evaluation of each SV random forest model. Second, tumor-only calls were generated for the matched data using the tumor WGS for Manta and Delly. For SvABA, the somatic and germline SVs called by the somatic mode were merged to generate the tumor-only calls from the matched data. Third, the panel of normals for each matched WGS and SV caller combinations were generated by integrating 45 germline SV calls (excluding the respective normal sample) with SURVIVOR (merge command). Fourth, a total of 12,454, 16,720, and 12,264 SVs at 1:1 somatic:germline proportions were used to train (75%) and evaluate (25%) the random forest models of Delly, Manta, and SvABA, respectively.. The accuracy achieved on the benchmark set was 0.9, 0.89 and 0.88 for Delly, SvABA, and Manta SV RF models, respectively. Finally, the SV random forest models were used to filter the germline SVs from tumor-only samples using a cutoff (RF votes) of 0.5 and 0.75 for coding and non-coding SVs, respectively. SVs matching one present in the custom PON or located in centromeric regions were discarded. SV call set for each tumor-only sample was created using the same steps performed for the matched WGS (merging Delly, SVaba and Manta calls with SURVIVOR and keeping single caller predictions with read support >= 15). Moreover, SVs found in more than 4 samples in the tumor-only series were also classified as potentially germline and removed from the final consensus set. The SVs calls for the MESOMIC cohort (**Table S6**, *n*=12,914) include the matched SV calls (43 WGS, *n*=4,685) and the filtered tumor-only SV calls (72 WGS, *n*=8,229). The source code and the SV random forest models are available in the Github repository at https://github.com/IARCbioinfo/ssvht.

#### Damaging variants and driver detection

Mutational cancer driver genes have been detected using the state-of-the-art integrative oncogenomics pipeline (IntOGen; Martínez-Jiménez et al., 2020), that distinguishes signals of positive selection from neutral mutagenesis across a cohort of tumors by combining multiple driver detection methods. The IntOGen pipeline was run for each cohort separately, and also for the combined cohort to gain in statistical power, and to detect mutational driver genes that may be specific to each of them (**Figure S4F** and Figure 4, left panel). Of note, variants occurring on mitochondria chromosome chrM have not been considered in this analysis. Genes that drive tumorigenesis upon SVs have simply been selected based on their recurrence, using a cutoff of 5 samples (**Figure S4C**).

The damaging SNVs, indels and structural variants have been selected as follows. First, for SNVs and small indels, we used ANNOVAR annotations to restrict the list to the exonic or splicing, non-synonymous variants. For multi-nucleotide polymorphisms (MNP), we used the Coding Change ANNOVAR procedure to infer the protein changes occurring and in case of any amino acid changes, we classified the event as damaging. Finally, we removed structural variants for which the breakpoints lead to harmless changes for the coding sequence of the gene such as large in frame deletion in a single intron.

#### Comparison with PCAWG and TCGA

Tumor mutational Burden comparison of Mesothelioma and TCGA cohorts (**Figure S4A**) was performed with the mafftool (Mayakonda et al., 2018) package (v2.6.05). The PCAWG data for mRNA fusions (version 1.0), SVs (version consensus_1.6.161116), CNVs (consensus.20170119), and number of SNVs represented in **Figure S4B** were downloaded from the PCAWG site (https://dcc.icgc.org/releases/PCAWG).

#### Identification of complex mutational process in MPM tumors

Mutational SBS signatures were de novo discovered and decomposed into COSMIC mutational signatures with the SigProfilerExtractor (Alexandrov et al., 2020) tool. The SNVs called by both Mutect2 and Strelka (Kim et al., 2018, nextflow workflow https://github.com/IARCbioinfo/strelka2-nf v1.2a) on the T/N samples were used as input for SigProfilerExtractor (v1.0.17) to avoid caller specific signatures. Copy Number signatures were called using SigProfilerExtractor as described in (Steele et al., 2021) (**Table S7**) and using as input the PURPLE copy number segments. SV signatures were also called using the SigProfiler framework but using a newer version of SigProfilerExtractor (v1.1). Finally, detection and classification of clustered mutations (kataegis analysis) was performed as described in (Bergstrom et al., 2021). The list of clustered mutations per tumor including their classes are provided in **Table S9**, and represented in **Figure S3C**.

Chromothripsis regions were identified by combining SVs and CNV calls with the svpluscnv R package (Lopez et al., 2021). To identify shattered regions’ breakpoints from CNVs and SVs, breakpoints were counted by splitting the genome into 10Mb windows. Contiguous windows with a high density of breakpoints were merged into larger shattered regions. Then interleaved SVs and variations in copy number state signatures were used to differentiate chromothripsis from focal events such as double minutes. Additionally, following recent practices (Cortés-Ciriano et al., 2020), we classified the shattered region into high and low confidence by considering the number of oscillating CN segments: high-confidence calls were classified as those displaying an oscillation pattern between two copy number states in at least seven adjacent CN segments, others were classified as low-confidence calls (**Table S9**, **Figure S3D**).

Amplicon predictions were performed using the AmpliconArchitect program version 1.2 (Deshpande et al., 2019). In Brief, the copy number variants were called using the CNVkit program (version 0.9.7), which is the recommended CNV caller to identify seed for AmpliconArchitect. Seed selection was performed following the recommended criteria (minimum segment length of 50Kb and minimum copy number gain of 4.5) using the amplified_intervals.py (amplified_intervals.py --gain 4.5 --cnsize_min 50000 --ref GRCh38) script provided by the AmpliconArchitect package. AmpliconArchitect (Version 1.2) was then run with default parameters using the selected seeds and the tumor CRAM files as input. Finally, the AmpliconClassifier program was run to classify the amplicons generated by AmpliconArchitect into ecDNA, BFS, Complex, linear or non-amplified classes (**Table S9**, **Figure S3B**). Finally, the homologous recombination deficiency samples were identified using the R package CHORD (Nguyen et al., 2020) version 2.0 (**Table S9**). Following CHORD recommendation, four HRD positive samples were marked with a not determined HRD type because they have less than 30 SV.

#### TERT promoter mutation analyses

Point mutations within the *TERT* promoter region (chr5: 1,294,956-1,295,406, hg38) were identified from the VCF file outputs of WGS prior to filtering T-only variants using the random forest filter (see Variant calling and filtering on DNA). Pre-filtered VCF files were used due to low mappability of the region that results in high false negative point mutation detection rates. Genomic coordinates were selected specifically as all previously reported *TERT* promoter mutations in mesothelioma (C158A, A161C, C228T, C250T) are contained within the above region (Pirker et al., 2020; Quetel et al., 2020). Three of four reported mutations were identified in seven samples: A161C (chr5: 1,295,046 in hg38 coordinates), C228T (chr5: 1,295,113), and C250T (chr5: 1,295,135). Results are presented in **Figure S5A**.

### RNA Sequencing

#### RNA Sequencing (RNA-seq)

RNA sequencing was performed on 126 fresh frozen MPM in the Cologne Centre for Genomics, of which 109 MPM belonged to the discovery cohort (**Table S1**). In addition, we collected two technical replicates, MESO_051_TR and MESO_115_TR, from two different patients and coming from the same RNA extraction as MESO_051_T and MESO_115_T respectively but sequenced separately. Libraries were prepared using the Illumina® TruSeq® mRNA stranded sample preparation Kit. Library preparation started with 1 µg total RNA. After poly-A selection (using poly-T oligo-attached magnetic beads), mRNA was purified and fragmented using divalent cations under elevated temperature. The RNA fragments underwent reverse transcription using random primers. This is followed by second strand complementary DNA (cDNA) synthesis with DNA Polymerase I and RNase H. After end repair and A-tailing, indexing adapters were ligated. The products were then purified and amplified (14 PCR cycles) to create the final cDNA libraries. After library validation and quantification (Agilent 4200 Tapestation), equimolar amounts of library were pooled. The pool was quantified by using the Peqlab KAPA Library Quantification Kit and the Applied Biosystems 7900HT Sequence Detection System. The pool was sequenced by using an Illumina Novaseq 6000 sequencing device and a paired end 100nt protocol.

#### Data processing

The 126 raw reads files from the MESOMICS cohort and the 21 files from the Iorio and colleagues (2016) mesothelioma cohort (downloaded from the EGA and SRA websites, datasets EGAS00001000828 and PRJNA523380) were processed in three steps using the RNA-seq processing workflow based on the Nextflow language and accessible at https://github.com/IARCbioinfo/RNAseq-nf (release v2.3), as described first in Alcala, Leblay, Gabriel *et al*. (2019b). In summary, reads were trimmed for the adapter sequence using Trim Galore v0.4.2, then mapped to reference genome GRCh38 (using annotation gencode version 33) with STAR software (v2.7.3a). Then, reads were realigned locally using ABRA2 (Mose et al., 2019; workflow https://github.com/IARCbioinfo/abra-nf release v3.0), and base quality scores were recalibrated using GATK (workflow https://github.com/IARCbioinfo/BQSR-nf release v1.1). Once processed, expression was quantified for each sample, generating a raw read count table with gene-level quantification for each gene of the comprehensive gencode gene annotation file (release 33), as well as a table with Gene fragments per kilobase million (FPKM), using StringTie software (v2.1.2) (Nextflow pipeline accessible at https://github.com/IARCbioinfo/RNAseq-transcript-nf release v2.2). Quality control of the samples was performed at each step. FastQC software (v0.11.9; https://www.bioinformatics.babraham.ac.uk/projects/fastqc/) was used to check raw reads quality and RSeQC software (v3.0.1) was used to check alignment quality

#### Normalisation and quality controls

The raw read counts of the 59,607 genes in the expression data matrix, from MESOMICS, TCGA, and Bueno cohorts, from which we removed non-chimionaif samples, were normalised using the variance stabilisation transform (vst function from R package DESeq2 v1.14.1); this transformation enables comparisons between samples with different library sizes and different variances in expression across genes. We performed dimensional reduction on expression data as quality control, using Principal Component Analysis (PCA) (function dudi.pca from R package ade4 v1.7-16). PCA was performed on the variance-stabilised read counts of the 5,000 most variable genes for (i) (MESOMICS) 109, (ii) (Bueno) 180, (TCGA) 73, and (iii) (3-cohorts: MESOMICS, Bueno, and TCGA) 362 samples (**Table S3**). For each set, samples were plotted by their coordinates to visualise outliers. For each dataset, linear regression analysis was performed to determine any significant association between these PCs and technical variables such as RNA-seq batch, macrodissection and provider. We found no outliers, and no batch effect in this data.

#### Variant calling and filtering on RNA

We used Mutect2 with the --allele flag to force genotyping of variants identified by mutect in the whole-genome sequencing data to call variants on the 126 RNA sequencing data for validation (workflow https://github.com/IARCbioinfo/mutect2-nf release v2.2b with option --genotype).

#### Fusion transcript analysis

Fusion transcripts were detected using Arriba (Uhrig et al., 2021) for the MESOMICS, Bueno, and TCGA cohorts. First, RNA-seq reads were aligned using STAR (2.7.6a) to the hg38 reference. Second, Arriba was used to call mRNA-fusions using the STAR alignment (BAM) and Arriba blacklisted regions (-b option). For the MESOMICS cohort, we additionally integrated the genomic SVs by including the SV breakpoints into the calling (-d option). Finally, high-quality mRNA-fusion predictions for all MPM cohorts were defined as those Arriba predictions classified as high confidence and with a minimum support of 10 reads from paired-end and split-read alignments. The **Table S6** contains all the mRNA fusions passing the aforementioned filters, and **Figure S4E** represents recurrently altered genes.

#### Processing of publicly available expression array data

Raw expression array CEL files from Iorio and colleagues (Iorio et al., 2016) and de Reynies and colleagues (de Reyniès et al., 2014) were downloaded from public repositories (GEO: GSE29354 and ArrayExpress: E-MTAB-1719, respectively) and processed using the RMA algorithm (justRMA function from the affy R package v1.68.0). Annotations were downloaded from the hgu219.db and hgu133plus2.db packages (v3.2.3).

#### Immune contexture deconvolution from expression data

The proportion of cells that belong to each of ten immune cell types (B cells, macrophages M1, macrophages M2, monocytes, neutrophils, NK cells, CD4+ T cells, CD8+ T cells, CD4+ regulatory T cells, and dendritic cells) were estimated from the RNA-seq data using software quanTIseq (downloaded 14 September 2020) using our workflow for parallel processing of samples (https://github.com/IARCbioinfo/quantiseq-nf release v1.1). Additionally, as technical validation, we used EpiDISH R package (v2.6.0) to estimate seven immune cell types (B cells, monocytes, neutrophils, NK cells, CD4+ T cells, CD8+ T cells, and eosinophils) as well as epithelial cells and fibroblasts from the DNA methylation data. The immune cell types for which the association with archetypes were the strongest (absolute Pearson’s correlation coefficient r > 0.4, B cells, CD8+ T, and neutrophils) presented significant concordance between softwares (additionally to monocytes). The other estimates (NK cells and CD4+ T) have not been confirmed in EpiDISH estimation, possibly because of the reference differences —such as the reference size, the number of cell types estimated— between softwares. Proportion of cells in the TCGA and Bueno samples were taken from the supplementary tables of Alcala et al. (Alcala et al., 2019a), which used the exact same software and version.

#### WGD expression analyses

To identify significant differentially expressed genes associated with WGD status, we employed the same strategy introduced by (Quinton et al., 2021). In brief, the expression of each gene (TPM values) was modeled as a function of WGD + purity + local_copy_Number. The purity, local_copy_number (log2(total copy number)), and WGD status were obtained from PURPLE predictions. Genes were considered significantly associated with WGD status if they had an FDR q-value of less than 0.05. Pathway enrichment analyses were performed with the hypeR (Federico and Monti, 2020) package (v1.9.0) using the MSigDB Hallmark gene sets (v7.4.1) and the list of differentially expressed WGD genes. Pathways with an FDR q-value of less than 0.05 were considered significantly associated with WGD status. Results are reported in **Table S10** and presented in Figures 5B **and S5C**.

### DNA methylation

#### EPIC 850k methylation array

Epigenome analysis was performed on 119 MPMs (**Figure S1A, Table S1**), two technical replicates and three adjacent normal tissues. Epigenomic studies were performed at the International Agency for Research on Cancer (IARC) with the Infinium EPIC DNA methylation beadchip platform (Illumina) used for the interrogation of over 850,000 CpG sites (dinucleotides that are the main target for methylation). Each chip holds eight samples, and the 140 samples were spread over 19 chips. We used stratified randomisation to mitigate the batch effects, samples were arranged over the chips to evenly distribute, in order of priority, histopathological type, major epithelioid subtype, provider, sex, smoking status, age and professional asbestos exposure. However, due to differences in the number of each histopathological type, and date of sample arrival, four of the 19 chips contained exclusively one type. Technical replicates were placed on different chips, whilst ITH and adjacent normal samples were placed on the same chip as their corresponding tumour sample. The position of samples on each chip was then randomised.

For each sample, 600 ng of purified DNA were bisulfite converted using the EZ DNA Methylation kit (Zymo Research Corp., CA, USA) following the manufacturer’s recommendations for Infinium assays. Then, 250 ng of bisulfite-converted DNA was used for amplification, fragmentation and finally hybridisation on Infinium Methylation EPIC beadchip, following the manufacturer’s protocol (Illumina Inc.). Chips were scanned using Illumina iScan to produce two-colour raw data files (IDAT format).

#### Data processing

The resulting IDAT raw data files were pre-processed using R packages minfi (v. 1.34.0) and ENmix (v. 1.25.1). We first performed quality control checks on the raw data. Two-colour intensity data of internal control probes were manually inspected to check the quality of successive sample preparation steps (bisulfite conversion, hybridisation, extension, and staining; function plotQC, ENmix). There was one outlier, the technical replicate MESO_056_T1, when comparing per sample log2 methylated and unmethylated chip-wise median signal intensity (function getQC, minfi), and no samples displayed an overall *p*-detection value > 0.01 (function detectionP, minfi). The poor quality sample, MESO_056_T1, was excluded from subsequent processing. Sex was assigned using a predictor based on the median total signal intensity of sex chromosomes, with a cutoff of -2 log2 estimated copy number difference between males and females (function getSex, minfi). One sample was identified to be discordant between predicted (female) and clinically reported (male) sex, MESO_071_T. Whole genome sequencing results from matched blood confirmed that the participant was male, whilst the tumour displayed losses on chrY and gains on chrX.

Raw data were then normalised using functional normalisation (function preprocessFunnorm, minfi) to reduce technical variation within the data, and probe removal steps were performed to ensure reliability and accuracy of the final dataset. Four groups of probes were removed: (i) poor performing probes with a *p*-detection value > 0.01 in at least one sample (16,497 probes discarded), *p*-detection value was computed by comparing the total signal (methylated and unmethylated) of each probe with the background signal level from non-negative control probes (function detectionP, minfi) (ii) cross-reactive probes (42,552 probes discarded), cross-reactive probes co-hybridise to multiple locations within the genome and therefore cannot be reliably investigated (Pidsley et al. Genome Biology 2016 PMID: 27717381) (iii) probes on the sex chromosomes (17,144 probes discarded), and (iv) probes with SNPs within the single base extension site, or target CpG site, at a minor allele frequency of > 5% (database dbSNP build 137), (8,411 probes discarded, function dropLociWithSnps, minfi). This resulted in a normalised, filtered dataset of 781,245 probes for 139 samples. Finally, beta and M-values were extracted (functions getBeta and getM, minfi). Nine probes recorded m-values of -∞ for at least one sample, and these values were replaced with the next lowest m-value in the dataset. The three normal tissues and one remaining technical replicate were then removed from the beta and m matrices for the subsequent analyses. This resulted in 135 samples, 122 for discovery and an additional 13 for ITH analyses.

#### Processing of publicly available DNA methylation data

DNA methylation array data (IlluminaHumanMethylation450k BeadChip array IDAT files) from the TCGA mesothelioma cohort (Hmeljak et al., 2018) were downloaded from the GDC legacy archive (https://portal.gdc.cancer.gov/legacy-archive/search/), and from the Iorio et al. (2016) cell line cohort from GEO repository (dataset GSE68379), respectively. Datasets were then imported into R and pre-processed using R packages minfi (v. 1.34.0) and ENmix (v. 1.25.1) individually. Data processing was performed as per the MESOMICS cohort, no samples failed QC steps or were discordant for sex. Probes with *p*-detection value > 0.01, cross-reactive probes (Chen et al., 2013), probes on sex chromosomes, and those associated with SNPs were discarded. This resulted in normalised, filtered datasets of 439,417 probes for 74 samples for the TCGA cohort, and 436,125 probes for 21 samples for the Iorio cell line cohorts. Beta and M-values were extracted (functions getBeta and getM, minfi), and probes recording M-values of -∞ for at least one sample were replaced with the next lowest m-value in the dataset.

Where DNA methylation array data was required for the MESOMICS and TCGA cohorts together (see ***Integrative unsupervised analyses***), data were combined and processed as follows. IDAT files for 126 MESOMICS samples (excluding ITH samples), and 74 TCGA samples were imported into R as separate RGSets. The TCGA RGSet was converted to array type IlluminaHumanMethylationEPIC and combined with the MESOMICS RGSet (function convertArray and combineArrays from R package minfi). All samples passed QC. One sample was identified to be discordant between predicted and clinically reported sex, MESO_071_T, as previously described. Subsequent processing was as per the MESOMICS cohort, and 56,308 probes were discarded (16,588 with *p*-detection value > 0.01, 26,254 cross-reactive, 8,838 sex chromosome, and 4,628 SNP-associated). This resulted in a normalised, filtered dataset of 396,145 probes for 200 samples. Beta and M-values were extracted (functions getBeta and getM, minfi), one hundred and twenty eight probes recorded m-values of -∞ for at least one sample and were replaced with the next lowest M-value in the dataset. Seven samples were then removed from the beta and m matrices (all MESOMICS samples), three normal tissues, one technical replicate and three non-chimionaif samples, resulting in a dataset of 193 samples.

#### Global methylation level

DNA methylation level at LINE1 repetitive elements was used as an estimate of global methylation level. Methylation level at LINE1 repetitive elements were calculated using the REMP package (v 1.12.0) functions to extract m and beta values of CpGs that are located in LINE1 (Zheng et al., 2017). REMP functions were performed on the normalised, filtered *M* table containing 781,245 probes, and identified 23,906 probes located in LINE1 elements. Average *M* and beta values were then calculated for each individual sample across all LINE1 probes respectively to obtain mean LINE1 methylation levels per sample. The mean *M* values were used for statistical analysis of associations between global methylation levels and features of interest (**Table S2**), while beta values were used for plotting significant findings. An examination of the mean methylation level across LINE1 probes identified one outlier, MESO_040_T, for which the global level of methylation appears particularly low in comparison to the rest of the cohort, nevertheless this single sample only marginally influenced the relationship between LINE1 and other variables mentioned in the main text.

#### Regional methylation analysis

Methylation profile within promoter, enhancer and gene body regions were examined as follows. Array probes were classified as promoter, enhancer, gene body or other, using annotations provided in the EPIC 850K array manifest b5 (version 1.0 b5, downloaded from: https://emea.support.illumina.com/downloads/infinium-methylationepic-v1-0-product-files.html).

Probes with a value of “Promoter_Associated” in the column ‘Regulatory_Feature_Group’ were assigned as promoter probes, those with any value in the column ‘Phantom5_Enhancers’ were assigned as enhancer probes, and probes with a value including “Body” or “1stExon” in the column ‘UCSC_RefGene_Group’ were assigned as gene body probes. Probes which fell into multiple groups were classified as promoter first, if applicable, then as enhancer probes. The dataset of 781,245 probes contained 102,341 promoter-assigned probes, 23,858 enhancer-assigned probes, and 317,281 gene body assigned probes.

Average M and beta values were calculated for each individual sample across all promoter, enhancer and gene body probes to obtain mean promoter, enhancer and gene body methylation levels per sample respectively. The mean M-values were used for statistical analysis of associations between regional methylation levels and features of interest, while beta values were used for plotting significant findings.

#### Deconvolution of enhancer methylation profile

Deconvolution of enhancer methylation levels was performed with non-negative matrix factorisation using R package MeDeCom (v1.0.0) (Lutsik et al., 2017). The 5,000 most variable enhancer probes (variance calculated from beta values) were input to identify latent methylation components (LMC, cell-type specific methylation profiles). Values of *k* = 3 and *λ* = 0.01 were selected by examining the resulting cross-validation error plot; LMC values are reported in **Table S2**. Attributing the three resulting latent methylation components to cell types was performed through Pearson correlation tests of proportion of each LMC present in a sample against the proportion of individual cell types within each sample (**Figure S1H-J**). Proportions of B cells, M1 and M2 macrophages, monocytes, neutrophils, NK cells, T-CD8+, T-CD4+, T regulatory cells and dendritic cells were estimated from the result of quanTIseq analysis of RNA sequencing data (**Figure S1H**), see ***Immune contexture deconvolution from expression data***. Proportions of sarcomatoid and epithelioid cell types were estimated by histopathological review by the study pathologist FGS, see ***Pathological review* (Table S1)**.

#### CpG island methylator phenotype index

A CpG island methylator phenotype (CIMP) index value was calculated for all samples as follows. Probes located within Cpg islands (denoted as “Island” in the Epic 850k array manifest b5 column *Relation_to_UCSC_CpG_Island*) were retained, the mean beta value across all probes within each island (identified from manifest column *Island_name*) was calculated per sample resulting in beta values for 24,891 and 24,924 CpG islands, MESOMICS (EPIC array), TCGA (Hmeljak et al., 2018), and Iorio and colleagues (Iorio et al., 2016) cell lines (HM450K array), respectively. The CIMP index was then calculated as the proportion of these 24,891 or 24,924 islands with ≥ 30% methylation (beta value ≥ 0.3) per sample. CIMP index values ranged from 0.32 to 0.56, meaning 32% to 56% of all islands represented on the array were considered methylated per sample. A proxy for this CIMP index was computed based on the mean methylation level of promoter CpG islands for five genes only: *CACNA1G* (island coordinates (hg19): chr17:48636103-48639279), *IGF2* (chr11:2158951-2162484), *NEUROG1* (chr5:134870740-134872051), *RUNX3* (chr1:25255527-25259005), and *SOCS1* (chr16:11348541-11350803) (selected from Weisenberger et al., 2006, **Table S2**).

A previously published method for calculating CIMP index that was also tested (in the MESOMICS cohort only), here called CIMP-normal index, as follows (Blum et al., 2019). Probes located within CpG islands were retained, the mean beta value across all probes within each island was calculated for the three adjacent normal tissues available in the MESOMICS cohort. Islands whose methylation level was < 30% in all three adjacent normal samples were retained (*n* = 15,824), denoted as normally hypomethylated islands. The CIMP-normal index was then calculated as the proportion of these 15,824 islands with ≥ 30% methylation (beta value ≥ 0.3) per sample. CIMP-normal index values ranged from 0.13 to 0.19, corresponding to 0.13% to 19% of normally hypomethylated islands to be hypermethylated per sample (**Table S2**). There was a significant correlation between the two CIMP index values calculated (*p*-value = 3.27e-66, *r* = 0.96). The method for CIMP-normal index was based on first identifying normally hypomethylated islands, therefore requiring normal pleura or mesothelium. The normal tissues available in the MESOMICS cohort are adjacent to mesothelioma samples, therefore that they are unlikely to be pure non-tumour tissues, as such, the CIMP index rather than the CIMP-normal index was used for subsequent analysis.

#### Annotating IlluminaHumanMethylationEPIC array probes with gene ID

Probes were assigned to a gene based on the contents of the EPIC 850K array manifest b5 column ‘UCSC_RefGene_Name’. Additionally, promoter and enhancer only associated probes which did not have any gene annotation in the manifest column ‘UCSC_RefGene_Name’ were then assigned a ‘nearest gene’ annotation using the function matchGenes with the TxDb.Hsapiens.UCSC.hg19.knownGene library from R package bumphunter.

#### Correlation between methylation and expression

Correlation between methylation levels and gene expression was performed as follows. Regional level testing: probes were divided into promoter, enhancer and gene body (see ***Regional methylation analysis***), probe groups were then filtered to retain only those with a difference of > 0.1 beta value between lowest and highest methylation level across 119 samples (samples input to MOFA analysis i). Pearson correlation tests were performed between the m-value of all probes within a region group and their corresponding gene expression level (normalised using variance stabilisation transformation, filtered for genes having > 1 FPKM difference across 109 samples). This resulted in testing within 109 samples with both methylation and expression data of 37,067 promoter probes against expression of 8,444 genes, 20,308 enhancer probes against expression of 6,539 genes, and 262,820 gene body probes against expression 15,825 genes. *p*-values were adjusted for multiple testing using Benjamini-Hochberg method within region groups, probes were considered correlated with expression at *q*-value ≤ 0.05.

Island level testing: probes located within Cpg islands (denoted as “Island” in the Epic 850k array manifest b5 column *Relation_to_UCSC_CpG_Island*) were retained, the mean M-value across all probes within each island (identified from manifest column *Island_name*) was calculated per sample resulting in M-values for 24,891 CpG islands. Pearson correlation tests were performed between the M-value of each island and their corresponding gene expression level (normalised using variance stabilisation transformation, filtered for genes having > 1 FPKM difference across 109 samples). Corresponding genes for each island were identified as the corresponding gene for each probe within the island (see *Annotating IlluminaHumanMethylationEPIC array probes with gene ID*). This resulted in testing within 109 samples with both methylation and expression data of 21,189 islands against expression of 12,992 genes.

### Epithelial-mesenchymal transition methylation quantification

#### Epithelial-mesenchymal transition expression score

A score of epithelial-mesenchymal transition (EMT) per sample was calculated from variance-stabilized read counts as the mean expression of 52 mesenchymal-associated genes minus the mean expression of 25 epithelial-associated genes, as previously described (Hmeljak et al., 2018; Mak et al., 2016). A higher EMT score indicates a more mesenchymal-like gene expression profile than epithelial-like. Results are reported in **Table S2**.

#### Methylation

EMT gene methylation levels were calculated as follows. Firstly, all probes within promoter, enhancer, or gene body groups associated with at least one of the panel of 77 EMT-associated genes (Mak et al., 2016) in the manifest column ‘UCSC_RefGene_Name’ or ‘nearest gene’ annotation (function matchGenes, bumphunter) were selected. This resulted in 3,764 probes across all 77 genes, specifically 150 promoter probes corresponding to 17 EMT genes, 207 enhancer probes corresponding to 54 EMT genes, and 2,446 body probes corresponding to 76 EMT genes. The mean *M*- and beta-values across all epithelial and mesenchymal genes separately for each region group were then calculated per sample.

### Epithelial (E) and Sarcomatoid (S) scores

For each sample, E- and S-scores were computed for the MESOMICS, TCGA, Bueno and cell-lines samples using expression data (normalized read count for MESOMICS, TCGA, Bueno and expression array data for cell-lines) and the method WISP from Blum et al. (2019). The method relies en unsupervised clustering to identify three clusters, enriched for samples of the Epithelioid, Sarcomatoid histopathological types and Normal cells, respectively, and then uses these samples to produce signature expression profiles that are used to perform a deconvolution of all the samples using a constrained linear model. Results are presented in **Table S2, S13, and S14**.

### Genomic instability scores

We estimated genomic instability from all omic’s layers: genomic, expression, and methylation profiles. From the genome, we calculated the proportion of changes in the genome in terms of copy number. From expression data, we computed a hallmark score using hallmarks of cancer (Keifer et al. 2017) by summing the normalized read count of the genes belonging to each hallmark. Finally, we used global methylation level (see ***Global methylation level*** section) as a third proxy of genomic instability. Values are reported in **Table S2**.

### Integrative unsupervised analyses

We performed four series of analyses, with different subsets of samples: (**i**) discovery analyses with all our discovery cohort (MESOMICS cohort, 120 samples) for which WGS, RNA-seq, and/or 850 K methylation array data are available, (**ii**) and (**iii**) replication analyses with the already published data from Bueno (2016) (Bueno cohort, 181 samples after exclusion of non-chemonaive samples) and Hmeljak and colleagues respectively (2018) (TCGA cohort, 73 samples in the curated list), (**iv**) combined analyses integrating the MESOMICS, Bueno, and TCGA cohorts with a total of 374 samples, (**v**) replication combining cell lines from the Iorio (2016)—for which whole-exome sequencing, expression arrays and RNA-seq, 450K methylation arrays, and drug responses in the form of IC_50_ scores are available—(21 samples, 265 drugs) and the de Reyniès (2014) and Blum et al. (2019) datasets—for which expression arrays and drug responses are available—(38 samples, 3 drugs). In addition, some single-omic analyses are also described in this section.

#### Pre-processing of expression data

We used normalised read counts matrices (see ***RNA Sequencing***) for (**i**), (**ii**),(**iii**) and (**iv**) encompassing 59,607 genes. Among these genes, those having less than one FPKM difference across the samples have been excluded from the unsupervised analyses. Also, in order to mitigate sex influence on the expression profiles, we removed genes from the sex-chromosomes. For each analysis, the top 5,000 most variable genes were selected. Similarly, the 5,000 most variable genes from the normalized array expression of cell lines (see *expression array processing* section) were selected. Whenever several probes were available for a same gene, the one with the highest intensity was selected.

#### Pre-processing of methylation data

DNA methylation was available for both MESOMICS and TCGA cohorts. Firstly, we extracted the M-values of the resulting 781,245, 426,213, and 396,145 CpGs from MESOMICS, TCGA, combined MESOMICS/TCGA, and Iorio cell line cohorts cohorts respectively, which theoretically range from −∞ to +∞ and have a bimodal distribution, being not affected by heteroscedasticity contrary to beta-values (Du et al., 2010). Following the same approach as for expression data, sex-chromosomes CpGs have been excluded (see Methylation section), and from the resulting 781,245, 426,213, 396,145, and 436,125 CpGs available following QC (see DNA methylation Sequencing section), those having less than 0.1 β-value difference across the (**i**) 119, (**iii**) 73, (**iv**) 192, and (**v**) 59 samples have been excluded from the unsupervised analyses. Based on this annotation, the CpGs list representing the methylation data has been divided according to their association with promoters, enhancers or gene body using EPIC 850K array manifest b5 (see ***Regional methylation analysis*** section) resulting in three data sets respectively named MethPro, MethEnh, and MethBod gathering respectively, 37,884, 23,169 and 291,877 CpGs for analysis (**i**); 27,235, 4,953 and 125,228 for analysis (**iii**); 37,951, 5,174, and 132,546 for analysis (**iv**), and 30,387, 4,757, and 111,774 for analysis (**v**). For each analysis and data set, the top 5,000 most variable CpGs were selected.

#### Pre-processing of copy number changes

Copy number changes data was available for both MESOMICS and TCGA cohorts. We assessed the global (Total) and the minor (Minor) allele copy number states at the gene level using respectively the total (*total*) and the minor (*minor*) copy number estimate given by PURPLE (see ***Copy number variant calling*** section) on hg38 genome for the MESOMICS cohort, and SNP array estimates downloaded from the GDC portal for the TCGA MESO cohort. Of note, for the TCGA samples, the copy number state has been aligned on the hg19 genome. Hence, specifically for analysis (**iv**), the transformation of hg19 coordinates into hg38 has been required to integrate copy number data from both MESOMICS and TCGA samples within the same data set. To do so, we used liftOver R package (v.1.14.0) to transform segment coordinates into hg38 genome. Hg19 positions not found by the software because overlapping uncertain regions such as centromeres, have been replaced by the corresponding hg38 centromere coordinates. Then, for the remaining positions not found in the hg38 genome, we first listed, for each segment, the overlapping genes in hg19 and hg38 coordinates and compared the two lists. Then, we saved the same coordinates in case of identical lists and expanded the coordinates to include the overlapping genes that are missing. This expansion has been made only and only if the resulting segment length did not exceed an increase of 5% of the original segment and less than the maximum length difference observed in the transformation process made by liftOver. If these criteria were not filled, the given gene was not included and thus, the coordinates remained unchanged.

For the three analyses, the resulting value assigned to each gene is an average of the copy number estimate of the tumor by taking into account the tumour purity (*purity*) estimated by PURPLE. As a result, *total* = *purity* × *total* + (1-*purity*) × 2 and *minor* = *purity* × *minor* + (1-*purity*) with total and minor the value assigned for each gene in the Total and Minor data set, respectively. In case of segment breaks occurring within a given gene sequence, the mean value of the two segments overlapping is assigned to the gene. In order to avoid redundancy, genes with exactly the same resulting copy number value in all samples (because of their genome location proximity) were grouped as one single feature in the data set. Only the genes or groups of genes altered in at least three samples have been selected. For consistency, each feature of the resulting data sets (10,292 genes or groups of genes—Total and Minor) were centered and scaled to unit variance using the scale R function, and SCNVs occuring on sex-chromosomes were removed. Finally, the top 5,000 most variable genes or groups of genes were selected to be integrated. Note that although available (see **Figure S3A**), because they were computed from exome data instead of genome-wide data as the MESOMICS and TCGA cohorts, CNVs from the Bueno cohort were not included in MOFA.

#### Pre-processing of genomic alterations data

Somatic structural variants data has been used only for the integrative analyses (**i**) and (**iv**), while somatic mutations have been used in all analyses. Each gene, altered by somatic splicing or exonic, damaging mutations or structural variants (see ***Damaging variants and driver detection*** section), has been integrated in a common data set. Of note, for missense mutations, we used REVEL annotation included in ANNOVAR, for predicting the pathogenicity of these variants and used a 0.5 cut-off to restrict to the most likely damaging missense events. We also removed genes altered in less than three samples. For consistency, we selected genes in non sex-chromosomes, protein coding or long non-coding RNA genes and with minimum expression of 0.01 FPKM within the cohort to be sure to include genes expressed in mesothelioma. We integrated the resulting data sets as a boolean variable in the following analyses.

#### Pre-processing of drug response data

We used drug response data (used IC_50_ in units of mean µM) only for the analysis (**v**) on MPM cell lines, combining the drug response of 265 drugs from Iorio. Among them, 3 have also been tested on the de Reyniès cell lines and their responses are reported in Blum et al. (2019).

#### Multi-omic integrative analyses

To provide an integrative low-dimensional summary of the molecular variation across the samples, we performed continuous latent factors identification using software MOFA (R package MOFA2 v1.1.21). Indeed, MOFA is able to integrate different omic data sets by generating independent continuous variables, named latent factors (LF) that transcribe most variation from the joint data sets. In total, we performed four analyses (**i**) MOFA-MESOMICS (*n* = 120, Figures 1 and **S1A**), (**ii**) MOFA-Bueno (*n* = 181, **Figure S1C**), (**iii**) MOFA-TCGA (*n* = 73, **Figure S1C**), (**iv**) MOFA-3-cohorts (*n* = 374, **Figure S1B**), and (**v**) MOFA-Cell lines described above (*n* = 59, **Figure S7A**). Additionally, we ran MOFA on our discovery cohort including the ITH samples (MOFA-ITH, *n* = 134) to evaluate the ITH within MPM samples. Also, note that some samples did not have all the data sets chosen to be integrated available, such as for Bueno and colleagues’ samples missing methylation array data. Fortunately, MOFA was shown to handle missing data, including samples with entire ‘omic techniques missing, by using the correlated signals from several datasets to accurately reconstruct latent factors (Arguelaget et al. 2018).

MOFA was performed independently for each analysis, setting the number of latent factors to 10 (function runMOFA from R package MOFA2 v1.1.21). The summary of all these runs are given in Figures 1, **S1A-G**, and **S7** and coordinates and proportions of variance explained for (**i**)-(**iv**) are given in **Table S2**, while those for MOFA-ITH are given in **Table S4,** and those for the cell lines (model v) are given in **Table S14.** To compare multi-omic with uni-omic unsupervised analyses, we correlated the MOFA coordinates of the samples shared by MOFA and the PCAs with their coordinates in PCA-exp (see ***RNA Sequencing***). Results show that the main 4 MOFA factors all have a counterpart in the PCA. For the MOFA-Cell lines, weights of the features from the drug layer and their correlations with the latent factors are represented in **Figure S7H-I** and **Table S14**.

#### Interpreting MOFA latent factors

We tested the association between each LF and clinical, morphological, and epidemiological variables using linear regression (**Table S2**). As quality control, we also assessed their associations with the technical variables selected to detect potential batch effects in the data using linear regression (**Figure S6D**). The proportion of cells that belong to different immune cell types (see ***Immune contexture deconvolution from expression data*** section) has been quantified and correlated with each dimension using Pearson correlation tests (**Table S2**).

##### Survival prediction

In order to evaluate the ability of the MOFA factors to predict survival, we compared twenty-two survival models: (i) a model based on the three histopathological types (categorical variables, MME, MMB, and MMS); (ii) a model based on the proportion of sarcomatoid content (continuous variable, fitted with a penalized cubic spline); (iii) a model based on the log2 ratio of *CLDN15*/*VIM* (C/V) expression in Bueno and colleagues (2016) (continuous variable, fitted with a penalized cubic spline; see values in **Table S2**); (iv-vi) a model based on the E-score, S-score and the combination of the two, respectively from Blum and colleagues (2019) (continuous variable, fitted with penalized cubic splines without interaction; see values in **Table S2**); (vii) a model based on AI prognostic score (continuous variable, fitted with a penalized cubic spline); (viii-xi) models based on the a-dimensional summary of molecular data using either LF1, LF2, LF3, or LF4 as a continuous variable, respectively (each with a single continuous variable, fitted with a penalized cubic spline); (xii-xxii), models based on the two, three, four-dimensional summary of molecular data using both the combination of two, three, and four of the four LFs as continuous variables, respectively (continuous variables, fitted with penalized cubic spline without interaction). To do so, we assessed their fits using the time-dependent Area Under the ROC Curve (AUC) and its integral (iAUC; Chambless and Diao, 2006; R package survAUC, v.1.0–5), computed using the test set. This time-dependent AUC is used to evaluate the ability of an explanatory variable to predict patients with a survival lower or higher than a given threshold. Its integral summarises the results of time-dependent AUC over the threshold value, providing an interpretation similar to that of classical AUC. In each model, we included sex and age; penalized splines were fitted using the *pspline* function from package survival, with three degrees of freedom. Because of the high proportion of missing asbestos and smoking status information and the absence of significant association between these variables and survival in univariate models (see ***Survival analysis*** section), smoking and asbestos were not included in the model as covariables. Nevertheless, results for survival prediction from these twenty-two models and including both asbestos and smoking status as covariables are also reported in **Table S13**.

To assess the out-of-sample prediction performance, we used 4-fold cross-validation in the MESOMICS cohort (**Figures S6H-J** and Figure 6B). We also assessed the prediction performance on a completely independent cohort by fitting the model on the whole MESOMICS cohort and testing it on the TCGA cohort, using bootstrapping (*n*=2,000 bootstraps) on the test set to assess variation in performance (Figure 6B). Standard errors in the iAUC mean estimate were computed either from the 4 folds or the 2000 bootstraps, respectively for the MESOMICS and TCGA. We also looked at the model fits on the MESOMICS cohort (**Figure S6** E-G and **Table S13**), which confirmed that the MOFA with 4 LFs (xxii) provided the best fit of all models, and also led to the lowest Akaike Information Criteria (AIC=554.324 for the model with the LFs, vs AIC=569.806 for the best AIC of non-MOFA models, that of the S-score + E-score; **Table S13**), which shows that the greater number of parameters in the MOFA survival model is not enough to explain its better performance.

##### Intra tumor heterogeneity analyses

The ploidy, morphology, and CIMP factors represented in Figure 2 were identified in the MOFA-ITH by correlating the coordinates of non-ITH samples in the MOFA-ITH with their coordinates in the MOFA from Figure 1, and choosing the largest match (correlations were all *r*>0.9). To avoid spurious ITH to be detected, and because the ploidy factor overwhelmingly represents variance in genomic data (CNVs; **Table S4**), samples with missing WGD information were not represented in the ploidy factor (NA values in **Table S4**). Similarly, the Pareto front was fitted on the MOFA-ITH using the method described below (**Table S4**). Euclidean distances between each pair of samples were then computed for each factor of the MOFA-ITH separately (Figures 2A **and S2A**). Proportions of the tumor from different components (% Sarcomatoid, % Acinar, % immune infiltration) presented in Figure 2B matched that reported by the pathologist, and include the constraint that % Sarcomatoid + % Epithelioid + % infiltration=100%, and % Acinar <= % Epithelioid (see data in **Table S4**).

### Evolutionary tumor trade-off analyses

#### Pareto theory fit

The Pareto front model has been fitted to different sets of samples using the ParetoTI R package (https://github.com/vitkl/ParetoTI, release v0.1.13), following the above mentioned analyses (**i**), (**ii**), (**iii**) and (**iv**), and additionally, on two different kinds of molecular maps: using MOFA, restricting to latent factors LF1, LF2, LF3, and LF4 and using expression PCA as technical validation (see ***RNA Sequencing***). In brief, the algorithm tries to find polyhedra by testing successively 1 to *n* axes adding them one after another in a decreasing order of transcriptomic variance explained. For this technical reason, the MOFA LFs have been ordered as follows by decreasing transcriptomic variance explained: Morphology factor (LF2), Adaptive-response factor (LF3), CIMP factor (LF4), and Ploidy factor (LF1). For each number *n* of axes used, ParetoTI identifies the position of the *n*+1 = *k* vertices (archetypes) in the molecular map defined. Each polyhedron fit is assessed by the ratio of the volume of the best-fitting polyhedron to the volume of the convex hull of the data (t-ratio). The more the data follows the Pareto optimality theory, the more the *t*-ratio metric, higher than 1, approaches 1. Finally, the algorithm re-calculates the *t*-ratio on 1000 shuffles keeping the distribution of loading on each axis but not the associations between them and computes a one-sided *p*-value to estimate the statistical significance of the fit.

Here, we chose to represent the most significant fit with the smallest number *k* because of the limited number of samples. Using MOFA axes, we found *k* = 3 archetypes in the LF2-LF3 space and reproduced, for each analysis (**i**), (**ii**), (**iii**), and (**iv**), the fit using the corresponding expression PCA in the PC1-PC2 space (see **Figure S1G** for the fit for model ii, and iii). In order to evaluate the reproduction of the three archetypes discovered in (**i**) (MESOMICS cohort) into (**ii**) (Bueno’s cohorts) and (**iii**) (TCGA cohort), we used (**iv**) (3-cohorts) and correlated the pairwise distance between archetypes and samples within each molecular map (**Table S3**). Overall, we found a strong concordance between the three analyses (minimum absolute Pearson’s *r* = 0.84).

#### Interpretation of MPM polyhedron

To further characterise the phenotype of each archetype we used the proportion of each archetype for each sample estimated by ParetoTI. These proportions have been used as continuous variables to further test the association between each archetype and clinical, epidemiological, morphological variables, as well as molecular data (**Table S3**).

More specifically, we inferred each archetype phenotype by performing integrative gene set enrichment analysis (IGSEA) on the expression data. To do so, we used the ActivePathways r package (https://github.com/reimandlab/ActivePathways, release v1.02) which is a tool able to integrate different sources of molecular variations to assess the enrichment of GO terms, by combining *p*-values from different association tests between sources and gene level data. Here, we integrated these proportions as different axes of molecular variation. We restricted the GO terms to a minimum size of 20 genes to a maximum size of 1000 genes as the default parameters of ActivePathways. To infer the pathways specifically altered in each archetype, we integrated the Pearson’s *p*-value correlation of each gene from the expression matrix of 59,607 genes with the proportion from each archetype and we selected the pathways for which the enrichment source only correspond to the tested archetype. We performed two kinds of analyses: one restricted to the genes positively correlated with the proportion to get the upregulated pathways and a second one restricted to the negatively correlated genes to identify the down-regulated pathways. In all the analyses, the proportion of enriched genes within the enriched pathways ranged from 0.04 to 0.75 (**Table S3**). Used as a quality control of the enrichment results, we assessed the fold change between the 10% closest samples vs the 10% furthest samples from each archetype of the enriched genes belonging to each enriched pathway. More specifically, in order to assign universal cancer task to each archetype, we referred to Hausser et al. (2019) and examined the GO term descriptions to gather pathways in super-pathways as reported in **Table S3**.

Similarly, we tested the association between each archetype and genomic event using linear regression and more specifically, in order to infer genomic event effect-size on the Pareto front, we calculated the vector linking the centroids of the altered and wild-type groups (*centroids* function from sda R package v.1.3.7). To infer to what extent alterations drives the tumour cells toward specialisation, we followed the method from Hausser et al. (2019) and calculated the alignment of vectors with the front (angle between the Pareto front and the vector built from the altered and wild-type groups within the 4-factors space), after having normalised each LF (centering and division by standard deviation). Finally, we evaluated the driving role of genomic events associated to at least one archetype (**Table S11**) using these two variables (vector size and angle to Pareto front) by permutation tests (with 1,000 permutations) in which we randomised, one genomic event at a time, the altered and wild-type groups and compared this distribution from shuffled values with the observed values (Figure 5*C*).

### Clonal reconstruction

#### Small variants subclonal reconstruction

To obtain accurate estimates of variant allelic fractions (VAFs), we restricted the model fitting to somatic alterations with high-confidence VAF estimations (read depth greater than or equal to 60X and VAF greater than or equal to 0.05), and focused on samples with matched normal tissue or blood (*n*=43). We only selected variants in high-confidence CN calls: regions with confident calls, excluding centromeric regions (based on UCSC annotation) that have notoriously more difficult CNV calling due to larger variance in reads mapping.

We inferred whether small somatic variants were clonal or subclonal using R package *MOBSTER* v1.00. MOBSTER uses a mixture model to identify different clones in the VAF distribution. Importantly, the model uses evolutionary theory predictions to perform more accurate subclonal reconstructions, and can test whether subclones are under natural selection or neutral evolution by testing the presence of a “neutral tail” component, a Pareto type I distribution that is expected to be present in exponentially growing tumors evolving under neutral evolution (Williams et al., 2016). For each sample, we first fit a mixture model to the VAF distribution from variants in regions with the most frequent CN (major and minor CN of 1 for most samples, major and minor CN of 2 and 1 or 3 and 1 for WGD samples, and 1 and 0 or 2 and 0 for GNH samples). For each sample, we compared the fit of models with or without neutral tail and with 1 to 3 clusters, with 10 repetitions per model with different initializations, resulting in 6×10 models per sample; we chose the best model using the ICL statistic. We assessed the robustness of the fit using parametric bootstrapping; only models that were correctly inferred in more than 80% of the simulations were used. Of Note, the clonal cluster also provides an estimate of the sample purity based on the VAF distribution. Multiple clusters denoting the presence of a subclone were identified in 13 samples. All 13 samples presented a single subclone, and all were in the low-adaptive response factor range (close to archetype 3), as expected from the high purity required to detect subclonal alterations (see **Table S1**); 3 samples presented a neutral tail, while the 10 others presented a selected subclone (**Figure S5I**).

We finally assigned mutations that were not included in the model fit (small variants in regions with another CN, subclonal CNVs) to clones and subclones using their VAF and the fitted model. For each of the 13 samples where a subclone was identified, we recovered the cutoff cancer cell fraction (CCF) separating clonal and subclonal alterations according to the selected MOBSTER model. We then converted this threshold CCF to a threshold VAF by taking into account the CN state of each alteration using the formula VAF_thres_ = CCF_thres_*ɸ*/[CN_normal_ ✕ (1-*ɸ*) + CN_total_ ✕ *ɸ*], where CN_normal_ is 2 for autosomal regions and 1 for sex chromosomes, CN_total_ is the total CN of the tumor, and *ɸ* is the MOBSTER-estimated purity. Variants were then assigned to the clonal and subclonal categories depending on which side of the threshold they fell. Note that this approach is similar to that used in the DPClust software (Nik-Zainal et al., 2012), but using the recent evolutionary-theory aware probability distributions from MOBSTER instead of Dirichlet distributions. The proportion of clonal and subclonal alterations in the 13 samples where this analysis was possible are reported in **Table S1** and **Figure S5J**.

Note that the small number of samples with such clusters of subclonal alterations detected allowed to further check visually the consistency between the fits of different CN regions (e.g., CN neutral LOH regions should have two clonal modes-one corresponding to variants in 1 or 2 copies-while diploid regions should have only clonal mode). Results for alterations in the driver list from Figure 4 are presented in **Figures S5G** and **I**.

#### CNV clonality reconstruction

Clonality of CNVs was assessed using the estimated fractional copy number from PURPLE. Indeed, the PURPLE algorithm uses a penalised estimation of CN so that clonal CN segments are expected to have CN values close to an integer while subclonal segments have non-integer CN values; we thus classified as subclonal segments with a CN deviating from an integer value (fractional part between 0.2 and 0.8). Because of the difficulty of inferring the clonality of CNVs, we also assessed the clonality of CNVs using software Facets (Shen and Seshan, 2016; https://github.com/IARCbioinfo/facets-nf; release 2.0); only CNVs consistently called as clonal or subclonal by PURPLE and Facets are reported in **Figure S5H** (see **Table S7** for all CNV clonalities). Although CNVs are generally called more accurately than small variants in T-only samples, for consistency with the rest of the clonality and evolutionary analyses, we restricted the analyses to tumors with a matched normal (*n*=43).

### Inferring the timing of alterations

Due to the low tumor mutational burden in MPM, we restricted the analysis to samples with large scale events (WGD or more than 10% of the genome with LOH), and to samples with matched normal tissue or blood.

#### Molecular time dating

Similarly to the approach from Gerstung and colleagues (2020) implemented in package MutationTimeR, copy number gains and copy neutral losses of heterozygosity (LOH) were dated by comparing the number of alterations that were present in a single copy (that appeared after the event), *N_r_*, to the ones that were present in multiple copies (that appeared before the event) *N_l_* (**Figure S5F**).

We computed Bayesian credibility intervals (BCI) for the timing of each gain and compared results with parametric bootstrapping confidence intervals (CI). BCI were obtained by assuming that *N_r_* followed a Poisson distribution of parameters *λ_r_* and *λ_l_*, and uniform prior distributions over the interval [0,10^4^] with the constraints that *λ_r_* < *λ_l_ ,* because the mutation rate *λ_l_* includes both mutations that occurred before and after the copy number gain, while *λ_r_* is limited to mutations that occurred before the gain; posterior distributions were numerically computed using a discrete grid approximation of size 1001, and used to compute the posterior distribution of timings *t_e_*. Bootstrapping CI proceeded as in Gerstung and colleagues (2020), first drawing 1000 *N_ri_* values from a Poisson distribution of parameter *N_r_*, and finally inferring *t_e_* from the simulations. We show in **Figure S5E** that both approaches provide very similar results (correlation between the center of the CI and BCI across dated events is *r*=0.99, *p*=4.8×10^-14^), but the Bayesian estimates have the advantage of ensuring that 0<*t_e_<*1, because of the prior imposing that *λ_r_* < *λ_l_* , so they are the ones reported in the main text.

Synchronicity of duplications in the WGD sample was assessed by checking the overlap between CI for gains in the different segments considered.

#### Chronological time dating

We used the method validated by Gerstung and colleagues to date amplification events (Gerstung et al., 2020) first estimated the temporal accumulation of CpG to TpG mutations ([C>T]pG), mostly due to spontaneous de-aminations that would accumulate at an approximately constant rate through time. In order to check whether the small number of alterations present in some segments led to biases in our estimates, we performed the same analysis but using all mutations instead of just [C>T]pG mutations. Results showed no significant systematic bias (linear regression coefficient estimated at 0.85, with 95%CI [0.69,1.01]; **Figure S5D**), and CI of [C>T]pG and CI of all mutations overlapped except for MESO_008, a non-chemonaive tumor that showed an excess of chemotherapy associated mutational signatures that likely influenced the proportion of signatures associated with ageing relative to other signatures. Overall, our results show that using all mutations increases the precision of estimates but does not bias the results as long as we exclude non-chemonaive samples, probably because MPM do not have SBS signatures of exogenous sources but rather only a slow temporal accumulation of mutations (see ***mutational signatures section***), so results in the main text correspond to results for all mutations. Finally, we checked whether mutation accumulation showed a sign of temporal acceleration by comparing the number of small variants corrected for the effective genome size (defined as in Gerstung et al. 2020 as 1/mean(*m_i_*/*C_i_*), with *m_i_* the number of copies of alteration *i* and *C_i_* the total CN at this position) with the age at diagnosis (**Figure S5G**); the analysis showed that small variants fit a linear accumulation model, thus we used a rate of x1 for chronological dating.

## Supplemental Information

### List of Supplementary Tables

Table S1. Sample overview.

Table S2. MOFA MESOMICS and its associations, replication in the TCGA and Bueno cohorts.

Table S3. Pareto front, associations, and IGSEA of cancer task archetypes.

Table S4. ITH analyses.

Table S5. Small variants.

Table S6. Structural variants (SVs) and fusion transcripts.

Table S7. Copy number variants (CNVs).

Table S8. Amplification and deletion peaks and broad events (GISTIC2).

Table S9. Complex genomic events.

Table S10. Differential expression analysis between WGD- and WGD+ samples.

Table S11. Association between MOFA factors or Archetype proportions and genomic events.

Table S12. Tumor suppressor genes (TSG) and CIMP-index.

Table S13. Survival analyses.

Table S14. MOFA Cell lines.

### List of Supplementary Figures

Figure S1. MOFA of the MESOMICS, TCGA, and Bueno cohorts.

Figure S2. Patterns of intra-tumor heterogeneity in 13 samples.

Figure S3. Detailed genomic profiles.

Figure S4. MPM driver detection.

Figure S5. Detailed impact of genomic alterations on molecular profiles of MPM.

Figure S6. Association between survival and MOFA in the MESOMICS cohort.

Figure S7. MOFA of cell lines.

## References

Alcala, N., Mangiante, L., Le-Stang, N., Gustafson, C.E., Boyault, S., Damiola, F., Alcala, K., Brevet, M., Thivolet-Bejui, F., Blanc-Fournier, C., et al. (2019a). Redefining malignant pleural mesothelioma types as a continuum uncovers immune-vascular interactions. EBioMedicine 48, 191–202.

Alcala, N., Leblay, N., Gabriel, A.A.G., Mangiante, L., Hervas, D., Giffon, T., Sertier, A.S., Ferrari, A., Derks, J., Ghantous, A., et al. (2019b). Integrative and comparative genomic analyses identify clinically relevant pulmonary carcinoid groups and unveil the supra-carcinoids. Nat. Commun. 10, 3407.

Alexandrov, L.B., Kim, J., Haradhvala, N.J., Huang, M.N., Tian Ng, A.W., Wu, Y., Boot, A., Covington, K.R., Gordenin, D.A., Bergstrom, E.N., et al. (2020). The repertoire of mutational signatures in human cancer. Nature 578, 94–101.

Argelaguet, R., Velten, B., Arnol, D., Dietrich, S., Zenz, T., Marioni, J.C., Buettner, F., Huber, W., and Stegle, O. (2018). Multi-Omics Factor Analysis-a framework for unsupervised integration of multi-omics data sets. Mol. Syst. Biol. 14, e8124.

Argelaguet, R., Arnol, D., Bredikhin, D., Deloro, Y., Velten, B., Marioni, J.C., and Stegle, O. (2020). MOFA+: a statistical framework for comprehensive integration of multi-modal single-cell data. Genome Biol. 21, 111.

Baas, P., Fennell, D., Kerr, K.M., Van Schil, P.E., Haas, R.L., and Peters, S. (2015). Malignant pleural mesothelioma: ESMO Clinical Practice Guidelines for diagnosis, treatment and follow-up. Annals of Oncology 26, v31–v39.

Baas, P., Scherpereel, A., Nowak, A.K., Fujimoto, N., Peters, S., Tsao, A.S., Mansfield, A.S., Popat, S., Jahan, T., Antonia, S., et al. (2021a). First-line nivolumab plus ipilimumab in unresectable malignant pleural mesothelioma (CheckMate 743): a multicentre, randomised, open-label, phase 3 trial. Lancet 397, 375–386.

Baas, P., Scherpereel, A., Nowak, A.K., Oukessou, A., and Zalcman, G. (2021b). Heterogeneity of treatment effects in malignant pleural mesothelioma - Authors’ reply. Lancet 398, 302.

Baylin, S.B., and Jones, P.A. (2016). Epigenetic Determinants of Cancer. Cold Spring Harb. Perspect. Biol. 8.

Benjamin, D., Sato, T., Cibulskis, K., Getz, G., Stewart, C., and Lichtenstein, L. Calling Somatic SNVs and Indels with Mutect2.

Bergstrom, E.N., Luebeck, J., Petljak, M., Bafna, V., Mischel, P.S., Harris, R.S., and Alexandrov, L.B. (2021). Comprehensive analysis of clustered mutations in cancer reveals recurrent APOBEC3 mutagenesis of ecDNA.

Bielski, C.M., Zehir, A., Penson, A.V., Donoghue, M.T.A., Chatila, W., Armenia, J., Chang, M.T., Schram, A.M., Jonsson, P., Bandlamudi, C., et al. (2018). Genome doubling shapes the evolution and prognosis of advanced cancers. Nat. Genet. 50, 1189–1195.

Blum, Y., Meiller, C., Quetel, L., Elarouci, N., Ayadi, M., Tashtanbaeva, D., Armenoult, L., Montagne, F., Tranchant, R., Renier, A., et al. (2019). Dissecting heterogeneity in malignant pleural mesothelioma through histo-molecular gradients for clinical applications. Nat. Commun. 10, 1333.

Bueno, R., Stawiski, E.W., Goldstein, L.D., Durinck, S., De Rienzo, A., Modrusan, Z., Gnad, F., Nguyen, T.T., Jaiswal, B.S., Chirieac, L.R., et al. (2016). Comprehensive genomic analysis of malignant pleural mesothelioma identifies recurrent mutations, gene fusions and splicing alterations. Nat. Genet. 48, 407–416.

Caravagna, G., Sanguinetti, G., Graham, T.A., and Sottoriva, A. (2020). The MOBSTER R package for tumour subclonal deconvolution from bulk DNA whole-genome sequencing data. BMC Bioinformatics 21, 531.

Carbone, M., Adusumilli, P.S., Alexander, H.R., Jr, Baas, P., Bardelli, F., Bononi, A., Bueno, R., Felley-Bosco, E., Galateau-Salle, F., Jablons, D., et al. (2019). Mesothelioma: Scientific clues for prevention, diagnosis, and therapy. CA Cancer J. Clin. 69, 402–429.

Chambless, L.E., and Diao, G. (2006). Estimation of time-dependent area under the ROC curve for long-term risk prediction. Stat. Med. 25, 3474–3486.

Chapel, D.B., Schulte, J.J., Berg, K., Churg, A., Dacic, S., Fitzpatrick, C., Galateau-Salle, F., Hiroshima, K., Krausz, T., Le Stang, N., et al. (2020). MTAP immunohistochemistry is an accurate and reproducible surrogate for CDKN2A fluorescence in situ hybridization in diagnosis of malignant pleural mesothelioma. Mod. Pathol. 33, 245–254.

Chen, X., Schulz-Trieglaff, O., Shaw, R., Barnes, B., Schlesinger, F., Källberg, M., Cox, A.J., Kruglyak, S., and Saunders, C.T. (2016). Manta: rapid detection of structural variants and indels for germline and cancer sequencing applications. Bioinformatics 32, 1220–1222.

Chen, Y.-A., Lemire, M., Choufani, S., Butcher, D.T., Grafodatskaya, D., Zanke, B.W., Gallinger, S., Hudson, T.J., and Weksberg, R. (2013). Discovery of cross-reactive probes and polymorphic CpGs in the Illumina Infinium HumanMethylation450 microarray. Epigenetics 8, 203–209.

Choi, J., Southworth, L.K., Sarin, K.Y., Venteicher, A.S., Ma, W., Chang, W., Cheung, P., Jun, S., Artandi, M.K., Shah, N., et al. (2008). TERT promotes epithelial proliferation through transcriptional control of a Myc- and Wnt-related developmental program. PLoS Genet. 4, e10.

Cortés-Ciriano, I., Lee, J.J.-K., Xi, R., Jain, D., Jung, Y.L., Yang, L., Gordenin, D., Klimczak, L.J., Zhang, C.-Z., Pellman, D.S., et al. (2020). Comprehensive analysis of chromothripsis in 2,658 human cancers using whole-genome sequencing. Nat. Genet. 52, 331–341.

Courtiol, P., Maussion, C., Moarii, M., Pronier, E., Pilcer, S., Sefta, M., Manceron, P., Toldo, S., Zaslavskiy, M., Le Stang, N., et al. (2019). Deep learning-based classification of mesothelioma improves prediction of patient outcome. Nat. Med. 25, 1519–1525.

Damotte, D., Warren, S., Arrondeau, J., Boudou-Rouquette, P., Mansuet-Lupo, A., Biton, J., Ouakrim, H., Alifano, M., Gervais, C., Bellesoeur, A., et al. (2019). The tumor inflammation signature (TIS) is associated with anti-PD-1 treatment benefit in the CERTIM pan-cancer cohort. J. Transl. Med. 17, 357.

De Rienzo, A., Archer, M.A., Yeap, B.Y., Dao, N., Sciaranghella, D., Sideris, A.C., Zheng, Y., Holman, A.G., Wang, Y.E., Dal Cin, P.S., et al. (2016). Gender-Specific Molecular and Clinical Features Underlie Malignant Pleural Mesothelioma. Cancer Res. 76, 319–328.

Deshpande, V., Luebeck, J., Nguyen, N.-P.D., Bakhtiari, M., Turner, K.M., Schwab, R., Carter, H., Mischel, P.S., and Bafna, V. (2019). Exploring the landscape of focal amplifications in cancer using AmpliconArchitect. Nat. Commun. 10, 392.

Di Maio, M., and Tagliamento, M. (2021). Heterogeneity of treatment effects in malignant pleural mesothelioma. Lancet 398, 301–302.

Di Tommaso, P., Chatzou, M., Floden, E.W., Barja, P.P., Palumbo, E., and Notredame, C. (2017). Nextflow enables reproducible computational workflows. Nat. Biotechnol. 35, 316–319.

Du, P., Zhang, X., Huang, C.-C., Jafari, N., Kibbe, W.A., Hou, L., and Lin, S.M. (2010). Comparison of Beta-value and M-value methods for quantifying methylation levels by microarray analysis. BMC Bioinformatics 11, 587.

Dulloo, S., Bzura, A., and Fennell, D.A. (2021). Precision Therapy for Mesothelioma: Feasibility and New Opportunities. Cancers 13.

Federico, A., and Monti, S. (2020). hypeR: an R package for geneset enrichment workflows. Bioinformatics 36, 1307–1308.

Fernandez-Cuesta, L., Mangiante, L., Alcala, N., and Foll, M. (2021). Challenges in lung and thoracic pathology: molecular advances in the classification of pleural mesotheliomas. Virchows Arch. 478, 73–80.

Fu, Y., Jung, A.W., Torne, R.V., Gonzalez, S., Vöhringer, H., Shmatko, A., Yates, L.R., Jimenez-Linan, M., Moore, L., and Gerstung, M. (2020). Pan-cancer computational histopathology reveals mutations, tumor composition and prognosis. Nature Cancer 1, 800–810.

Gabriel, A.A.G., Atkins, J.R., Penha, R.C.C., Smith-Byrne, K., Gaborieau, V., Voegele, C., Abedi-Ardekani, B., Milojevic, M., Olaso, R., Meyer, V., et al. (2021). Genetic analysis of lung cancer reveals novel susceptibility loci and germline impact on somatic mutation burden. medRxiv 2021.04.26.21254132.

Galateau-Salle, F., Churg, A., Roggli, V., and Travis, W.D. (2016). The 2015 World Health Organization Classification of Tumors of the Pleura: Advances since the 2004 Classification. Journal of Thoracic Oncology 11, 142–154.

Gerstung, M., Jolly, C., Leshchiner, I., Dentro, S.C., Gonzalez, S., Rosebrock, D., Mitchell, T.J., Rubanova, Y., Anur, P., Yu, K., et al. (2020). The evolutionary history of 2,658 cancers. Nature 578, 122–128.

Ghanim, B., Klikovits, T., Hoda, M.A., Lang, G., Szirtes, I., Setinek, U., Rozsas, A., Renyi-Vamos, F., Laszlo, V., Grusch, M., et al. (2015). Ki67 index is an independent prognostic factor in epithelioid but not in non-epithelioid malignant pleural mesothelioma: a multicenter study. Br. J. Cancer 112, 783–792.

Gray, S.G. (2021). Emerging avenues in immunotherapy for the management of malignant pleural mesothelioma. BMC Pulm. Med. 21, 148.

Griffin, B.A., Anderson, G.L., Shih, R.A., and Whitsel, E.A. (2012). Use of alternative time scales in Cox proportional hazard models: implications for time-varying environmental exposures. Stat. Med. 31, 3320–3327.

Halaburkova, A., Cahais, V., Novoloaca, A., Araujo, M.G. da S., Khoueiry, R., Ghantous, A., and Herceg, Z. (2020). Pan-cancer multi-omics analysis and orthogonal experimental assessment of epigenetic driver genes. Genome Res. 30, 1517–1532.

Hatzikirou, H., Basanta, D., Simon, M., Schaller, K., and Deutsch, A. (2012). “Go or grow”: the key to the emergence of invasion in tumour progression? Math. Med. Biol. 29, 49–65.

Hausser, J., and Alon, U. (2020). Tumour heterogeneity and the evolutionary trade-offs of cancer. Nat. Rev. Cancer 20, 247–257.

Hausser, J., Szekely, P., Bar, N., Zimmer, A., Sheftel, H., Caldas, C., and Alon, U. (2019). Tumor diversity and the trade-off between universal cancer tasks. Nat. Commun. 10, 5423.

Helleday, T. (2011). The underlying mechanism for the PARP and BRCA synthetic lethality: clearing up the misunderstandings. Mol. Oncol. 5, 387–393.

Hmeljak, J., Sanchez-Vega, F., Hoadley, K.A., Shih, J., Stewart, C., Heiman, D., Tarpey, P., Danilova, L., Drill, E., Gibb, E.A., et al. (2018). Integrative Molecular Characterization of Malignant Pleural Mesothelioma. Cancer Discov. 8, 1548–1565.

Hylebos, M., Op de Beeck, K., van den Ende, J., Pauwels, P., Lammens, M., van Meerbeeck, J.P., and Van Camp, G. (2018). Molecular analysis of an asbestos-exposed Belgian family with a high prevalence of mesothelioma. Fam. Cancer 17, 569–576.

IARC/WHO (2015). WHO Classification of Tumours of the Lung, Pleura, Thymus and Heart (International Agency for Research on Cancer).

soit Ilg, A.G., Ducamp, S., Grange, D., Audignon, S., Gramond, C., Le Stang, N., Frenay, C., Galateau-Sallé, F., Pairon, J.-C., Astoul, P., et al. (2020). Programme national de surveillance du mésothéliome pleural (PNSM): vingt années de surveillance des cas, de leurs expositions et de leur reconnaissance médico-sociale (1998-2017). Archives Des Maladies Professionnelles et de l’Environnement 81, 672.

Iorio, F., Knijnenburg, T.A., Vis, D.J., Bignell, G.R., Menden, M.P., Schubert, M., Aben, N., Gonçalves, E., Barthorpe, S., Lightfoot, H., et al. (2016). A Landscape of Pharmacogenomic Interactions in Cancer. Cell 166, 740–754.

Jeffares, D.C., Jolly, C., Hoti, M., Speed, D., Shaw, L., Rallis, C., Balloux, F., Dessimoz, C., Bähler, J., and Sedlazeck, F.J. (2017). Transient structural variations have strong effects on quantitative traits and reproductive isolation in fission yeast. Nat. Commun. 8, 14061.

Kato, S., Tomson, B.N., Buys, T.P.H., Elkin, S.K., Carter, J.L., and Kurzrock, R. (2016). Genomic Landscape of Malignant Mesotheliomas. Mol. Cancer Ther. 15, 2498–2507.

Kim, S., Scheffler, K., Halpern, A.L., Bekritsky, M.A., Noh, E., Källberg, M., Chen, X., Kim, Y., Beyter, D., Krusche, P., et al. (2018). Strelka2: fast and accurate calling of germline and somatic variants. Nat. Methods 15, 591–594.

Ladan, M.M., van Gent, D.C., and Jager, A. (2021). Homologous Recombination Deficiency Testing for BRCA-Like Tumors: The Road to Clinical Validation. Cancers 13.

LaFave, L.M., Béguelin, W., Koche, R., Teater, M., Spitzer, B., Chramiec, A., Papalexi, E., Keller, M.D., Hricik, T., Konstantinoff, K., et al. (2015). Loss of BAP1 function leads to EZH2-dependent transformation. Nat. Med. 21, 1344–1349.

Ledermann, J., Harter, P., Gourley, C., Friedlander, M., Vergote, I., Rustin, G., Scott, C.L., Meier, W., Shapira-Frommer, R., Safra, T., et al. (2014). Olaparib maintenance therapy in patients with platinum-sensitive relapsed serous ovarian cancer: a preplanned retrospective analysis of outcomes by BRCA status in a randomised phase 2 trial. Lancet Oncol. 15, 852–861.

Levatic, J., Salvadores, M., Fuster-Tormo, F., and Supek, F. (2021). Mutational signatures are markers of drug sensitivity of cancer cells. bioRxiv.

Li, H., and Durbin, R. (2009). Fast and accurate short read alignment with Burrows-Wheeler transform. Bioinformatics 25, 1754–1760.

Liu, Z., Li, Q., Li, K., Chen, L., Li, W., Hou, M., Liu, T., Yang, J., Lindvall, C., Björkholm, M., et al. (2013). Telomerase reverse transcriptase promotes epithelial-mesenchymal transition and stem cell-like traits in cancer cells. Oncogene 32, 4203–4213.

Lopez, G., Egolf, L.E., Giorgi, F.M., Diskin, S.J., and Margolin, A.A. (2021). svpluscnv: analysis and visualization of complex structural variation data. Bioinformatics 37, 1912–1914.

Lutsik, P., Slawski, M., Gasparoni, G., Vedeneev, N., Hein, M., and Walter, J. (2017). MeDeCom: discovery and quantification of latent components of heterogeneous methylomes. Genome Biol. 18, 55.

Mak, M.P., Tong, P., Diao, L., Cardnell, R.J., Gibbons, D.L., William, W.N., Skoulidis, F., Parra, E.R., Rodriguez-Canales, J., Wistuba, I.I., et al. (2016). A Patient-Derived, Pan-Cancer EMT Signature Identifies Global Molecular Alterations and Immune Target Enrichment Following Epithelial-to-Mesenchymal Transition. Clin. Cancer Res. 22, 609–620.

Mansfield, A.S., Peikert, T., Smadbeck, J.B., Udell, J.B.M., Garcia-Rivera, E., Elsbernd, L., Erskine, C.L., Van Keulen, V.P., Kosari, F., Murphy, S.J., et al. (2019). Neoantigenic Potential of Complex Chromosomal Rearrangements in Mesothelioma. J. Thorac. Oncol. 14, 276–287.

Margueron, R., and Reinberg, D. (2011). The Polycomb complex PRC2 and its mark in life. Nature 469, 343–349.

Martínez-Jiménez, F., Muiños, F., Sentís, I., Deu-Pons, J., Reyes-Salazar, I., Arnedo-Pac, C., Mularoni, L., Pich, O., Bonet, J., Kranas, H., et al. (2020). A compendium of mutational cancer driver genes. Nat. Rev. Cancer 20, 555–572.

Mayakonda, A., Lin, D.-C., Assenov, Y., Plass, C., and Koeffler, H.P. (2018). Maftools: efficient and comprehensive analysis of somatic variants in cancer. Genome Res. 28, 1747–1756.

McLoughlin, K.C., Kaufman, A.S., and Schrump, D.S. (2017). Targeting the epigenome in malignant pleural mesothelioma. Transl Lung Cancer Res 6, 350–365.

Mermel, C.H., Schumacher, S.E., Hill, B., Meyerson, M.L., Beroukhim, R., and Getz, G. (2011). GISTIC2.0 facilitates sensitive and confident localization of the targets of focal somatic copy-number alteration in human cancers. Genome Biol. 12, R41.

Mose, L.E., Perou, C.M., and Parker, J.S. (2019). Improved indel detection in DNA and RNA via realignment with ABRA2. Bioinformatics 35, 2966–2973.

Nguyen, L., W M Martens, J., Van Hoeck, A., and Cuppen, E. (2020). Pan-cancer landscape of homologous recombination deficiency. Nat. Commun. 11, 5584.

Nicholson, A.G., Sauter, J.L., Nowak, A.K., Kindler, H.L., Gill, R.R., Remy-Jardin, M., Armato, S.G., 3rd, Fernandez-Cuesta, L., Bueno, R., Alcala, N., et al. (2020). EURACAN/IASLC Proposals for Updating the Histologic Classification of Pleural Mesothelioma: Towards a More Multidisciplinary Approach. J. Thorac. Oncol. 15, 29–49.

Nik-Zainal, S., Van Loo, P., Wedge, D.C., Alexandrov, L.B., Greenman, C.D., Lau, K.W., Raine, K., Jones, D., Marshall, J., Ramakrishna, M., et al. (2012). The life history of 21 breast cancers. Cell 149, 994–1007.

Pastorino, S., Yoshikawa, Y., Pass, H.I., Emi, M., Nasu, M., Pagano, I., Takinishi, Y., Yamamoto, R., Minaai, M., Hashimoto-Tamaoki, T., et al. (2018). A Subset of Mesotheliomas With Improved Survival Occurring in Carriers of BAP1 and Other Germline Mutations. J. Clin. Oncol. JCO2018790352.

PCAWG Consortium (2020). Pan-cancer analysis of whole genomes. Nature 578, 82–93.

Pertea, M., Pertea, G.M., Antonescu, C.M., Chang, T.-C., Mendell, J.T., and Salzberg, S.L. (2015). StringTie enables improved reconstruction of a transcriptome from RNA-seq reads. Nat. Biotechnol. 33, 290–295.

Pirker, C., Bilecz, A., Grusch, M., Mohr, T., Heidenreich, B., Laszlo, V., Stockhammer, P., Lötsch-Gojo, D., Gojo, J., Gabler, L., et al. (2020). Telomerase Reverse Transcriptase Promoter Mutations Identify a Genomically Defined and Highly Aggressive Human Pleural Mesothelioma Subgroup. Clin. Cancer Res. 26, 3819–3830.

Priestley, P., Baber, J., Lolkema, M.P., Steeghs, N., de Bruijn, E., Shale, C., Duyvesteyn, K., Haidari, S., van Hoeck, A., Onstenk, W., et al. (2019). Pan-cancer whole-genome analyses of metastatic solid tumours. Nature 575, 210–216.

Quetel, L., Meiller, C., Assié, J.-B., Blum, Y., Imbeaud, S., Montagne, F., Tranchant, R., de Wolf, J., Caruso, S., Copin, M.-C., et al. (2020). Genetic alterations of malignant pleural mesothelioma: association with tumor heterogeneity and overall survival. Mol. Oncol. 14, 1207–1223.

Quinton, R.J., DiDomizio, A., Vittoria, M.A., Kotýnková, K., Ticas, C.J., Patel, S., Koga, Y., Vakhshoorzadeh, J., Hermance, N., Kuroda, T.S., et al. (2021). Whole-genome doubling confers unique genetic vulnerabilities on tumour cells. Nature 590, 492–497.

Rausch, T., Zichner, T., Schlattl, A., Stütz, A.M., Benes, V., and Korbel, J.O. (2012). DELLY: structural variant discovery by integrated paired-end and split-read analysis. Bioinformatics 28, i333–i339.

de Reyniès, A., Jaurand, M.-C., Renier, A., Couchy, G., Hysi, I., Elarouci, N., Galateau-Sallé, F., Copin, M.-C., Hofman, P., Cazes, A., et al. (2014). Molecular classification of malignant pleural mesothelioma: identification of a poor prognosis subgroup linked to the epithelial-to-mesenchymal transition. Clin. Cancer Res. 20, 1323–1334.

Schaeffner, E.S., Miller, D.P., Wain, J.C., and Christiani, D.C. (2001). Use of an asbestos exposure score and the presence of pleural and parenchymal abnormalities in a lung cancer case series. Int. J. Occup. Environ. Health 7, 14–18.

Shen, R., and Seshan, V.E. (2016). FACETS: allele-specific copy number and clonal heterogeneity analysis tool for high-throughput DNA sequencing. Nucleic Acids Res. 44, e131.

Shipony, Z., Mukamel, Z., Cohen, N.M., Landan, G., Chomsky, E., Zeliger, S.R., Fried, Y.C., Ainbinder, E., Friedman, N., and Tanay, A. (2014). Dynamic and static maintenance of epigenetic memory in pluripotent and somatic cells. Nature 513, 115–119.

Shukuya, T., Serizawa, M., Watanabe, M., Akamatsu, H., Abe, M., Imai, H., Tokito, T., Ono, A., Taira, T., Kenmotsu, H., et al. (2014). Identification of actionable mutations in malignant pleural mesothelioma. Lung Cancer 86, 35–40.

Sondka, Z., Bamford, S., Cole, C.G., Ward, S.A., Dunham, I., and Forbes, S.A. (2018). The COSMIC Cancer Gene Census: describing genetic dysfunction across all human cancers. Nat. Rev. Cancer 18, 696–705.

Sreejit, G., Ahmed, A., Parveen, N., Jha, V., Valluri, V.L., Ghosh, S., and Mukhopadhyay, S. (2014). The ESAT-6 protein of Mycobacterium tuberculosis interacts with beta-2-microglobulin (β2M) affecting antigen presentation function of macrophage. PLoS Pathog. 10, e1004446.

Steele, C.D., Abbasi, A., Ashiqul Islam, S.M., Khandekar, A., Haase, K., Hames, S., Tarabichi, M., Lesluyes, T., Flanagan, A.M., Mertens, F., et al. (2021). Signatures of copy number alterations in human cancer.

Tarasov, A., Vilella, A.J., Cuppen, E., Nijman, I.J., and Prins, P. (2015). Sambamba: fast processing of NGS alignment formats. Bioinformatics 31, 2032–2034.

TCGA Network (2012). Comprehensive genomic characterization of squamous cell lung cancers. Nature 489, 519–525.

TCGA Network (2013). Integrated genomic characterization of endometrial carcinoma. Nature 497, 67–73.

TCGA Network (2014). Comprehensive molecular profiling of lung adenocarcinoma. Nature 511, 543–550.

Turini, S., Bergandi, L., Gazzano, E., Prato, M., and Aldieri, E. (2019). Epithelial to Mesenchymal Transition in Human Mesothelial Cells Exposed to Asbestos Fibers: Role of TGF-β as Mediator of Malignant Mesothelioma Development or Metastasis via EMT Event. Int. J. Mol. Sci. 20.

Uhrig, S., Ellermann, J., Walther, T., Burkhardt, P., Fröhlich, M., Hutter, B., Toprak, U.H., Neumann, O., Stenzinger, A., Scholl, C., et al. (2021). Accurate and efficient detection of gene fusions from RNA sequencing data. Genome Res. 31, 448–460.

Van der Auwera, G.A., and O’Connor, B.D. (2020). Genomics in the Cloud: Using Docker, GATK, and WDL in Terra (“O’Reilly Media, Inc.”).

Wala, J.A., Bandopadhayay, P., Greenwald, N.F., O’Rourke, R., Sharpe, T., Stewart, C., Schumacher, S., Li, Y., Weischenfeldt, J., Yao, X., et al. (2018). SvABA: genome-wide detection of structural variants and indels by local assembly. Genome Res. 28, 581–591.

Wang, K., Li, M., and Hakonarson, H. (2010). ANNOVAR: functional annotation of genetic variants from high-throughput sequencing data. Nucleic Acids Res. 38, e164.

Weisenberger, D.J., Siegmund, K.D., Campan, M., Young, J., Long, T.I., Faasse, M.A., Kang, G.H., Widschwendter, M., Weener, D., Buchanan, D., et al. (2006). CpG island methylator phenotype underlies sporadic microsatellite instability and is tightly associated with BRAF mutation in colorectal cancer. Nat. Genet. 38, 787–793.

Williams, M.J., Werner, B., Barnes, C.P., Graham, T.A., and Sottoriva, A. (2016). Identification of neutral tumor evolution across cancer types. Nat. Genet. 48, 238–244.

Wu, S., Turner, K.M., Nguyen, N., Raviram, R., Erb, M., Santini, J., Luebeck, J., Rajkumar, U., Diao, Y., Li, B., et al. (2019). Circular ecDNA promotes accessible chromatin and high oncogene expression. Nature 575, 699–703.

Wuilleme, S., Robillard, N., Lodé, L., Magrangeas, F., Beris, H., Harousseau, J.-L., Proffitt, J., Minvielle, S., Avet-Loiseau, H., and Intergroupe Francophone de Myélome (2005). Ploidy, as detected by fluorescence in situ hybridization, defines different subgroups in multiple myeloma. Leukemia 19, 275–278.

Zanetti, M. (2017). Chromosomal chaos silences immune surveillance. Science 355, 249–250.

Zauderer, M.G., Tsao, A.S., Dao, T., Panageas, K., Lai, W.V., Rimner, A., Rusch, V.W., Adusumilli, P.S., Ginsberg, M.S., Gomez, D., et al. (2017). A Randomized Phase II Trial of Adjuvant Galinpepimut-S, WT-1 Analogue Peptide Vaccine, After Multimodality Therapy for Patients with Malignant Pleural Mesothelioma. Clin. Cancer Res. 23, 7483–7489.

Zhang, M., Luo, J.-L., Sun, Q., Harber, J., Dawson, A.G., Nakas, A., Busacca, S., Sharkey, A.J., Waller, D., Sheaff, M.T., et al. (2021a). Clonal architecture in mesothelioma is prognostic and shapes the tumour microenvironment. Nat. Commun. 12, 1751.

Zhang, T., Joubert, P., Ansari-Pour, N., Zhao, W., Hoang, P.H., Lokanga, R., Moye, A.L., Rosenbaum, J., Gonzalez-Perez, A., Martínez-Jiménez, F., et al. (2021b). Genomic and evolutionary classification of lung cancer in never smokers. Nat. Genet. 53, 1348–1359.

Zhang, X., Zhang, W., and Cao, P. (2021c). Advances in CpG Island Methylator Phenotype Colorectal Cancer Therapies. Front. Oncol. 11, 629390.

Zheng, Y., Joyce, B.T., Liu, L., Zhang, Z., Kibbe, W.A., Zhang, W., and Hou, L. (2017). Prediction of genome-wide DNA methylation in repetitive elements. Nucleic Acids Res. 45, 8697–8711.

Zhou, L., Zheng, D., Wang, M., and Cong, Y.-S. (2009). Telomerase reverse transcriptase activates the expression of vascular endothelial growth factor independent of telomerase activity. Biochem. Biophys. Res. Commun. 386, 739–743.

