## Supplementary material for "Disentangling heterogeneity of Malignant Pleural Mesothelioma through deep integrative omics analyses": Figure S

### Supplementary Information File for Manuscript "*Disentangling heterogeneity of Malignant Pleural Mesothelioma through deep integrative omics analyses*"

September 26, 2021

Lise Mangiante, Nicolas Alcala, Alex Di Genova, Alexandra Sexton-Oates, ... , Matthieu Foll, Lynnette Fernandez-Cuesta

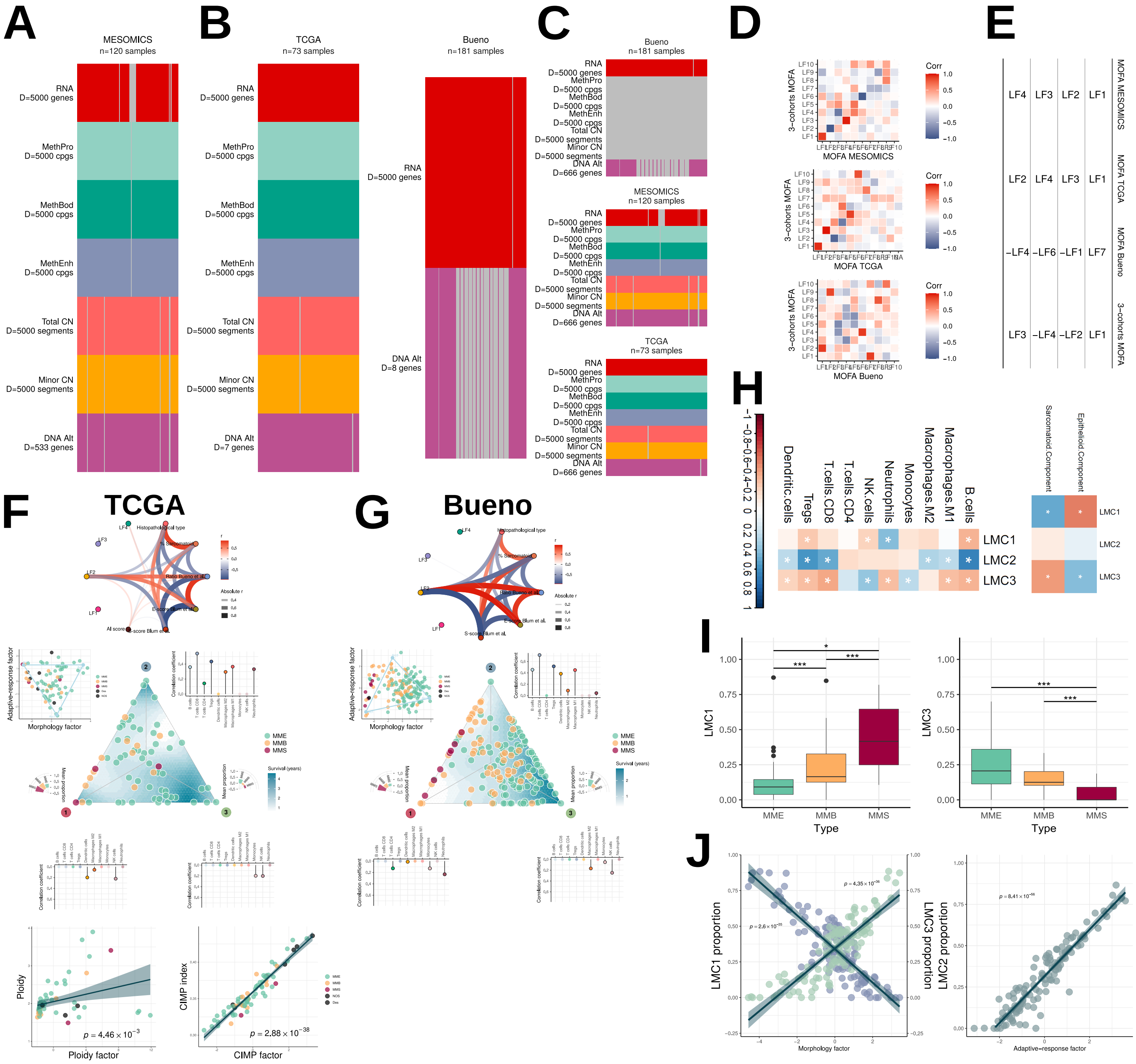

**Figure S1. MOFA of the MESOMICS, TCGA, and Bueno cohorts.** Overview of the omic data sets integrated into MOFAs, for (A) a MOFA of the MESOMICS cohort ($n = 120$), (B) separate MOFAs of the TCGA ($n = 73$) and Bueno cohorts ($n = 183$), and (C) a MOFA integrating the three cohorts (3-cohort MOFA; $n = 403$). *D* is the number of integrated omic features from genomic (rearrangements and mutations within *DNA Alt*; allele-specific copy number in *Total CN* and *Minor CN*), transcriptomic (*RNA*), and epigenomic data at promoter (*MethPro*), gene body (*MethBod*), and enhancer regions (*MethEnh*). D) Correlation between the different MOFA LFs. E) Matching between LF1, LF2, LF3, and LF4 (columns) in the different MOFAs (rows). F-G) Replication of **Figure 1B-E** with the TCGA and Bueno cohorts. H-J) Reference-free deconvolution (MeDeCom) of MESOMICS enhancer DNA methylation data. H) Correlation between proportion of each inferred latent methylation component (LMC) and the proportion of cell types within a sample (Pearson's correlation test, * = $q$-value < 0.05). I) Proportion of LMCs 1 and 3 were significantly associated with tumour type ($t$-test, * = $p$-value < 0.05, ** = $p$-value < 0.01, *** = $p$-value < 0.001). J) Correlation between LMCs 1 and 3 and MOFA's Morphology factor (left), and LMC2 and MOFA's Adaptive-response factor (Pearson correlation test).

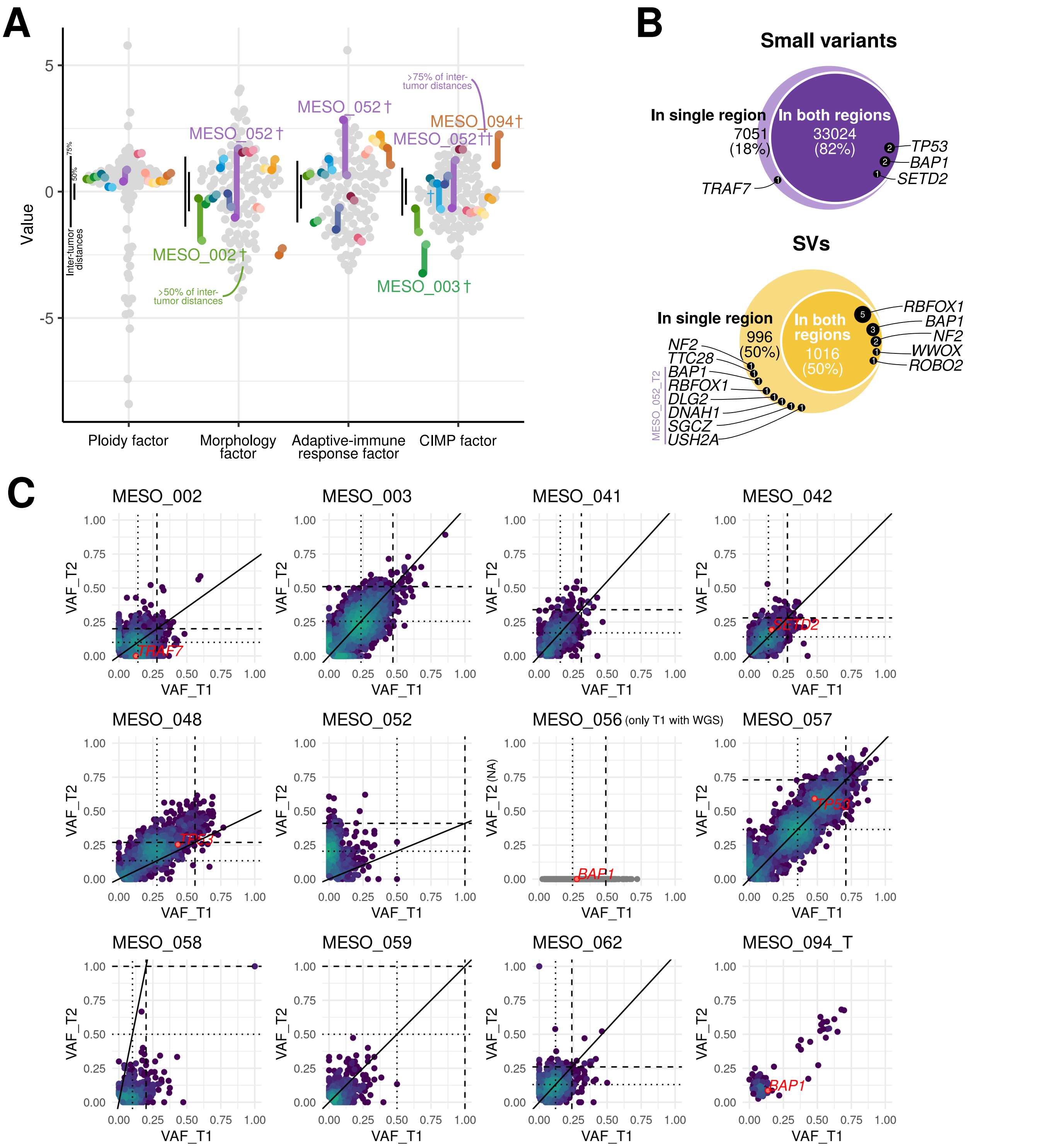

**Figure S2. Patterns of intra-tumor heterogeneity in 13 samples.** A) Position of multi-regional samples in MOFA LFs. Colors represent samples (gray: samples with a single region sequenced; colors: multi-regional samples), segments connect regions from a same tumor. Black bars on the left of each plot represent the empirical 50% and 75% percentiles of the distribution of pairwise inter-tumor distances, and tumors with an intra-tumor distance in the range [50%, 75%) and [75%, 100%) are annotated by a † and ††, respectively. Note that samples with missing genomic data are not represented on Factor 1. B) Venn-Euler diagrams of small variants (top) and SVs (bottom), indicating which alterations are found in a single or both regions. Driver mutations as highlighted in **Figure 4** are mentioned, with numbers corresponding to the number of such alterations among the 13 samples. C) Joint distribution of VAFs for the 12 multi-regional samples with WGS data (out of 13). Point colors: density measured by the number of points within a fixed radius. Damaging alterations in driver genes are represented by red points and text. Dashed lines: expected VAF for clonal alterations in diploid regions given tumor purity; dotted line: expected VAF for clonal alterations in haploid regions given tumor purity; solid line: ratio of tumor purities estimated from WGS data.

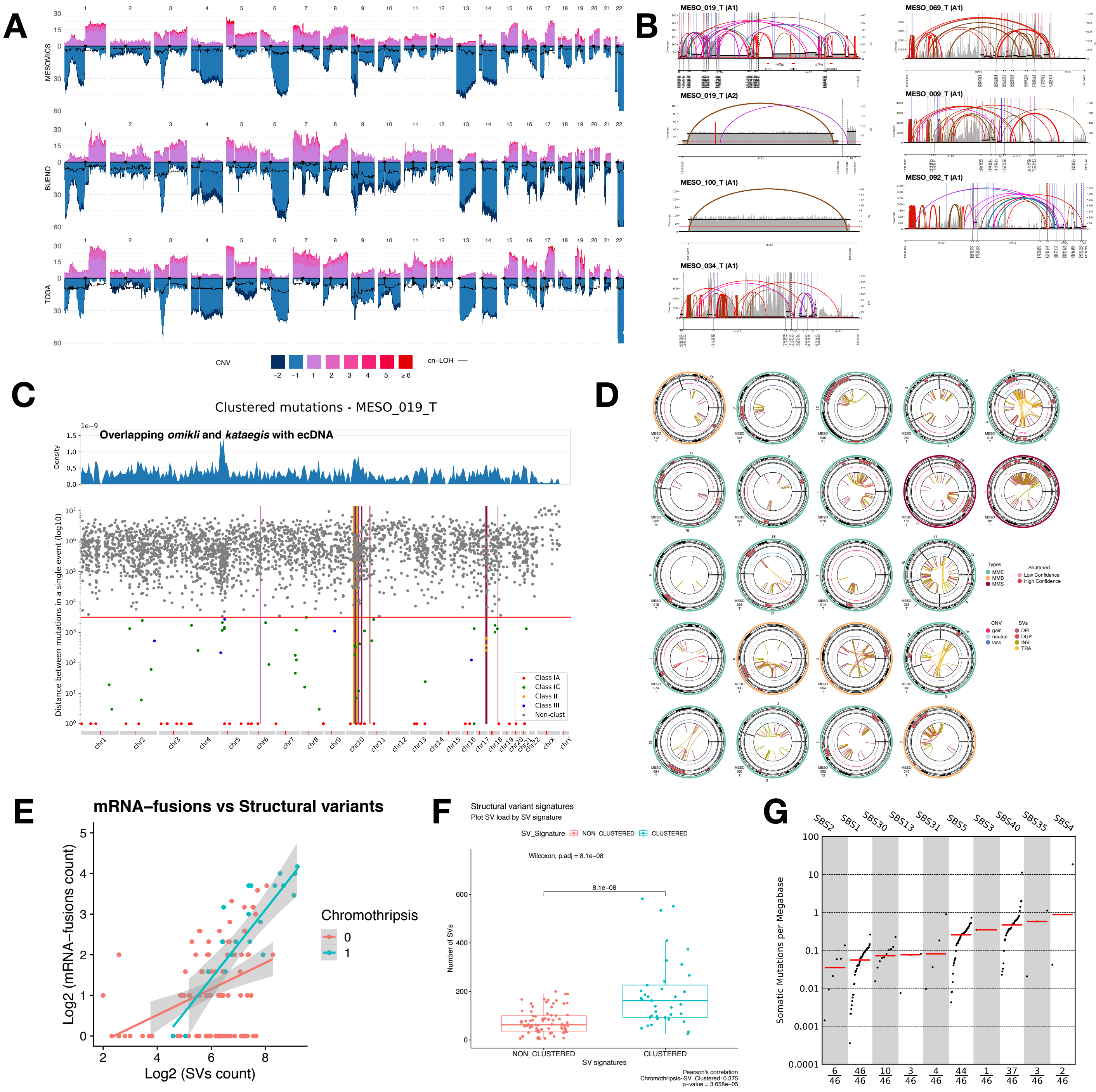

**Figure S3. Detailed genomic profiles.** A) Highly-consistent CNV profiles across MPM cohorts. B) Amplicon Architect ecDNA predictions. C) Characterization and classification of clustered point mutations of one sample with co-occurrence of kataegis and ecDNA (MESO_019_T). D) Chromothripsis analysis combining Structural Variants and Copy Number Variants. E) MPM tumors with chromothripsis tend to have more mRNA fusions per structural variant. F) Association between clustered SV signature, SV load, and chromothripsis (Pearson correlation $R = 0.38$, $p$-value=3.658e-05). G) Tumor Mutational Burden of the 10 Single Nucleotide Variant Signatures detected in the MESOMICS cohort.

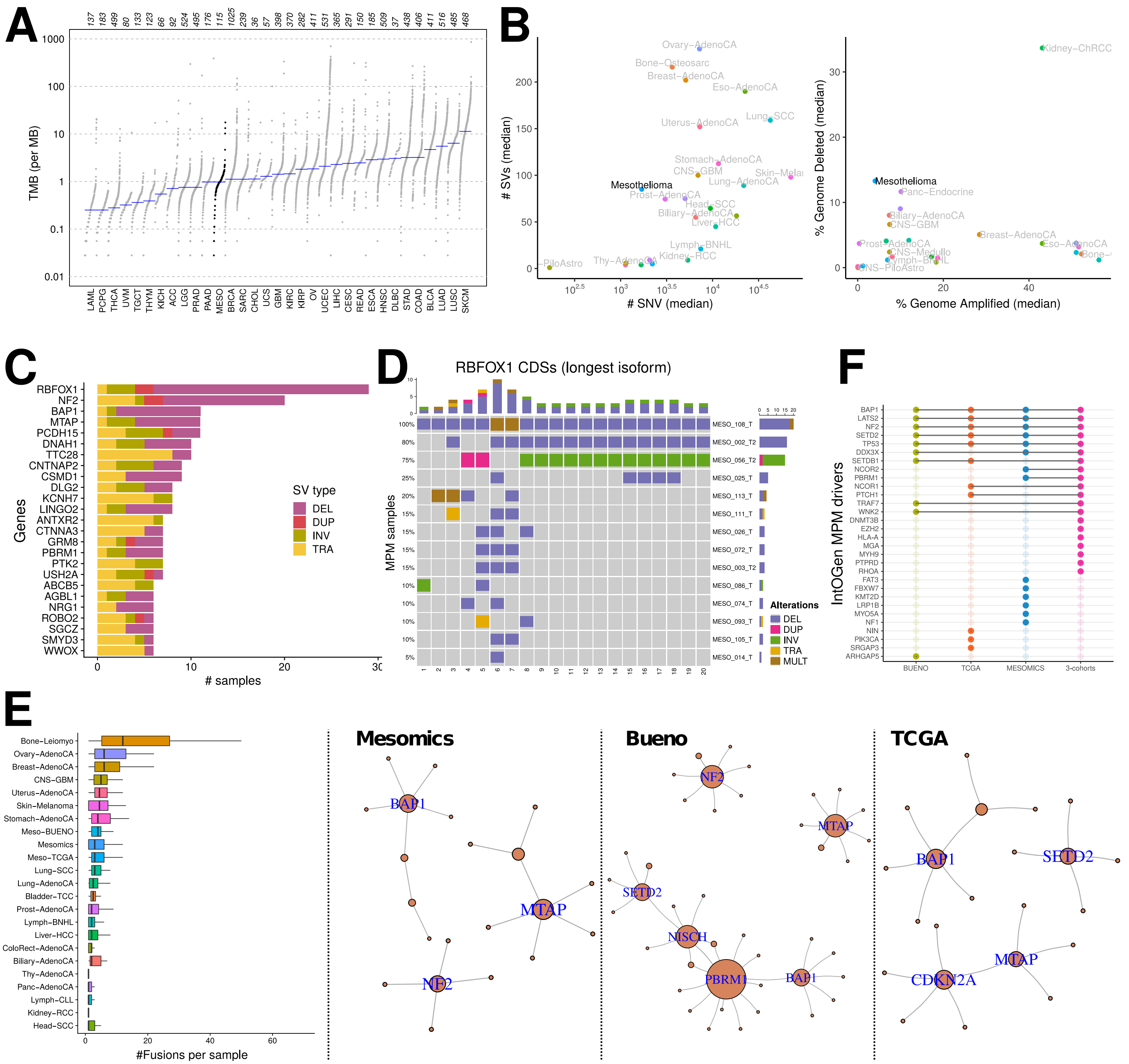

**Figure S4. MPM driver detection.** A)Tumor Mutational Burden of Mesothelioma and TCGA tumors. B) Comparison of the mutational load between mesothelioma and tumor-types from the Pan Cancer Analysis of Whole Genome data (PCAWG). Left: median number of Structural Variants (SVs) as a function of the median number of SNVs per tumor type. Right: median percentage of the genome affected by amplifications and deletions per tumor type. C) Recurrent genes affected by SVs the MESOMICS cohort. D) Structural and CNV variants affecting the coding regions of the RBFOX1 gene. E) Number of mRNA fusions per Tumor type and recurrent mRNA-fusion network for Mesomics, Bueno and TCGA cohorts. F) IntOGen MPM drivers (based on SNVs and small indels) identified within each individual MPM cohort (Bueno, TCGA, and MESOMICS) and in the pooled cohort (3-cohorts). The upset plot represents the intersections between the four sets of drivers.

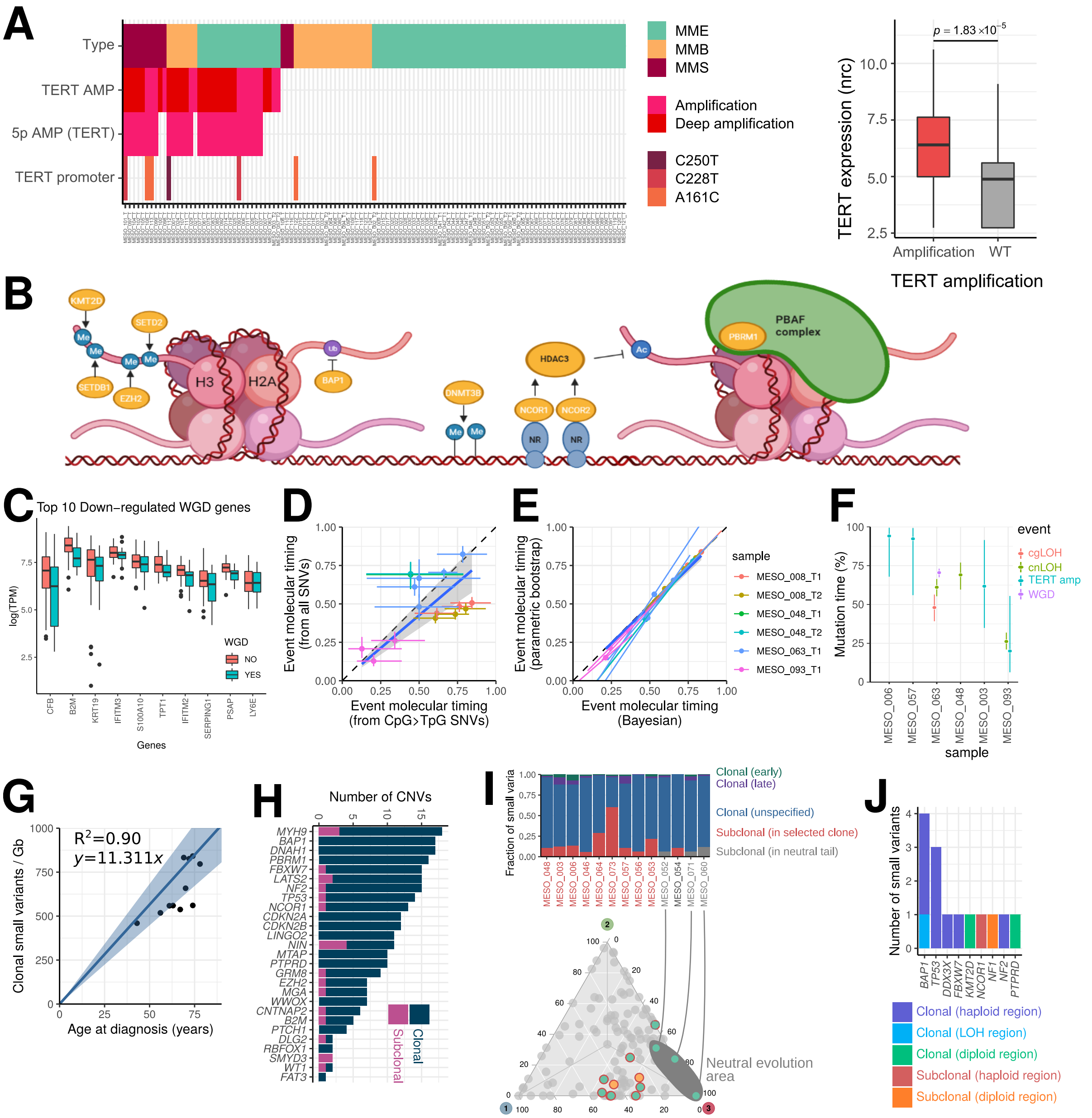

**Figure S5. Detailed impact of genomic alterations on molecular profiles of MPM.** A) Overview of TERT alterations in the MESOMICS samples. Left: oncoplot of the different types of alterations affecting *TERT* and its chromosomal arm. Right: impact of *TERT* amplification on gene expression (in normalized read counts, nrc). B) Schematic representation of mesothelioma driver ERG functions. Histone methyltransferases *KMT2D*, *SETDB1*, *EZH2* and *SETD2* methylate histone positions H3K4, H3K9, H3K27 and H3K36 respectively (Me), DNA methyltransferase *DNMT3B* maintains DNA methylation marks (Me), *BAP1* deubiquitinates H2AK119 (Ub), *NCOR1* and *NCOR2* bind to nuclear receptors (NR) as part of corepressor complexes and recruit histone deacetylase HDAC3, and *PBRM1* binds acetylated histone tails (Ac) as part of the PBAF chromatin remodelling complex (created with BioRender). C) top 10 down-regulated genes of WGD+ samples. D) Comparison of copy gain timing estimates based on CpG to TpG mutations and based on all mutations. Points represent point estimates for an event and a tumor sample, and segments 95% Bayesian CI. Samples included either underwent a WGD event (MESO_008 and MESO_063), or a large-scale LOH allowing timing (MESO_048, MESO_093). E) Comparison of copy gain timing CI using Bayesian inference or parametric bootstrapping. Points represent the centers of CI, and segments 95% CI. Samples included correspond to that presented in panel D. F) Timing of copy number gains in mutation time. Points represent tumor sample estimates and segments their 95% CI. G) Analysis of the small variant mutational burden as a function of age at diagnosis. The line denotes the mean and standard error of a linear regression model without intercept ($R^2$ and best fit equation are mentioned in the plot). H) Clonality of CNVs affecting driver genes from **Figure 4**. I) Neutral evolution detected from the VAF distribution, and corresponding proportions of small variants belonging to selected (red), neutral subclones (gray), or clonal (blue, purple, and green) (top), and position of samples in the Pareto front from **Figure 1D**. J) Clonality of driver small variant mutations; "early" indicates that the alteration predates the LOH in the corresponding region. (H)-(I) display 13 high-purity samples where clonal reconstruction was possible (see list in panel I).

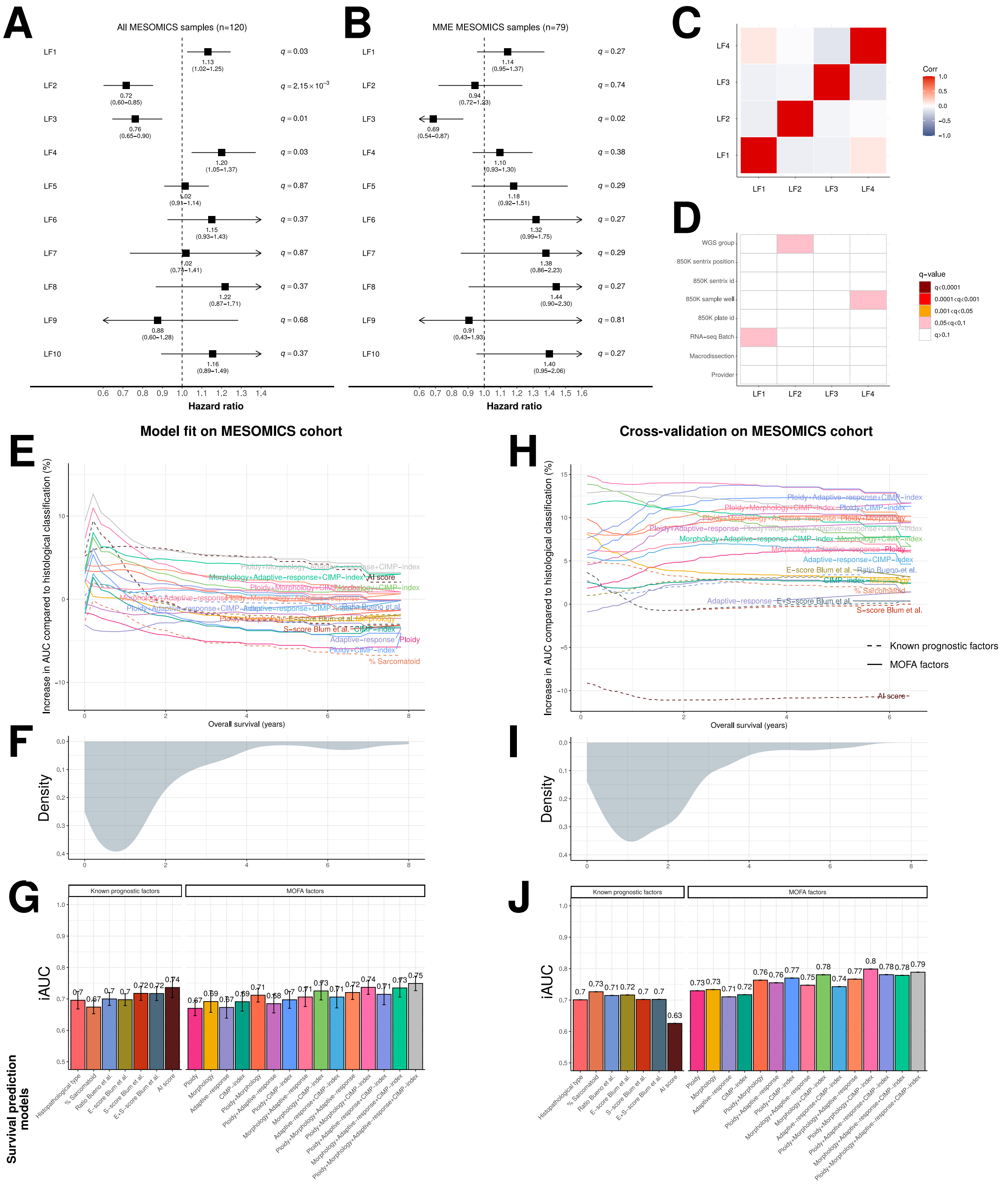

**Figure S6. Association between survival and MOFA in the MESOMICS cohort.** A) Forest plot of the survival analysis based on the ten MOFA latent factors (LFs), using a Cox proportional hazards model with LFs as continuous explanatory variables for (A) all 120 MESOMICS and (B) MME only samples ($n$=79). The black box represents estimated hazard ratios and whiskers represent the 95% confidence intervals. Wald test $p$-values are shown on the right. See corresponding data in Supplementary Table 1. C) Pearson correlation coefficients between factors. D) Linear regression test significance ($q$-value) between LFs (row) and each technical variable (column). E) Increase in AUC as a function of percentage of change compared to the AUC of model (i). F) Density of survival time within the MESOMICS cohort. G) Integral AUC (iAUC) of twenty-two Cox proportional hazards survival models based on: (i) the three histopathological types (MME, MMB, and MMS); (ii) the proportion of sarcomatoid content; (iii) the log2 ratio of CLDN15/VIM (C/V) expression proposed by Bueno and colleagues; (iv), (v) and (vi) the E-score, S-score, and combining both scores from Blum and colleagues, respectively; (vii) an AI prognostic score; (viii-xi) the one-dimensional summary of molecular data using LFs as a continuous variable; (xii-xvii), the two-dimensional summary of molecular data using either each combination of 2 LFs as continuous variables, respectively; (xviii-xxi), the three-dimensional summary of molecular data using each combination of 3 LFs as continuous variables; and (xxii), the four-dimensional summary of molecular data using all 4 LFs. Panels (E-G) present the model fit accuracy (no split between training and test sets), while (H-J) present the out-of-sample accuracy within the MESOMICS cohort (4-fold cross-validation).

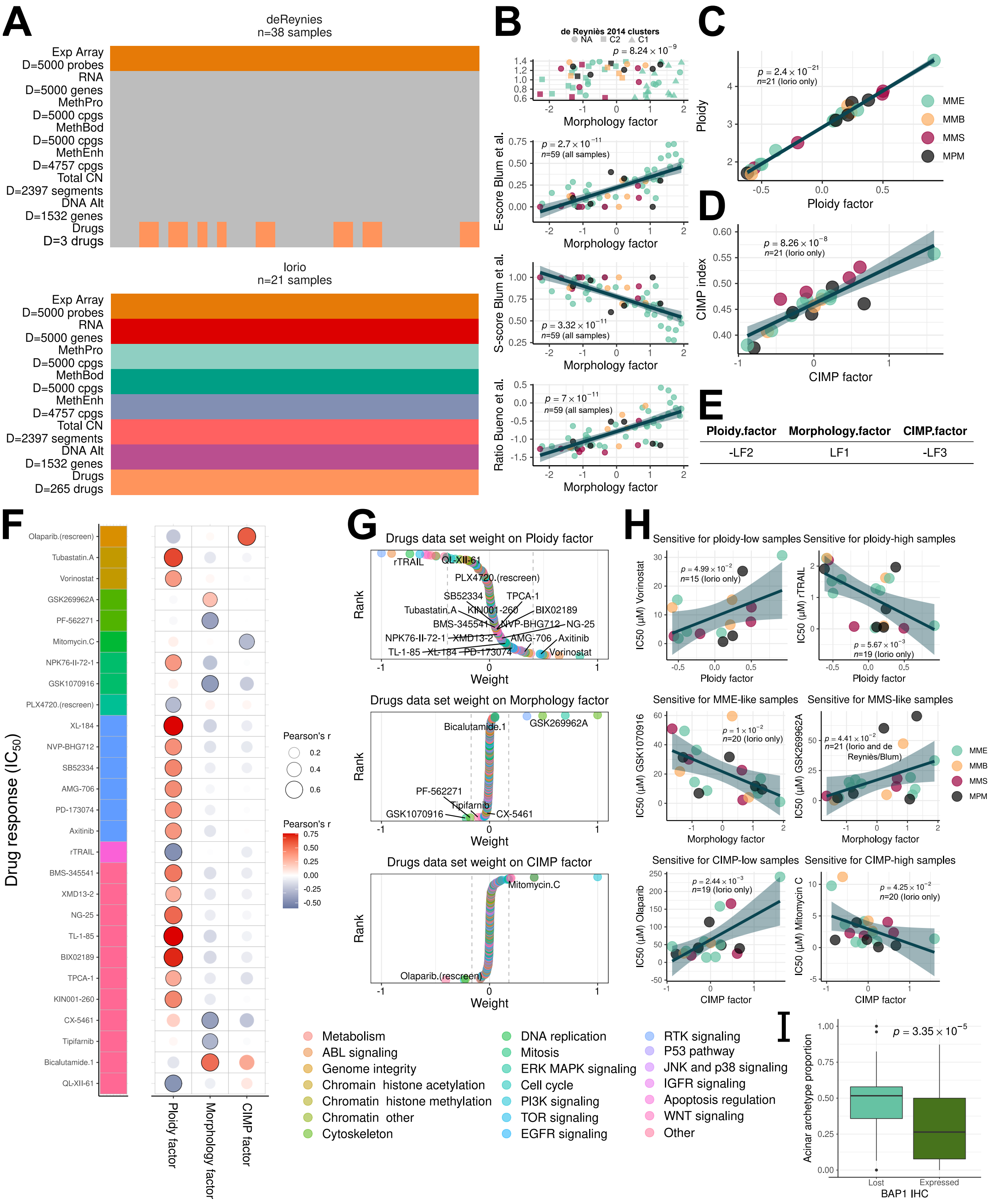

**Figure S7. MOFA of MPM cell lines.** A) Overview of the omic data sets integrated into MOFA (design follows that of **Figure S1A**). B) Association between MOFA cell lines morphological factor and the previously proposed molecular classifications. C-D) Association between Ploidy and ploidy factor, and between CIMP index and CIMP factor, respectively. E) Correspondance between MOFA cell lines LFs and the MESOMICS MOFA LFs from **Figure 1**. F) Correlations between drug responses ($IC_{50}$ in $\mu M$) and MOFA LFs of cell lines. Significant associations are annotated by black point border. G) Distribution of drug response weights from the *Drugs* data set, with drugs for which the response is significantly correlated with the given LF annotated in black. Targeted pathways are represented in (F) by a color bar (left), and in (G) by point colors. H) Correlations between representative drug responses significantly correlated with MOFA LFs from cell lines (left: negative correlations, right: positive associations). In (B) top and (I), $p$-values of an ANOVA is presented. In other (B) plots, (C), (D), (F), and (H), Pearson correlation coefficients and the associated $p$-values are displayed. I) Relationship between acinar phenotype and BAP1 expression measured by IHC. Sample sizes ($n$) and cohort of origin (Iorio or de Reyniès) are mentioned in each scatter plot.
